# Diving dinosaurs? Caveats on the use of bone compactness and pFDA for inferring lifestyle

**DOI:** 10.1101/2023.05.04.539484

**Authors:** Nathan P. Myhrvold, Paul C. Sereno, Stephanie L. Baumgart, Daniel Vidal, Frank E. Fish, Donald M. Henderson, Evan T. Saitta

**Affiliations:** Intellectual Ventures, Bellevue, Washington, United States of America; Department of Organismal Biology, University of Chicago, Chicago, Illinois, United States of America; Committee on Evolutionary Biology, University of Chicago, Chicago, Illinois, United States of America; Grupo de Biología Evolutiva, Departamento de Física Matemática y de Fluidos, Facultad de Ciencias, UNED, Madrid, Madrid, Spain; Department of Biology, West Chester University, West Chester, Pennsylvania, United States of America; Royal Tyrrell Museum of Palaeontology, Drumheller, Alberta, Canada

**Author notes:** These authors contributed equally to this work (Nathan P. Myhrvold, Paul C. Sereno). These authors also contributed equally to this work (Frank E. Fish, Donald M. Henderson, Evan T. Saitta). **Author Contributions** NPM and PCS wrote the original draft; SLB wrote individual sections; SLB, DV, NPM, and PCS collaborated on the investigation and visualization, including specimen and scan interpretations; and DV, FEF, DMH and ETS performed critical review and substantive editing of the article.

## Abstract

Measures of bone compactness in amniote tetrapods of varying lifestyle were used to infer that two spinosaurid dinosaurs (*Spinosaurus aegyptiacus*, *Baryonyx walkeri*) were diving “subaqueous foragers,” whereas a third spinosaurid (*Suchomimus tenerensis*) and other sampled nonavian dinosaurs were non-diving terrestrial feeders entering water only as waders. We outline shortcomings in this analysis that involve bone compactness sampling and measurement, lifestyle categorization, the inclusion and exclusion of taxa in the dataset, and flawed statistical methods and inferences. These many shortcomings undermine the evidence used to conclude that two spinosaurid taxa were avid divers. Bone compactness indices remain a valuable tool for interpretation of lifestyle in extinct species when based on sound dataset composition, robust statistical analysis, and consilience with evidence from functional, biomechanical, or paleoenvironmental considerations.

## Introduction

Secondarily aquatic tetrapods often exhibit greater bone density than terrestrial counterparts when examining cross sections from limb bones [1,2]. This density correlation in recent decades has been analyzed using quantitative indices based either on histologic thin sections or radiographic images [3–6]. Bone density has been used, in particular, to assess lifestyle within tetrapod clades with extinct members that have evolved secondary adaptions to an aquatic lifestyle, including amphibians (lissamphibians, temnospondyls) [5,7], reptiles (lizards, plesiosaurs, ichthyosaurs) [8–11], and eutherian mammals (talpid moles, sloths, mustelids cetaceans) [12–15]. Caution is warranted nonetheless because bone density also has been found to correlate with body mass and its skeletal manifestations as well as with stressful behaviors such as burrowing [12,14–17].

### Bone compactness in spinosaurid dinosaurs

Recently Fabbri *et al.* [18] paired a metric of bone compactness with a relatively new statistical method to investigate whether particular nonavian dinosaurs had an aquatic lifestyle. For most of the nonavian dinosaurs they examined, no association with an aquatic lifestyle was found. For the three spinosaurids, however, they reached definitive, if partially counterintuitive, conclusions. Spinosaurids were determined to be “aquatic specialists” but with “surprising ecological disparity.” *Spinosaurus* and *Baryonyx*, they argued, made regular use of “subaqueous foraging” with “fully submerged behavior,” whereas *Suchomimus*, a close relative of *Baryonyx*, was a non-diving terrestrial predator restricted to wading in the shallows [18: 852].

When Stromer first described *Spinosaurus* in 1915 from Upper Cretaceous outcrops in Egypt’s Western Desert, he highlighted the spaced, conical teeth and elongate jaws as crocodile-like adaptations for a piscivorous diet [19]. Similar inferences were made some 70 years later in descriptions of two closely related spinosaurids, *Baryonyx walker* [20] and *Suchomimus tenerensis* [21], from Lower Cretaceous outcrops in England and Niger, respectively. Although the particulars of lifestyle were not analyzed in detail in these papers, these large bipedal predators were viewed as semiaquatic, *i.e.,* hunting aquatic prey as ambush predators along the shore and while wading into shallow water.

In 2014, the notion of a semiaquatic lifestyle for *Spinosaurus* was reinforced with the description of a partial skeleton from Morocco’s Upper Cretaceous Kem Kem Group [22]. Again it was viewed as a wading shoreline predator or surface swimmer, not a diving pursuit predator. Later discovery of the high-spined tail of the Moroccan skeleton, however, inspired the “aquatic hypothesis,” which viewed the tail as an “aquatic propulsive structure” powering a “highly specialized aquatic predator that pursued and caught its prey in the water column” [23].

The central proposition that *Spinosaurus* was capable of submerged swimming in pursuit of prey drew challenges: critics pointed out the discordance in morphological comparisons to extant divers, presented paleogeographic evidence of this very large theropod dinosaur in inland habitats, and detailed biomechanical calculations that show both that *Spinosaurus* would have been unstable when floating in water and too buoyant to dive, and that available propulsive force from its hind limbs or tail was insufficient for pursuit predation [24–26].

In response, many of the authors of the “aquatic hypothesis” turned to bone compactness as an additional means to assess lifestyle. Fabbri *et al.* ([18]; “Fabbri *et al.*” below) used bone cross sections and an index of bone density, global bone compactness (*Cg*), to argue that *Spinosaurus* and *Baryonyx* were “fully submerged” subaqueous foragers but that the African spinosaurid *Suchomimus* was a terrestrial non-diver that at best waded into shallow waters.

### Categorization caveats

Fabbri *et al.* assembled datasets of exemplar taxa (“training datasets”) using two categorical variables for functional capability, “subaqueous foraging” and “flying,” with three potential values (*0–2*) for range of presence: s*ubaqueous foraging*: unable (*0*), able but infrequent (*1*), frequent (*2*); *flying*: unable (*0*), non-sustained flight (*1*), sustained flight (*2*). If behaviors of an extinct species were regarded as uncertain, it was categorized as “unknown.”

“Subaqueous foraging” was never defined in the original paper [18] and was supplanted by “diving” in a table of behavioral scores [18: Suppl. information]. “Subaqueous,” etymologically, means complete submergence and does seem equivalent to “diving.” “Foraging,” by definition, means searching for food, be it plant or animal. A “subaqueous forager” is thus either a habitually diving predator in pursuit of underwater prey, such as a sperm whale, or a habitually diving herbivore that feeds on underwater plant resources, such as a manatee.

Yet when challenged as to why they classified as “subaqueous foragers” hippos and tapirs, which do not forage appreciably underwater [27], the authors responded that what they really meant in using the term “subaqueous foraging” was habitual “subaqueous submersion” [28], whether for foraging or simply for concealment. “Habitual diving” thus seems to capture the behavior of interest, which then brings into question whether their subgroup of “subaqueous foragers” is adequate to evaluate spinosaurids as underwater pursuers of prey.

Fabbri *et al.* represented each sampled taxon with a two-dimensional data point (*Cg*, log_10_(*MD*)), where *Cg* is their chosen index for bone compactness calculated by the Bone Profiler program [3] and *MD* is the maximum diameter of a sampled bone. Two datasets were assembled from bone-shaft cross sections, one based on femora and the other on dorsal ribs. Most taxa are represented by one compactness measurement from a single thin section or from a radiograph of the shaft of one bone. A phylogenetic consensus tree was used to control for phylogenetic bias.

For clarity and concision, we abbreviate the functional groups identified by Fabbri *et al.*, using *F* and *D* to designate “flying” and “diving,” respectively, for the two lifestyle behaviors they identified. Each lifestyle was categorized using one of three potential values (*0–2*): “absent” (*0*), “present but infrequent” (*1*), and “frequent” (*2*) (Table 1). Here we abbreviate each taxon score as *FxDy*, where *x* and *y* denote the flying and diving variables, respectively. Habitual (frequent) divers, or “subaqueous foragers” in the terminology of Fabbri *et al.*, is a category they allied with the extinct spinosaurids *Spinosaurus* and *Baryonyx* (Table 1: *F0D2*, subgroup 6). *Suchomimus*, by comparison, was considered an extinct nonflying/nondiving (presumably “terrestrial”) nonavian dinosaur (Table 1: *F0D0,* subgroup 3). The razorbill *Alca torda*, as another example, is an extant seabird that frequently flies and dives (Table 1: *F2D2*, subgroup 8). In sum, there are two functions, three variables, and three ways to subdivide members as to whether they are living, extinct, or a mix of the two for a total of 18 potential subgroups (Table 1: 9 subgroups shown).

**Table 1.** Example functional groups and subgroups in Fabbri *et al.* [18] as designated in this paper.

| Our abbreviation | Lifestyle | Subgroup number | Taxon type | Subgroups |
| --- | --- | --- | --- | --- |
| *F0D0* | Terrestrial<br>(nonflying/nondiving) | 1 | All | Terrestrial |
|  |  | 2 | Extant | Extant terrestrial |
|  |  | 3 | Extinct | Extinct terrestrial |
| *F0D2* | Nonflying diver<br>(nonflying/frequent diving) | 4 | All | Divers |
|  |  | 5 | Extant | Extant divers |
|  |  | 6 | Extinct | Extinct divers |
| *F2D2* | Flying diver<br>(frequent flying & diving) | 7 | All | Flying divers |
|  |  | 8 | Extant | Extant flying divers |
|  |  | 9 | Extinct | Extinct flying divers |

Fabbri *et al.* [18] assigned for “flying” the values 0 = “unable,” 1 = “non-sustained flight,” and 2 = “sustained flight.” They assigned for “diving” the variables 0 = “unable” to dive, 1 = “infrequent” diving, and 2 = “frequent” diving. Some taxa were designated “unknown” rather than assigned a variable. The three most important subgroups are shown in the table.

### Full and culled datasets

Fabbri *et al.* used four datasets in their analysis, which are termed as “training” datasets for their classification method (Table 2). The first two datasets included all sampled taxa for femur data and ribs (labeled here ds1, ds2), whereas the last two datasets (labeled here ds3, ds4) purged taxa regarded as “graviportal” or “pelagic.” The taxa in these four datasets were categorized as described above according to their “flying” and “diving” capacities.

**Table 2.**
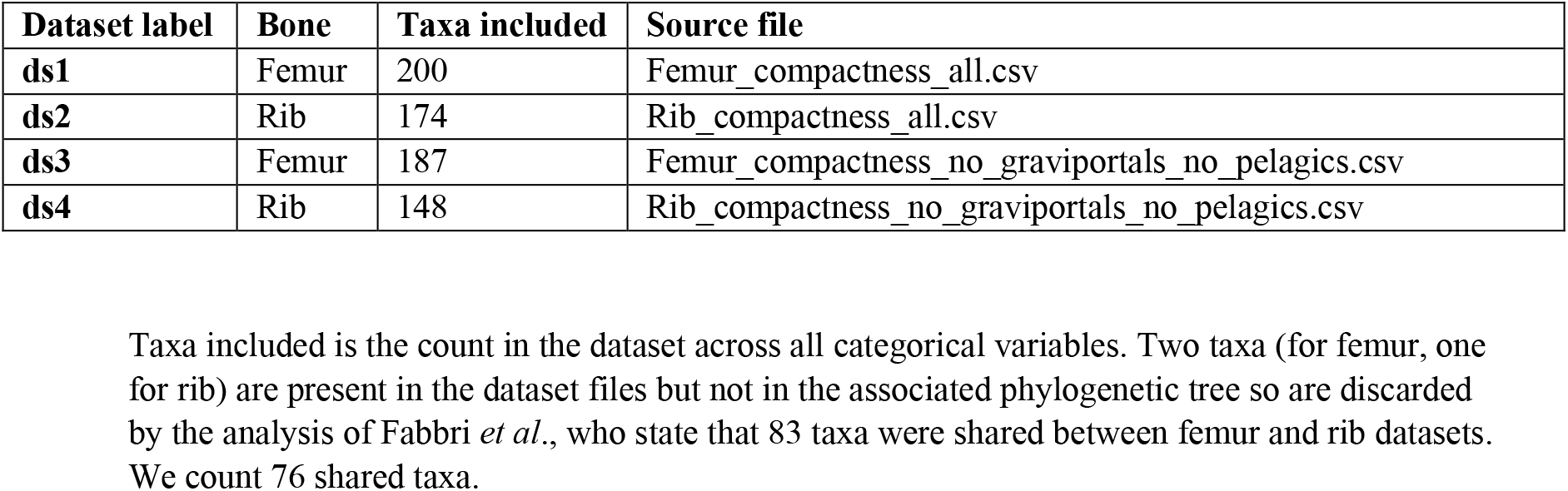
Four training datasets used by Fabbri *et al.* [18: Suppl. information].

| Dataset label | Bone | Taxa included | Source file |
| --- | --- | --- | --- |
| ds1 | Femur | 200 | Femur_compactness_all.csv |
| ds2 | Rib | 174 | Rib_compactness_all.csv |
| ds3 | Femur | 187 | Femur_compactness_no_graviportals_no_pelagics.csv |
| ds4 | Rib | 148 | Rib_compactness_no_graviportals_no_pelagics.csv |

Taxa included is the count in the dataset across all categorical variables. Two taxa (for femur, one for rib) are present in the dataset files but not in the associated phylogenetic tree so are discarded by the analysis of Fabbri *et al.*, who state that 83 taxa were shared between femur and rib datasets. We count 76 shared taxa.

We discuss below several issues regarding the composition of these datasets to adequately assess the meaning of bone density in spinosaurids.

### PGLS and pFDA analyses

Fabbri *et al.* first performed phylogenetic generalized least squares (PGLS) analyses that showed weak, but statistically significant, correlations between *Cg* and *D = 2* (“frequent diving”). Correlations between *Cg* and values for *F* (“flying”) were weak or not considered statistically significant. Their second tier of analyses use the relatively new phylogenetic flexible discriminant analysis (pFDA) algorithm [29,30] to process training datasets and then classify other data points that include spinosaurids and other taxa by statistical properties “learned” from the training dataset.

In this paper we reexamine the analysis of Fabbri *et al*., not only to evaluate the viability of their central conclusion regarding the lifestyle of three spinosaurids, but also as a case study to consider the relevance of variability in bone sampling, variation in the calculation of bone compactness indices, and the assumptions underlying the use of statistical analyses of bone compactness across living and extinct taxa. To that end, we take a close look at the extant taxa labeled “subaqueous foragers” by Fabbri *et al.* to determine the validity of that categorization. Then we critically examine the composition of the training datasets and the assumptions required for use of the pFDA algorithm.

Like many statistical methods, pFDA is based on assumptions about the statistical distribution of the data. In particular, pFDA requires multivariate normal distributions [29,30]. We applied multiple statistical tests to key datasets in Fabbri *et al.* but find that they fail to meet this prerequisite. Indeed, some of the datasets are statistically indistinguishable from a uniform random distribution of points in two dimensions. These findings question the validity of pFDA results for these data.

Because pFDA does not automatically generate a *p* value, confidence interval, or other quantitative metrics for random error, it is not rigorous as a statistical hypothesis test or as an estimation procedure. We performed supplementary analyses to generate such estimates. No prior analysis of pFDA has examined uncertainty due to the finite sizes of training datasets. We show that random effects and sample-size uncertainties appear to significantly undermine confidence in the results of Fabbri *et al*.

## Materials and methods

### Institutional Abbreviations

BSPG Bayerische Staatssammlung für Paläontologie und Geologie, Munich, Germany.

FMNH Field Museum of Natural History, Chicago, Illinois, United States of America.

FSAC Faculté des Sciences Aïn Chock, University of Casablanca, Morocco.

MNBH Musée National Boubou Hama, Niamey, République de Niger.

CMN Canadian Museum Nature, Ottawa, Canada.

UCRC University of Chicago Research Collection, Chicago, United States of America.

### Computed tomography

Computed-tomographic (CT) scans were generated for femora of *Suchomimus tenerensis* (MNBH GAD500, MNBH GAD72) and *Spinosaurus aegyptiacus* (FSAC-KK 11888) at the University of Chicago Hospitals by Dr. Nicholas Gruszauskas and Dr. David Klein using a Philips Brilliance iCT 256-slice multi-detector CT scanner. CT scans for the *Spinosaurus sp*. femora (CMN 41869, CMN 50382) were generated by Vincent Bolduc at the Transportation Safety Board of Canada’s North Star Imaging CT scanner. Scan settings for each of the specimens are included in S1 Table.

### Bone compactness measurement

We used Materialise Mimics Innovation Suite 23.0 to segment CT scans of specimens new to this study. We positioned long bones for cross section perpendicular to the shaft axis. We used a threshold that highlighted bone, exporting that image of the cross-sectional slice. To ensure pixels were correctly read by Bone Profiler, Affinity Photo was used to binarize the image.

We used the BoneProfileR R package [3] and the binarized femoral slice images provided by Fabbri *et al.* in their Fig 1 and Extended Data Figs 1–5 [18] to generate additional BoneProfileR variables along with *Cg* and to determine how they varied. Because user-input parameters for the program were not reported in Fabbri *et al.*, some variance in our results is likely. For complete sections, we used the ontogenetic center (recommended by BoneProfileR authors) in the BP_EstimateCompactness function and defaults of 60 angles and 100 distances. We collected bone compactness data from the flexit and flexit with pi rotation models. There were three partial cross sections, which were run using a user-defined center with setting partial = TRUE in the BP_EstimateCompactness function. A few of the cross sections in Fabbri *et al.* are low-resolution and necessitated re-binarization.

**Fig 1.**
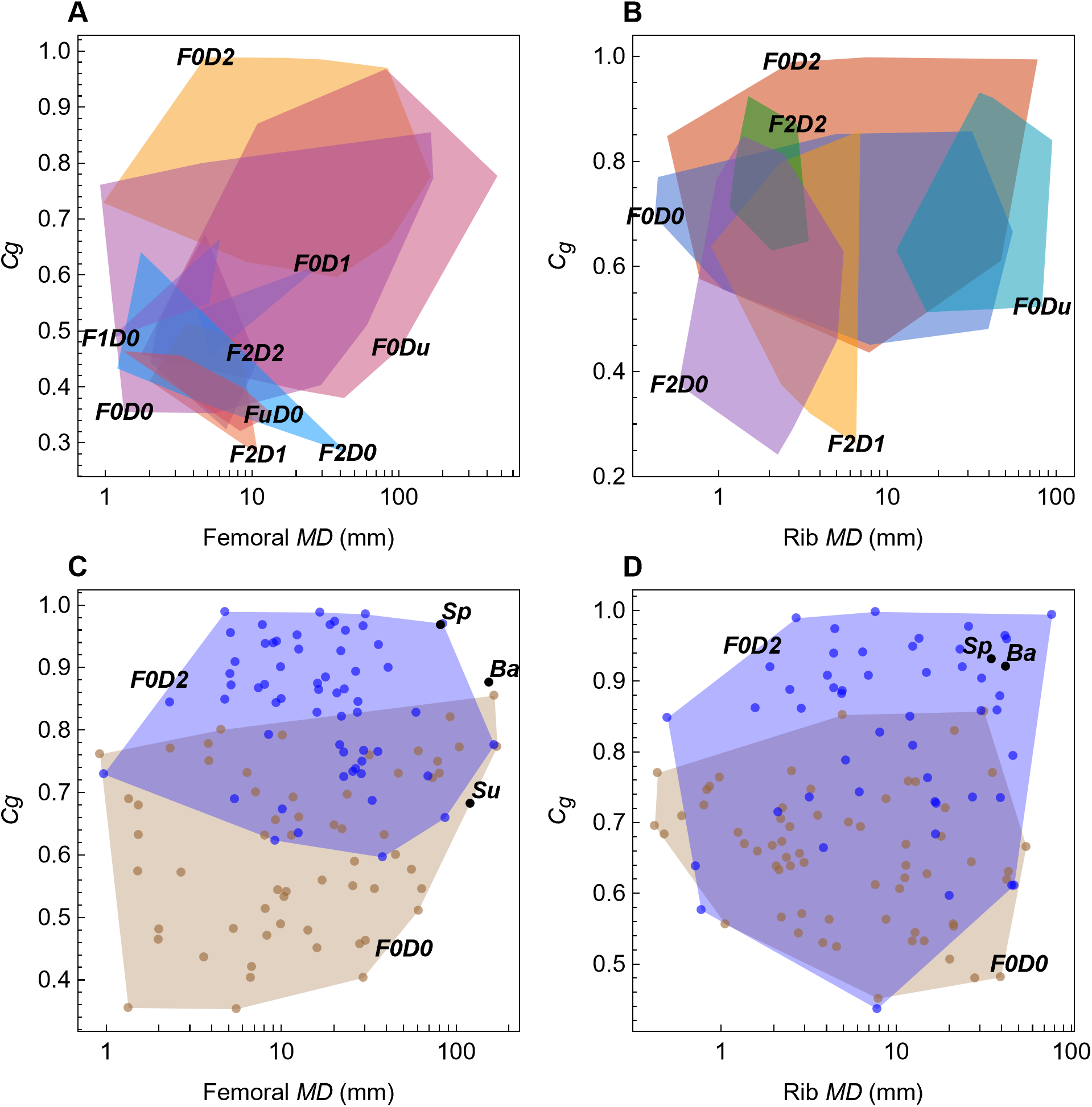
Lifestyle overlap in femoral and rib data from Fabbri *et al.* [18]. Femoral (A, C) and rib (B, D) plots of bone diameter versus bone compactness (*Cg*). (A, B) Convex hull polygons colored by functional group, as defined by Fabbri *et al*. (C, D) Points and corresponding convex hull polygons for terrestrial (*F0D0*) and nonflying/diving (*F0D2*) groups. Abbreviations: *0*, absent; *1*, rarer; *2*, habitual; *Ba*, *Baryonyx*; *D*, diving; *F*, flying; *Sp*, *Spinosaurus*; *Su*, *Suchomimus*; *u*, unknown.

Useful metrics, besides *Cg*, include *P*, *S*, and *TRC* [3,31]; we recorded the min, median, max of *P*, *S*, and *TRC* for each specimen. For standardization of the dataset, we *z*-scored all of these variables as well as midshaft diameters provided by Fabbri *et al.* [18: Suppl. information].

#### Statistical methods

All pFDA results in this paper were based on R scripts and associated data files obtained from the authors of Fabbri *et al.* [18] and on base-level pFDA code deposited by Motani and Schmitz in an online repository [32]. Bootstrap trials and related modifications were done in R, with minimal changes necessary to the base-level pFDA code for debugging. Bootstrapping pFDA requires randomly selecting with replacement a sample of the dataset taxa of the same length as the original dataset. Phylogenetic trees must be pruned appropriately, which was accomplished in the same manner as pFDA using the same R library functions. The data-gathering aspect of the BCa bootstrap algorithm, which is based on both bootstrap and jackknife trials, was also implemented in R.

Permutation tests on the rank distribution of *Cg* between extinct and extant taxa were implemented in Mathematica using standard methods [33]. Statistical analysis of the output of the trials gathered in R, along with the data tables and figures, were generated with code written in Mathematica v13.2 [34]. Statistical tests such as Brown-Forsythe, Conover, and Levene variance equivalence tests used standard library functions in Mathematica. Other library functions were used for distribution fitting in the construction of smooth kernel distribution plots and quantile-quantile plots.

Code was written by the authors for simple LDA and a Monte Carlo simulation using LDA. Confidence intervals were calculated in code written in Mathematica that processed data output from R scripts. Equation (3) was obtained by symbolic mathematical derivation in Mathematica.

Code implementing the Hopkins statistics tests was implemented in Mathematica, using the published algorithms [35,36]. Under the null hypothesis, the Hopkins statistic is expected to approximate a beta distribution Beta (*m*, *m*), where *m* is the number of points sampled. As recommend in the literature, a random sample of 20% of the points in a test set was used. As an additional verification, a Monte Carlo suite of 1000 pseudorandom examples of a uniformly random distribution were generated and tested to build an empirical sampling distribution for the null hypothesis. This was done separately for each of the variants of the Hopkins statistic test, as well as for each point count in a set being tested.

## Results

### Bone density interpretation

#### Bone density and aquatic function

Increased bone density as a secondary adaptation to an aquatic environment occurs by pachyostosis, which involves an increase in dense peripheral bone deposits, and/or by osteosclerosis, which involves an increase in bone deposition toward the center of the medullary cavity of long bones. The potential advantages for semiaquatic and fully aquatic tetrapods are well summarized by Houssaye [37]:

> Nopcsa (1923a) suggested that pachyostosis s.l. [*sensu lato*] might cause an increase in weight of the thoracic region in animals with large lungs, to correct the obliquity of their body trim to facilitate swimming and breathing. Kaiser (1960) considered that the aim is to counterpoise the volume of air present in the lungs to facilitate diving. Wall (1983) added the idea of a counteraction against waves to improve stability in agitated waters. More recently, Taylor (2000) suggested that increasing skeletal mass might be useful not so much to neutralize the buoyancy of existing lungs, but to allow their enlargement and thereby provide a larger oxygen store. Even if bone ballast reduces acceleration abilities and manoeuvrability, this cost in locomotion is probably less important in slow-swimming or bottom-walking animals. Taylor (2000) considered that pachyostosis s.l. allows these animals to lose buoyancy quickly with depth and, therefore, to reach neutral buoyancy at shallower depths, which means that they can float stationarily or hover at shallow depth, without expending energy to maintain their horizontal position; or (ii) walk on the bottom in shallow waters, while searching for slowly-moving or stationary food. Moreover, the increased oxygen store would allow them an extension of dive duration. This is typically the case for sirenians which, during their dives, can adjust their position with minor corrective movements of the forelimbs and tail fluke. Therefore, this increase in skeletal mass would have more advantages than disadvantages in forms that cannot swim very fast or dive very deep (Taylor 2000).

A key point in the passage is that the biomechanical needs of buoyancy for herbivores can be quite distinct from active predators. Predatory species chasing fast or deep aquatic prey, in fact, experience tradeoffs that can disfavor increased bone density, as summarized by Houssaye [37]:

> Whereas pachyostosis s.l. is observed in early mosasauroids, nothosaurs and Archaeoceti, it is absent in fast-swimming taxa. Therefore, it appears that pachyostosis s.l. is lost in a lineage in taxa adapting to open sea and feeding on mobile preys, and, therefore, needing good manoeuvrability, diving and accelerating abilities. Indeed, the new environmental constraints would favor bone lightening and lung reduction (as deep diving calls for storage of oxygen in tissues rather than in lungs to minimize problems of decompression) in these fast swimming taxa that use a hydrodynamic buoyancy control strategy (Taylor 2000).

Decreased bone density has been documented in many fully aquatic, deep-diving taxa [10,11,38–40]. In a PGLS model, Sun *et al.* [41] found a statistically significant correlation between diving depth and *reduced* bone density, including reduced *Cg*, which has been corroborated in studies of cetaceans and other taxa [42].

The relationship between *Cg* and semiaquatic or fully aquatic taxa is thus not simple [6,7,11,16], as some taxa exhibit increased and others decreased *Cg*. The fully aquatic sirenians have very dense bones, as noted above, because it suits minimal energy expenditure while foraging for stationary underwater vegetation, whereas the fully aquatic orca has lower *Cg* for fast pursuit of prey. The *D* = *2* (“diving”) datasets in Fabbri *et al.*, however, include herbivorous underwater grazers like the manatee, which are of questionable comparative value for spinosaurid predators hypothesized to be in pursuit of subaqueous prey. In sum, increased bone density per se does not characterize all fully aquatic or deep-diving pursuit predators nor correlate with average or maximum diving depth.

### Bone density and body size

Large-bodied terrestrial taxa often have increased *Cg* [7,11,16,43– 48], and *Spinosaurus* is in the top tier of body mass among theropods. Other lifestyles, namely arboreal [13] and fossorial (burrowing) taxa [12] have been associated with increased *Cg*, but these are not relevant for spinosaurids. The potential for increased *Cg* as a consequence of large body size, however, must be considered as a viable alternative hypothesis, especially when using cross-sectional data from the femur of a biped. The effect of large body size has been well considered in the common hippo [11]:

> However, it is difficult to determine whether the pattern observed in Hippopotamus reflects its graviportal limbs or the benefit of a slight increase in bone mass in its legs enabling their use as ballast and offering stability in water. As a result, both adaptations might be mistaken, or even synergistic, and it seems almost pointless to try to unravel their evolutionary integration. Adaptation to a graviportal limb morphology should thus be taken into consideration when analyzing possibly amphibious taxa displaying a terrestrial-like morphology, and thus notably in the study of the early stages of adaptation to an aquatic life in amniotes.

The dataset includes very few large nonavian dinosaurs comparable to spinosaurids, many of which have infilled medullary spaces in limb bones [5,6]. Large-bodied ornithischians are represented only by *Stegosaurus*, and *Alamosaurus* is the only full-sized sauropod in the dataset. Both of these genera as well as the African elephant *Loxodonta* have femoral compactness comparable to *Baryonyx*. Increased *Cg* in *Spinosaurus*, hence, may be a secondary semiaquatic adaptation, a consequence of its large body size, or conceivably a combination of the two.

### Lifestyle interpretation

#### Lifestyles of extant “subaqueous foragers.”

Datasets in Fabbri *et al.* include taxa in the “subaqueous forager” category that do not forage underwater. This includes the common hippo (*Hippopotamus amphibious*), pygmy hippo (*Choeropsis liberiensis*) [49], common tapir (*Tapirus terrestris*) [50], Malayan tapir (*Tapirus indicus*) [51], beaver (*Castor fiber*) [52], and European water vole (*Arvicola amphibius*) [53]. Although each of these taxa has secondary semi-aquatic adaptations to aquatic habitats, they nevertheless forage substantially—in some cases exclusively—on land and above water. These species habitually enter aquatic habitats to avoid predators, not to forage.

The Fabbri *et al.* training sets include the American mink (*Neovision vision*) [54] and Pyranean desman (*Galemys pyrenaicus*) [55] that forage for the majority of their diet above water. Both are assigned *D = 2* and therefore misclassified as “subaqueous foragers” in femoral and rib datasets. Similarly, the American alligator (*Alligator mississippiensis*) [56,57] and the Nile crocodile (*Crocodylus niloticus*) [58] are misclassified as subaqueous foragers despite ample evidence that both have a largely terrestrial diet as adults, though frequently concealing themselves partially or fully submerged while stalking animals on the shore [56,59–64]. These species also document the complexity of functional assignment, as they grow by orders of magnitude and often exhibit dietary change during ontogeny [62,65,66]. Crocodilians may be insectivores while very small, submerged piscivores at moderate size, and transition to terrestrial prey as adults. A scheme that does not specify ontogenetic stage cannot classify such species. This issue is highly relevant because ontogenetic dietary niche partitioning has also been identified in theropod dinosaurs [67–69]. Like crocodilians, some predatory dinosaurs spanned a similar range of body size from hatchling to adult and almost certainly accessed a range of size-appropriate prey. Spinosaurids, which include one of the largest known theropod dinosaurs, are likely to have sought a sequence of preferred ecological niches during ontogeny.

Many other terrestrial species forage by capturing aquatic prey just under the water surface without being submerged themselves. Terrestrial brown bears, black bears, and wolves (not included in Fabbri *et al.* training datasets) prey upon spawning salmon [70–73], often plucking them from below the surface. Jaguars hunt caiman or capybara both above and below water [74–76]. In each case, the prey being foraged is underwater, but the predator doing the foraging is either partly or wholly above the water. Occasional full submersion would never qualify them as secondarily aquatic or even semi-aquatic species, even when aquatic prey comprise a substantial, or even critical, component of their diet [77].

Eagles, osprey and other raptors, and many other birds such as skimmers and egrets similarly forage while flying by grabbing fish from under the water surface [78–81]. The foraging is clearly subaqueous, but it is equally clear that the airborne forager is almost entirely above the water. Birds such as herons, storks, egrets, and cranes stand in shallow water with most of their body above water, plunging their head underwater to capture fish and other aquatic prey [82–84]. That behavior, which literally would qualify as “subaqueous” foraging, does not involve diving.

The commonality across these cases is that the prey item is underwater, so the foraging by necessity occurs there, but the forager does not fully submerse its body. As a result, there is no connection between foraging behavior and diving frequency, and thus no plausible direct connection with bone density. Animals that forage for prey underwater may or may not be underwater themselves and thus may not need the ballast effect of higher bone density. Full representation of “subaqueous foragers” would necessitate including these taxa in the *D = 2* category.

The *D = 2* dataset in Fabbri *et al.* appears to have been composed of animals that dive, rather than those that forage underwater. The pivot between one concept to the other occurs in this passage [18]:

> Previous studies applied different categorizations for the characterization of aquatic lifestyles among extant and extinct taxa: ‘aquatic’ and ‘semiaquatic’ were used contra ‘subaqueous foraging’ applied in this study. Our ecomorphological attribution is focused on a specific behaviour linked to an ecology, rather than a categorization of its entirety. We find our categorization to be more accurate: for example, previous studies coded penguins and cetaceans as aquatic, while crocodilians were stated as semiaquatic. Whereas penguins and crocodilians are still ecologically dependent on terrestrial environments (for example, for laying eggs), cetaceans are completely independent from land. On the other hand, all these clades engage in subaqueous foraging. Therefore, our ecological attribution is in agreement with previously applied ecological categories, but do not exclude dependency to terrestrial environments to satisfy autecological requirements, such as reproductive behaviour.

Essentially no supporting evidence is presented to support the utility or “accuracy” of substituting “subaqueous foraging” in place of the more traditional characterizations “semi-aquatic” and “aquatic.” The examples we have cited above of animals that dive but do not forage underwater and those that forage underwater without diving show this proposition to be false. Again, it seems that Fabbri *et al.* use “subaqueous foraging” to categorize “subaqueous” activity or habitual diving.

The novel claim of Fabbri *et al.* is to demonstrate correlation between *Cg* and “underwater foraging” or habitual diving. Absent such a correlation, their analysis would conclude that spinosaurids had some degree of semi-aquatic adaptation, which has long been proposed.

#### Lifestyles of extinct “subaqueous foragers.”

Training datasets usually are restricted to extant taxa with observable lifestyles when the aim is to discern lifestyle in extinct species. The lifestyle for many extinct species, in contrast, is specified in the training datasets of Fabbri *et al*. The categorical variable *D* (“diving”) is scored in many extinct species as nondiving or as rarely or frequently diving “subaqueous foragers” (*0–2*). For taxa with flippers, such as *Plesiosaurus*, this interpretation is a reasonable extrapolation based on morphology and paleoenvironment of fossilization. For others, such as the extinct hippopotamus *Hexaprotodon*, there would be considerable uncertainty regarding its habits in water, as it has fewer secondary aquatic adaptations than the living common hippo [17], which forages in terrestrial environments.

### Lifestyles of extinct “nondivers.”

Similar extrapolation regarding the lifestyle of extinct species was not granted to those long interpreted as fully terrestrial. A large subset of taxa, 37 nonavian dinosaurs, are scored as nonflying reptiles with “unknown” diving capacity (*F = 0, D = unknown*). All nonavian dinosaurs in the analysis (including *Spinosaurus*), could thus potentially have been rare or frequent “divers.” That subset includes *Stegosaurus* with elephantine feet, *Oviraptor*, which is known to have lived and nested in xeric habitats far from any shoreline, and *Alamosaurus*, with columnar limbs discovered in inland terrestrial deposits. It seems arbitrary to be able to score in advance nearly all “subaqueous foragers,” while remaining blind to the habits of non-spinosaurid nonavian dinosaurs, all of which have long been regarded as fully terrestrial [5]. The categorization of these taxa as “unknown” for diving is a major reason that the terrestrial cohort (non-flying, non-diving; *F0D0*) consists almost entirely of extant species.

### Inclusion and exclusion of taxa

#### Body size inclusion and exclusion

The datasets in Fabbri *et al.* for lifestyle evaluation of large-bodied spinosaurids include taxa across a wide range of body size. Taxa of body size more comparable to spinosaurids were removed prior to analysis. Some taxa included in the datasts are truly miniscule by comparison: the smallest have femoral diameters less than 1 mm and body massed of 7 g or less. *Spinosaurus* achieved masses approximately 10^6^ times larger.

If the small end of vertebrate body size is correlated with low *Cg*, then including small crouching terrestrial taxa, such as shrews and voles, may bias the training dataset to associate the low *Cg* of small taxa with a non-diving lifestyle. Conversely, increased *Cg*, which is correlated with large body size, is an alternative hypothesis to account for elevated *Cg* in the (less hollow) hind limb long bones of *Spinosaurus* [26]. Fabbri *et al.* pruned “graviportal” taxa from the full dataset of 200 taxa (Table 2, ds1) prior to the pFDA analysis, effectively removing all large-bodied terrestrial taxa closest in body size to *Spinosaurus*.

Taxa removed can be assessed from two files provided by Fabbri *et al.*: “femur compactness all no graviportals and pelagic.csv” (Table 2, ds3) and “rib compactness all no graviportals and pelagic.csv” (Table 2, ds4). These files are read by their R script that performs the pFDA analysis. Taxa removed include the woolly mammoth *Mammuthus primigenius* with a large femoral diameter (172 mm). The largest taxon remaining in the pruned dataset is the digitigrade (hooved) European bison *Bison bonasus* (femoral diameter 63.6 mm), which is considerably smaller than adult *Spinosaurus* (femoral diameter ∼81 mm). Although *Hippopotamus amphibicus*, with adult body mass over one metric ton, exceeds that of *Bison bonasus* and would be the heaviest extant animal in the pruned dataset, the individual included has a smaller femoral diameter (59.3 mm).

Fabbri *et al.* defined “graviportal” succinctly as species with “columnar limbs” [18: 855]. The R script used for taxon exclusion, however, removed a more diverse set of species. Three rhinoceroses (*Ceratotherium simum*, *Rhinoceros sondaicus*, *Rhinoceros unicornis*) and the extinct hippopotamus *Hexaprotodon* were culled, despite their distinctly flexed limb postures. Culled taxa also include the living pygmy hippo *Choeropsis liberiensis* and extinct *Desmostylus hesperus*, neither of which can be construed as graviportal. Fabbri *et al.* also asserted that “graviportality does not affect rib compactness” [18], but recent studies suggest that bone density in these two skeletal components is often correlated [38,85].

The term “graviportal,” finally, has no universal definition [43,48,86,87]. Originally it referred to a specific posture or mode of terrestrial locomotion. More recent studies, however, have shown that both posture and locomotor mode lie on a continuum, with position better captured by osteological indices (long bone length/width ratios) [43,44,86,88,89]. Some regard “graviportal” as a mode of quadrupedal locomotion, whereas others outline criteria for “graviportal bipeds” [47,90–92] that would include spinosaurids [84,85]. The precise definition, however, is irrelevant as higher *Cg* tends to characterize all large-bodied animals irrespective of posture or mode of locomotion [17,44–46].

#### Removal of “deep diving” or “pelagic” taxa

Fabbri *et al.* also remove taxa that they deem “pelagic” or “deep diving.” Although these terms are used for very distinctive—and sometimes nonoverlapping—lifestyles, Fabbri *et al.* used them interchangeably. Setting that distinction aside, Fabbri *et al.* used anatomical observations to distinguish “deep diving” taxa:

> High bone density is therefore an excellent indicator for the initial stages of aquatic adaptation, but poorly distinguishes between wading, deep diving, and terrestrial habits. These limitations can be overcome using anatomical observations because deep diving shows other transformations of the body plan, such as presence of fins and flippers.

We agree that finned or flippered taxa are poor models for comparison to spinosaurids which manifestly *do* have terrestrial limbs. Even if one supposed that spinosaurids were on an evolutionary trajectory to become fully aquatic (highly speculative, as no fully aquatic descendants have been discovered), the best points of comparison would still be other taxa with terrestrially useful limbs at an early stage of secondary aquatic adaptation.

Applying their stated anatomical criteria for identifying “deep diving” would mean removing *all* taxa that show the anatomical features they identify. Taxa removed in the file “femur compactness all no graviportals and pelagic.csv,” however, do *not* follow the anatomical criteria specified —*i.e.*, transformations of the body plan or the presence of fins and flippers. The extant cetaceans Bryde’s whale *Balaenoptera brydei* and orca *Orcinus orca* were removed, however the flippered extinct whale *Basilosaurus* was retained. The extinct seal (*Callophoca obscura*) was removed, yet the extant harbor seal (*Phoca vitulina*) was retained. The plesiosaur *Cryptoclidus* was retained despite recent work suggesting an open-ocean lifestyle for many plesiosaurs [93]. Despite their flippers, fins, and flukes, sirenians, plesiosaurs, ichthyosaurs, and nothosaurs were all retained. Instead of following the anatomical signature, Fabbri *et al.* seem to have simply removed taxa with low *Cg* from the *D = 2* group.

#### Ignored and redundant taxa

Fabbri *et al.* incorporated 78 taxa in the rib dataset from Canoville *et al.* [38] but ignored an additional 43 extant species that merit inclusion, including semiaquatic, large-bodied terrestrial varanids (*Varanus salvator*, water monitor; *Varanus komodoensis*, Komodo dragon). The Triassic aquatic reptile *Nothosaurus*, in contrast, is represented by six specimens (Table 3). *Nothosaurus* and its close relatives (three genera) are overrepresented, accounting for ∼15% of the nonflying/diving (*F0D2*) dataset and 21% of the extinct taxa. Bone density in *Nothosaurus*, in addition, is significantly negatively correlated with body size (femoral diameter) (S1 Fig; *R*^2^ = 0.84). The bone density data for this taxon, thus, would not scale to the body size of a spinosaurid (where *Cg* would drop to near zero) and probably should be removed from the analysis for that reason alone.

**Table 3.**
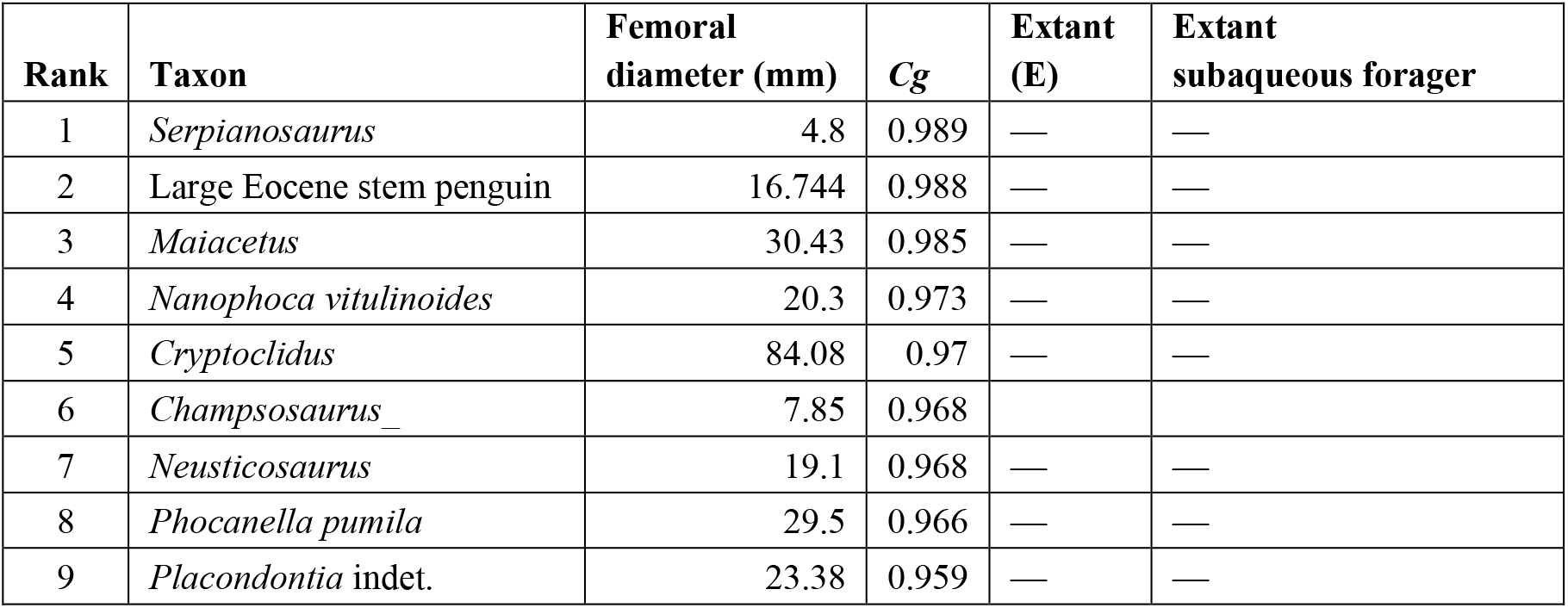
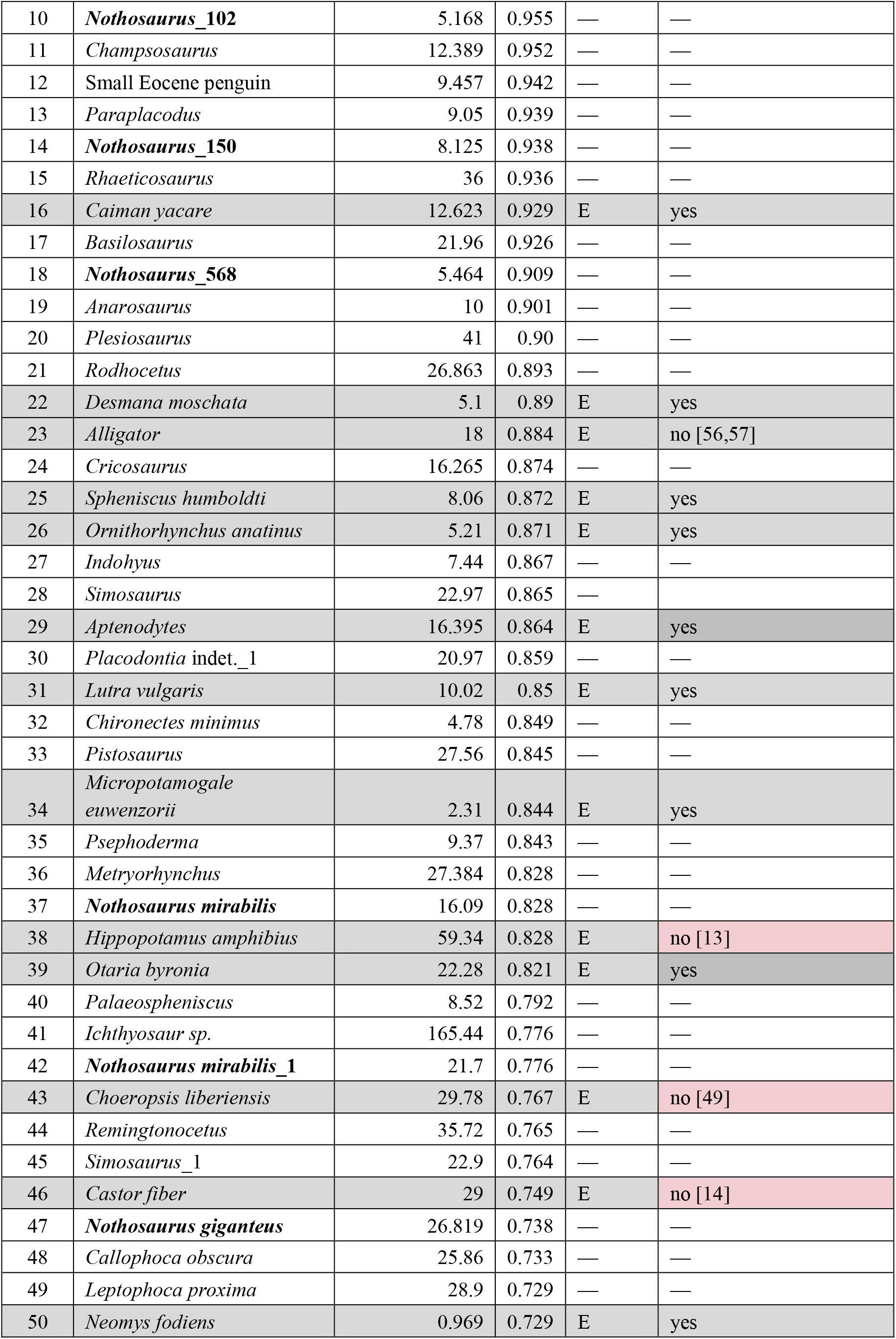
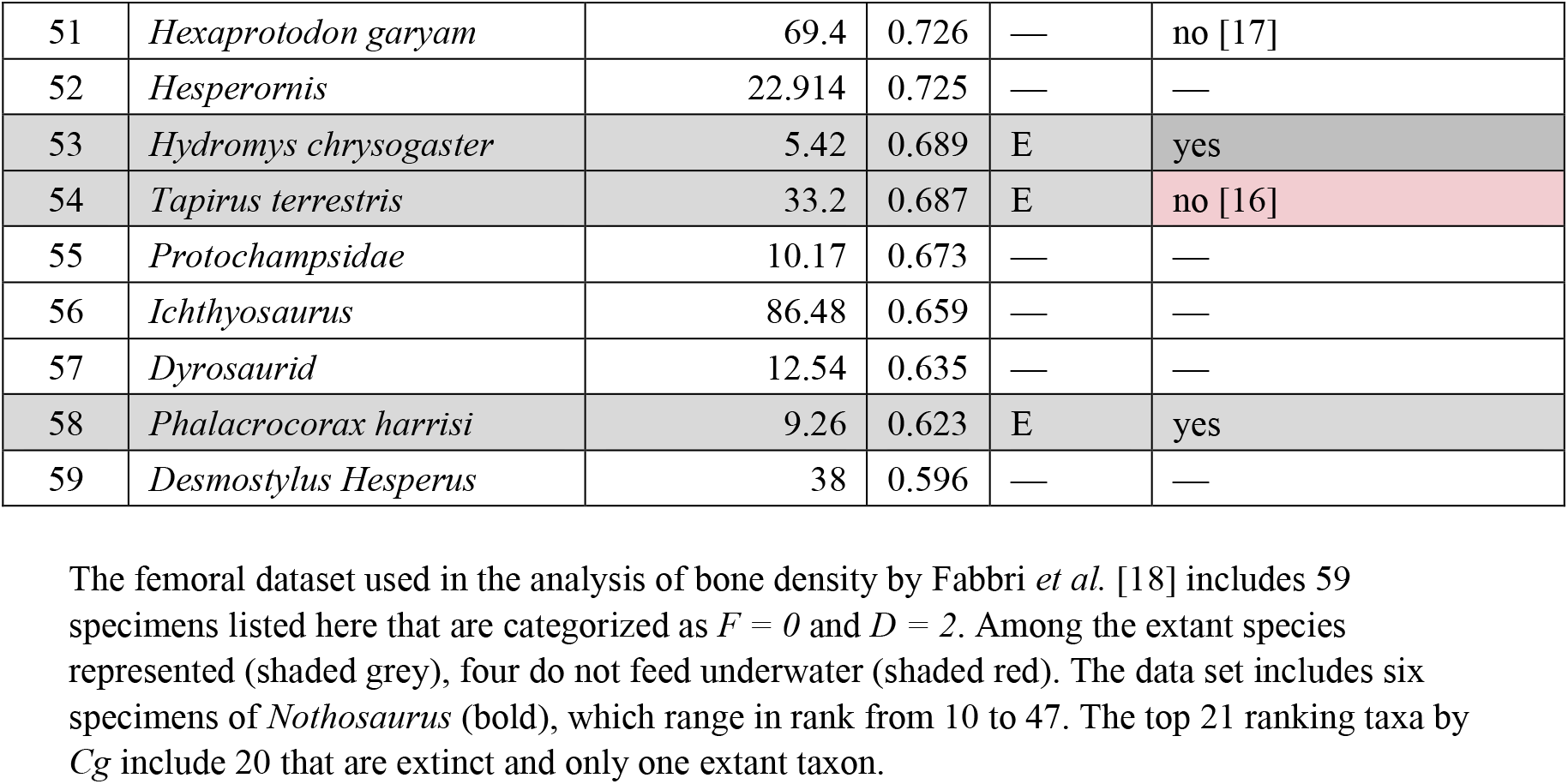
Nonflying, diving subsample of taxa (*F0D2*) based on femoral data are ranked by bone compactness (*Cg*).

### Other dataset concerns

#### Bone density disparity between extinct and extant taxa

Bone compactness (*Cg*) ought to be a determinant of secondary aquatic adaptation in both extinct and extant taxa. For a given lifestyle, bone compactness should not be systematically higher or lower in living versus extinct species. Striking differences in bone compactness, nonetheless, are apparent between extinct and extant taxa of similar lifestyle in some of the datasets (Table 3). The femoral dataset for nonflying divers (*F0D2*), for example, shows a strong bias in values of *Cg*, with values among extinct taxa higher than those of extant taxa. Of the top 21 taxa ranked by *Cg*, 20 are extinct (Table 3). The two extant taxa having the highest *Cg* rank 16 and 22 in this dataset (Table 3). The disparity is particularly worrisome because spinosaurids were clearly nonfliers (*F = 0*) and therefore must either have been nonflying divers (*F0D2*) or terrestrial (*F0D0*). The greater average density of extinct taxa may be the result of secondary mineral deposition/precipitation in porous bone during fossilization. The differences might instead arise from the specific choices of extinct taxa included or some other reason.

The nonflying diver (*F0D2*) subset of 59 taxa has more extinct (43) than extant (16) species, an imbalance that may have biased results given their differing *Cg*. To evaluate that possibility, we used a permutation test of *Cg* rank, the null hypothesis being no difference between the *Cg* values of extinct versus extant taxa (see Materials and methods). We calculated *p* values for the null hypothesis (Table 4). In a second, “coin-flip” test, a binomial distribution was used to determine *p* values for an alternative null hypothesis that the counts of extinct and extant specimens in each dataset resulted from random chance. These tests were performed on all four dataset variations (Table 1, ds1–ds4) for both *F0D0* and *F0D2* subsets of femur and rib data.

**Table 4.**
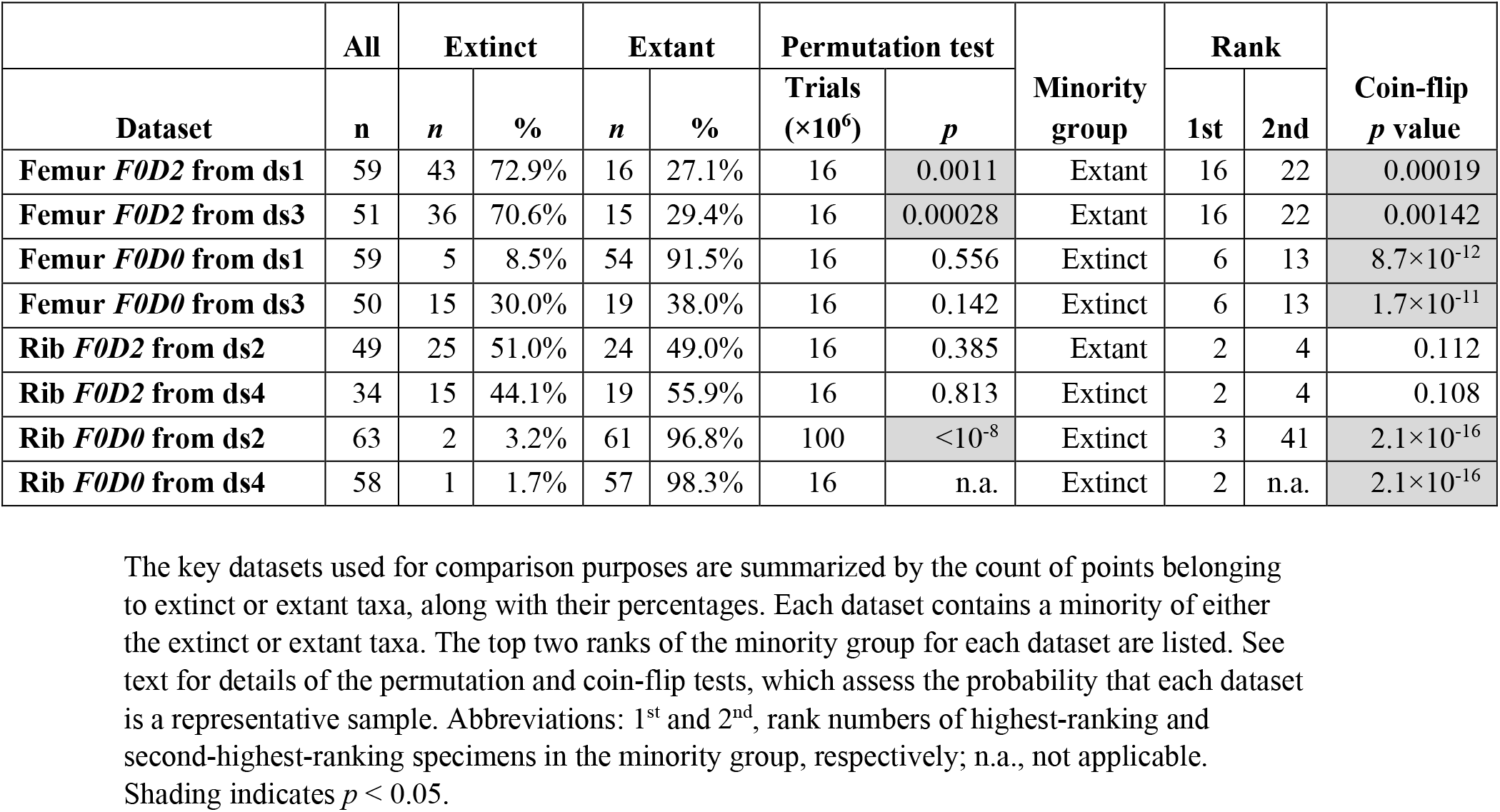
Permutation and coin-flip tests of rib and femoral datasets.

The permutation test on *F0D0* femoral data from ds1 and ds3 have *p* ≤ 0.0011; we therefore reject the null hypothesis that the distribution of *Cg* values is the same for extinct and extant taxa (Table 4, shaded). The test results for the *F0D0* rib data from ds2 similarly rejects the null hypothesis with high probability (Table 4, shaded). Overall, these statistical tests and *p* values suggest the sampling for *Cg* is incomparable between extinct and extant taxa. This violates the foundational assumption that *Cg* can be used as a marker for both groups.

The results of the coin-flip tests on *F0D0* (femora and rib) datasets and respective variants further indicate that it is extraordinarily unlikely that the pronounced imbalances between extinct and extant taxa in these datasets are the result of random chance (Table 4, shaded).

The rib (ds2) *F0D2* dataset, in contrast, sampled roughly equal numbers of extinct and extant taxa, and our statistical test are consistent with the null hypothesis. Discordant findings between rib and femoral datasets run contrary to the claim that the same *Cg* signal is preserved in both limb bones and axial skeleton.

A further concern about the imbalance between extinct and extant taxa is that pFDA relies on correction of bias based on a phylogenetic tree with branch lengths. Phylogenetic relationships and branch lengths are often less well known for extinct than extant taxa. Biased results may be generated if there is a marked imbalance between extinct and extant taxa, as occurs in the femoral dataset.

#### Inclusion of parameters and data shown to be non-significant

Fabbri *et al.* began their analysis with a PGLS linear regression model to determine the statistical significance of correlations among their categorical variables (*F*, *D*) and two measured parameters (diameter, *Cg*), alone and in combination. They found correlations between the *D = 2* (habitual diving) and femoral or rib *Cg* that are statistically significant (*p* < 0.001) although extremely weak (*R*^2^ = 0.172 for femora, 0.108 for ribs). Judging statistical significance solely on *p* values can be misleading [94]. They found no statistically significant correlation of *Cg* with *D = 0* (nondiving) or *D = 1* (occasional diving), further weakening the argument that *Cg* alone can predict ecology/behavior.

When bone diameter and *Cg* were included in the regression, the result was considered significant. Fabbri *et al.* computed the Akaike Information Criterion (AIC) weights [18: Supplementary tables 3,4]. Their own calculation thus shows that in the femoral dataset, the model that includes diameter *Cg* and *MD* is (under the assumptions behind AIC weights) has about 49 times less explanatory power than a model that includes *Cg* alone. For the rib dataset, the model including *Cg* and *MD* similarly has AIC weights 34 times smaller than those of the model that includes *Cg* without bone diameter.

Despite these results favoring simpler models based on *Cg* alone, Fabbri *et al.* conducted their main analysis using a two-dimensional (*MD, Cg*) dataset. A similar situation occurs for the cases where *D = 1* and *F = 1, 2*. For *D* = *1* in combination with *MD*, the correlation falls well short of statistical significance with *p* = 0.648 for femoral data and *p* = 0.492 for rib data. The correlation is also very weak for *F = 1, 2*.

In conventional practice, such PGLS results would require excluding irrelevant taxa with *D = 1* and *F = 1, 2*, from the training set to avoid further weakening the correlation with *D = 2*. Instead, the R script of Fabbri *et al.* treats these taxa as *D = 0* in the training dataset by means of an auxiliary variable (“diving_or_not”). In our analysis, we used both the Fabbri *et al.* method (denoted “Many vs. *F0D2*”) and a second analysis with the better-justified use of *F0D0* and *F0D2* taxa as the only members of the training dataset.

The pFDA method allows the assignment of prior probabilities for the membership of a test taxon for Bayesian calculation of posterior probabilities of class membership. The default approach used in pFDA is that the prior probabilities are proportional to class membership [95], which is the count of taxa in the *F0D2* group versus the count in the alternative group (either “Many” or *F0D0*). While this may make sense in some contexts, using class membership to bias the classification in this case is problematic. Whether spinosaurids are “subaqueous foragers” or not should not depend on the vagaries of how many taxa investigators collect. Analysis should employ unbiased priors, *i.e.,* an equal probability of membership in *F0D2* or not.

### Axial pneumaticity and buoyancy

#### Impact of pneumaticity

Research on increased bone density (pachyostosis, osteosclerosis) has focused on its potential role as ballast decreasing buoyancy in secondarily semiaquatic and aquatic amniotes. Buoyancy, nonetheless, is impacted far more strongly by pneumaticity in theropods like *Spinosaurus* because pneumatic invasion replaces soft tissue or bone of density greater than 1 g/ml with air. Cancellous or dense bone infilling, by comparison, replaces soft tissue (blood vessels/marrow of density near that of water, 1 g/ml) with bone of only slightly greater density (∼1.2 g/ml). Pneumaticity in the axial and appendicular bones in theropods (including birds) thus *increases* buoyancy far more than a comparable volume of dense bone *decreases* it. The simpler adaptation in extant nonflying diving birds to decrease buoyancy is to reduce or eliminate pneumaticity.

Extant birds include some that fly but do not dive (*F2D0*) and others that regularly perform both functions (*F2D2*). Bone compactness (*Cg*) is generally higher in those divers that do not fly, but their bone compactness polygons broadly overlap (Fig 1A and 1B). Bone compactness among proficient avian divers thus does not cleanly distinguish those that have abandoned flight and might be thought to have evolved heavier bone mass as ballast. Decreased pneumaticity, on the other hand, has been shown to be correlated with the evolution of pursuit diving in birds. The postcranial skeletons of darters, loons, cormorants, and penguins are all less pneumatic than those of other birds that fly or forage while wading or floating [96].

#### Spinosaurid axial pneumaticity

Fabbri *et al.* remarked that “osteosclerosis is observed across multiple skeletal elements in *Baryonyx* and *Spinosaurus*” and that the absence of similar bone density in *Suchomimus* is “secondary loss.” They concluded that “subaqueous foraging is ancestral for Spinosauridae” [18: 856]. On its face, the argument is ambiguous regarding the ancestral spinosaurid condition, given opposing conditions in closely related sister taxa (*Baryonyx*, *Suchomimus*) to *Spinosaurus*. A shift to osteosclerosis in ancestral spinosaurids with a reversal in *Suchomimus* (their argument), in other words, is parsimoniously equivalent to two independent shifts toward osteosclerosis in *Baryonyx* and *Spinosaurus*.

Bone structure in these three spinosaurids, however, provides ample evidence of well-developed paraxial pneumaticity that would supersede any ballast effect from variable long bone infilling. There are also large medullary cavities (presumably fat-filled) that hollow the centra at the base of the tail that would further reduce bone density [26]. The internal volume of cervical paraxial pneumaticity (∼25% by volume) is well documented in *Spinosaurus* [97] with evidence that external paraxial air sacs extended along the entire dorsosacral column (Fig 2). In *Suchomimus* and *Baryonyx*, most precaudal vertebrae have internal pneumatic chambers (camerae) within the centra and deep fossae for pneumatic sacs on the neural arch (Fig 2B–2D).

**Fig 2.**
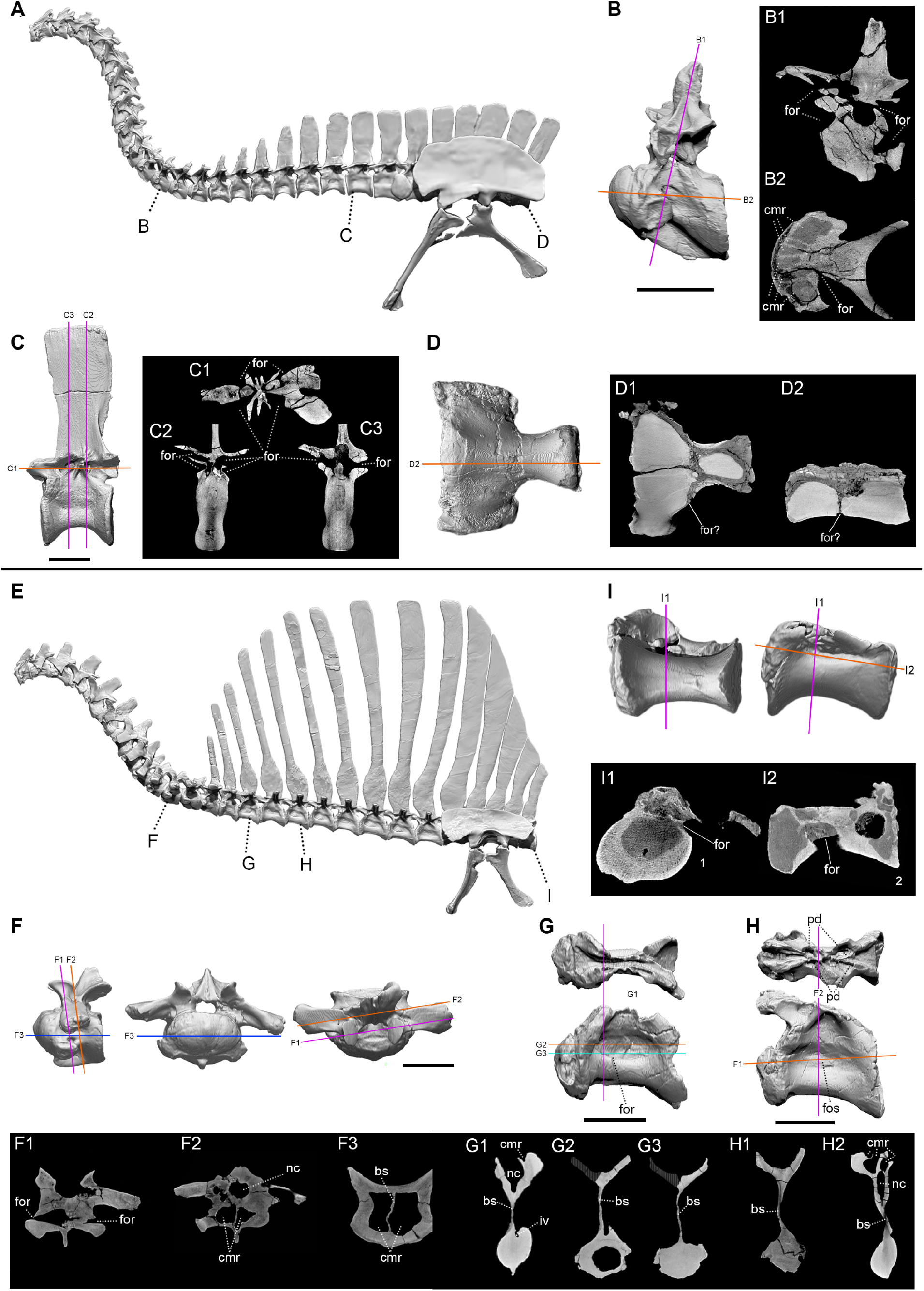
Pneumatic features in the dorosacral column in spinosaurids. (A) *Suchomimus tenerensis* (MNBH GAD500) precaudal column and pelvic girdle showing pneumatic features in (B) D2 in lateral view with coronal (B1) and axial (B2) CT cross sections, (C) D13 in lateral view with axial (C1) and sagittal (C2, 3) CT cross sections, and (D) S2 in ventral view with axial (D1) and coronal (D2) CT cross sections. (E) *Spinosaurus aegyptiacus* precaudal column and pelvic girdle showing pneumatic features in (F) ∼D2 in lateral, anterior and dorsal views with coronal (F1, 2) and axial (F3) CT cross sections, (G) ∼D6 in dorsal and lateral views showing coronal (G1) and axial (G,2, 3) CT cross sections, (H) ∼D8 in dorsal and lateral views with axial (H1) and coronal (H) CT cross sections, and (I) S3 centrum in ventral and lateral views with coronal (I1) and axial (I2) CT scan sections. Neotypes FSAC-KK-11888 (panels G, H, I) and BSPG-2006-I-54 (panel F). CT section lines are color-coded by orientation (*magenta*, coronal; *blue*, axial-horizontal; *black*, sagittal/parasagittal). Scale bars are 10 cm. Abbreviations: bs, bony septum; c, cervical vertebra; cmr, camera; d, dorsal vertebra; for, foramen; fos, fossa; nc, neural canal.

Precaudal vertebral pneumaticity is present in *Spinosaurus* to an even greater degree than in its baryonychine relatives *Baryonyx* and *Suchomimus*. The pneumatic foramina and camerae in the anterior dorsal vertebrae (Fig 2F) are larger than in *Suchomimus*, and mid dorsal centra have marked, oval pneumatic fossae that reduce intervening bone to a thin sagittal septum (Fig 2G and 2H). Similarly, mid-sacrals have large pneumatic foramina and internal camerae (Fig 2I).

These findings challenge the hypothesis that increased long bone or rib density served as ballast for diving in either *Baryonyx* or *Spinosaurus*, because decrease in buoyancy from increased long bone or rib compactness (*Cg*) is more than offset by pneumaticity in the axial skeleton. In *Spinosaurus*, in addition, bone mass loss from reduction in hind limb length is greater than the mass gained from medullary infilling [26]. A flesh model reconstruction of this genus that incorporates body partition densities confirms net buoyancy as a serious impediment for underwater submergence [26].

### Variation in *Cg*

Significant variation may exist in *Cg* values unrelated to ecology or behavior, such as variation among individuals of a species, changing values during ontogeny, variation in different skeletal elements, and variation at different locations along the shaft of a single bone. Measurement error may accrue from several sources, such as calculating the index from thin sections or CT scans, decisions taken in thresholding images, and the degree of repair of cracks and missing bone in damaged or incomplete specimens. Attempting to replicate *Cg* values from the original bone thin sections used in Fabbri *et al.* demonstrate that all these sources of variation are present and can figure importantly in the interpretation of bone compactness data.

#### Bone compactness variation among individuals, during ontogeny, and along bone shafts

Most species data points in bone compactness studies, including Fabbri *et al.*, are based on a single measurement from a single individual. The reliability of bone compactness data requires data on individual variation, variation during growth and maturation, and variation within individual bones, although very little research has established any of these across vertebrates.

In the manatee *Trichechus manatus latirostris*, *Cg* values in ribs were measured from thin sections in 12 individuals that included males and females, as well as growth stages from 50 to 1057 kg [98]. *Cg* values ranged from 0.8389 to 0.9962, with a mean of 0.9109 and a standard deviation of 0.0417, a relative range of 13.9% (relative range: high minus low values divided by the mean and then multiplied by 100). Excluding the youngest three individuals of least body mass reduces the size range (161 to 1057 kg) without reducing the relative range significantly (13.2%).

We compiled multiple measurements of *Cg* from multiple individuals within or across studies for taxa present in the datasets of Fabbri *et al.* (Table 5). We did the same for taxa in a recent multi-individual study of flightless birds (Table 6) [47]. Multiple measurements of *Cg* in the same bone of the same species often exhibit relative ranges exceeding 10%, and the median relative range among the entries in Tables 5 and 6 is 18.6%.

**Table 5.** Examples of individual variation in *Cg* measurements across sources used by Fabbri *et al.* [18].

| Taxon | Bone | References | *MD* (mm) | *Cg* | *Cg* range (%) |
| --- | --- | --- | --- | --- | --- |
| *Phoca vitulina* | Rib | [99] | 7.8 | 0.436 | 22.0% |
|  |  | [38] | 11.49 | 0.544 |  |
| *Sphenicus humboldti* | Rib | [18] | 4.08 | 0.908 | 24.3% |
|  |  | [38] | 4.98 | 0.711 |  |
| *Giraffa camelopardalis* | Rib | [38] | 21.4 | 0.544 | 1.6% |
|  |  |  | 18.16 | 0.553 |  |
| *Metriorhynchus* | Femur | [11] | 27.38 | 0.828 | 46.1% |
|  |  | [100] | — | 0.518 |  |
| *Ceratherium simum* | Femur | [11] | 78.6 | 0.669 | 20.5% |
|  |  |  | 92.6 | 0.819 |  |
|  |  |  | 70.2 | 0.827 |  |
| *Mammuthus* | Femur | [11] | 102.9 | 0.846 | 14.9% |
|  |  |  | 139.5 | 0.898 |  |
|  |  |  | 172 | 0.773 |  |
| *Nothosaurus mirabilis* | Femur | [101] | 16.09 | 0.828 | 6.5% |
|  |  |  | 21.7 | 0.776 |  |
| *Nothosaurus* | Femur | [102] | 5.168 | 0.955 | 4.9% |
|  |  |  | 8.125 | 0.938 |  |
|  |  |  | 5.464 | 0.909 |  |
| *Simosaurus* | Femur | [103] | 22.97 | 0.865 | 12.4% |
|  |  |  | 22.9 | 0.764 |  |
| *Diceros bicornis* | Humerus | [11] | 79.6 | 0.866 | 7.9% |
|  |  |  | 70.2 | 0.937 |  |
| *Ceratherium simum* | Humerus | [11] | 90.7 | 0.771 | 12.1% |
|  |  |  | 89.3 | 0.87 |  |
| *Dicerorhinus sumatrensis* | Humerus | [11] | 50.6 | 0.941 | 17.5% |
|  |  |  | 52.8 | 0.7895 |  |
| *Scutellosaurus lawleri* | Humerus | [11] | 6.1 | 0.767 | 18.6% |
|  |  |  | 6.2 | 0.748 |  |
|  |  |  | 139.5 | 0.898 |  |

Data compiled for taxa in Fabbri *et al.* [18] that have multiple *Cg* and *MD* measurements from the same bone. *Cg* variation (largest to smallest) ranges from 1.6 to 46.1% relative to respective means. ( $Cg\ range = (\max - \min) / \text{mean} \times 100$ ). Most are based on only two or three specimens. Abbreviations: *MD*, maximum bone diameter; *Cg*, global bone compactness; max, maximum; min, minimum.

**Table 6.**
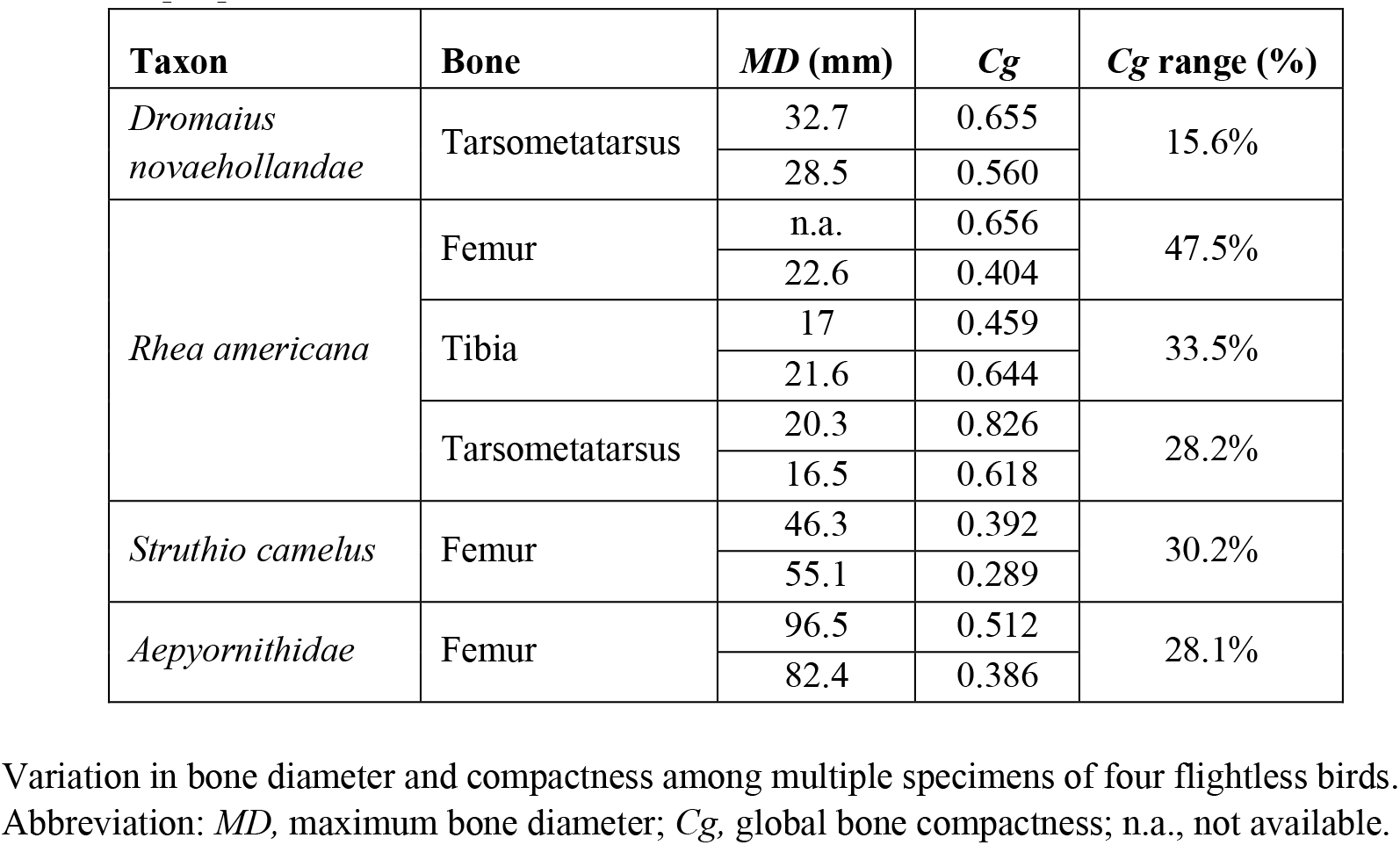
Examples of individual variation in *Cg* measurements of flightless birds from Canoville *et al.* [47].

| **Taxon** | **Bone** | ***MD* (mm)** | ***Cg*** | ***Cg* range (%)** |
| --- | --- | --- | --- | --- |
| *Dromaius novaehollandae* | Tarsometatarsus | 32.7 | 0.655 | 15.6% |
|  |  | 28.5 | 0.560 |  |
| *Rhea americana* | Femur | n.a. | 0.656 | 47.5% |
|  |  | 22.6 | 0.404 |  |
|  | Tibia | 17 | 0.459 | 33.5% |
|  |  | 21.6 | 0.644 |  |
|  | Tarsometatarsus | 20.3 | 0.826 | 28.2% |
|  |  | 16.5 | 0.618 |  |
| *Struthio camelus* | Femur | 46.3 | 0.392 | 30.2% |
|  |  | 55.1 | 0.289 |  |
| *Aepyornithidae* | Femur | 96.5 | 0.512 | 28.1% |
|  |  | 82.4 | 0.386 |  |

Variation in bone diameter and compactness among multiple specimens of four flightless birds. Abbreviation: *MD*, maximum bone diameter; *Cg*, global bone compactness; n.a., not available.

Median variation of 18.6% is a very large percentage, considering the limited range in *Cg* (0.43–1) reported by Fabbri *et al*. Our tabulation suggests that variation within a single taxon could account for as much as ∼33% of the total variation across taxa. Even if that variation proved atypical, even a few taxa with large discrepancies—such as the maximum range in *Cg* observed in flightless birds (47.5% in femora of *Rhea americana*, Table 6)—could bias a discriminant analysis. Most of the taxa in Tables 5 and 6 are represented by only two or three specimens. Better assessment of variation in *Cg* will require larger sample sizes for *Cg* among conspecifics and across a greater range of taxa. These data underscore the caution needed when drawing conclusions based on isolated measurement of *Cg*.

#### Variable long bone infilling

Medullary cavities in long bones of the fore and hind limb of *Spinosaurus* are variably infilled (Fig 3B and 3C). Fabbri *et al.* based their estimated *Cg* for *Spinosaurus* on one thin section taken from one fully infilled subadult femur (Fig 3D). A second femur of *Spinosaurus* [104] (Fig 3A and 3B) is slightly larger than the infilled neotypic femur (Fig 3C) but has a significant medullary cavity lined with cancellous bone that would register as significantly less dense in thin section at midshaft. With only two subadult femora available for *Spinosaurus*, the meaning of such variation is uncertain. In extant birds, intraspecific variation has also been recorded in the volume and location of medullary cavities [105]. These examples underscore the need to sample species more broadly rather than to accept a single measurement of bone compactness as representative of a given species.

**Fig. 3.**
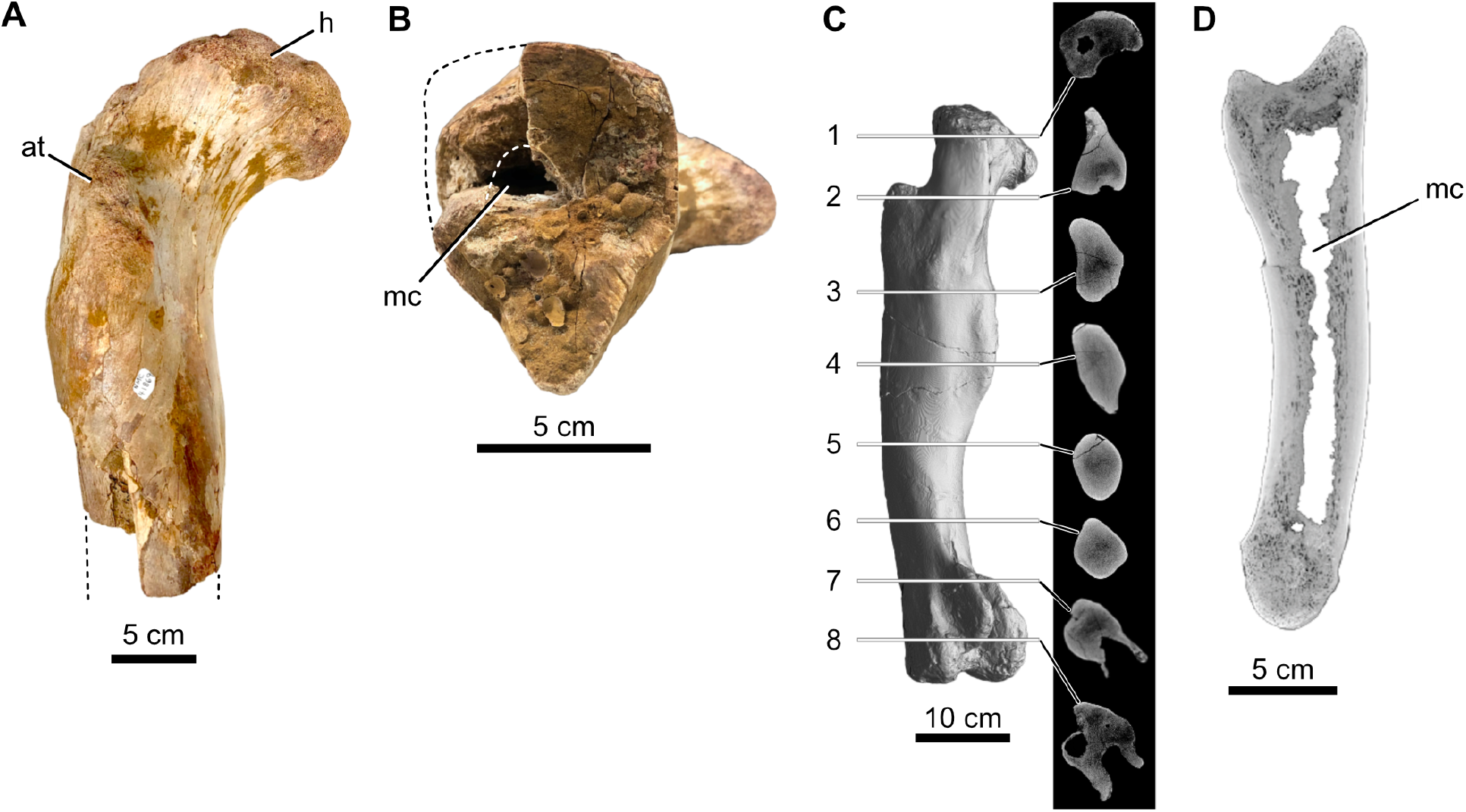
Femora and a manual phalanx of cf. *Spinosaurus aegyptiacus* from the Kem Kem Group in Morocco. (A, B) proximal one-half of an isolated right femur in anterior view and distal midshaft cross-sectional views (CMN 41869); (C) CT scan of the left femur of the neotype with eight cross sections (FSAC-KK 11888); (D) CT scan of an isolated right phalanx I-1 in sagittal cross section (UCRC PV8). Abbreviations: at, anterior trochanter; h, head; mc, medullary cavity.

#### Bone density metrics and variation

The *Cg* metric used by Fabbri *et al.* as a proxy for bone density is one of many metrics available to capture bone compactness. Over the last decade, other metrics generated by the Bone Profiler program (*S, P, Min, Maxrad, Cc, Cp*) have been shown to better correlate with lifestyle in extant amniotes [3,8,82,99]. Comparing results generated by metrics other than *Cg* is beyond the scope of the present study.

Fossil specimens are often damaged pre- or postmortem, as well as during collection and preparation. Spread cracks are common and often infilled with matrix. Exterior portions of bone are frequently eroded or broken away. Digital repair of these imperfections are needed to more accurately assess *Cg*. Repairs of this kind generally increase *Cg* slightly because repaired bone is usually filled in as solid black on a binarized image, no matter the vascularity present in adjacent bone. Fabbri *et al.*, for example, repaired the broken edge of a bone cross section from a marine reptile, filling in the missing piece with solid black (Fig 4). Comparing and remeasuring bone compactness in the sections suggests that the black infill for the missing bone may have raised the *Cg* by ∼5%.

**Fig 4.**
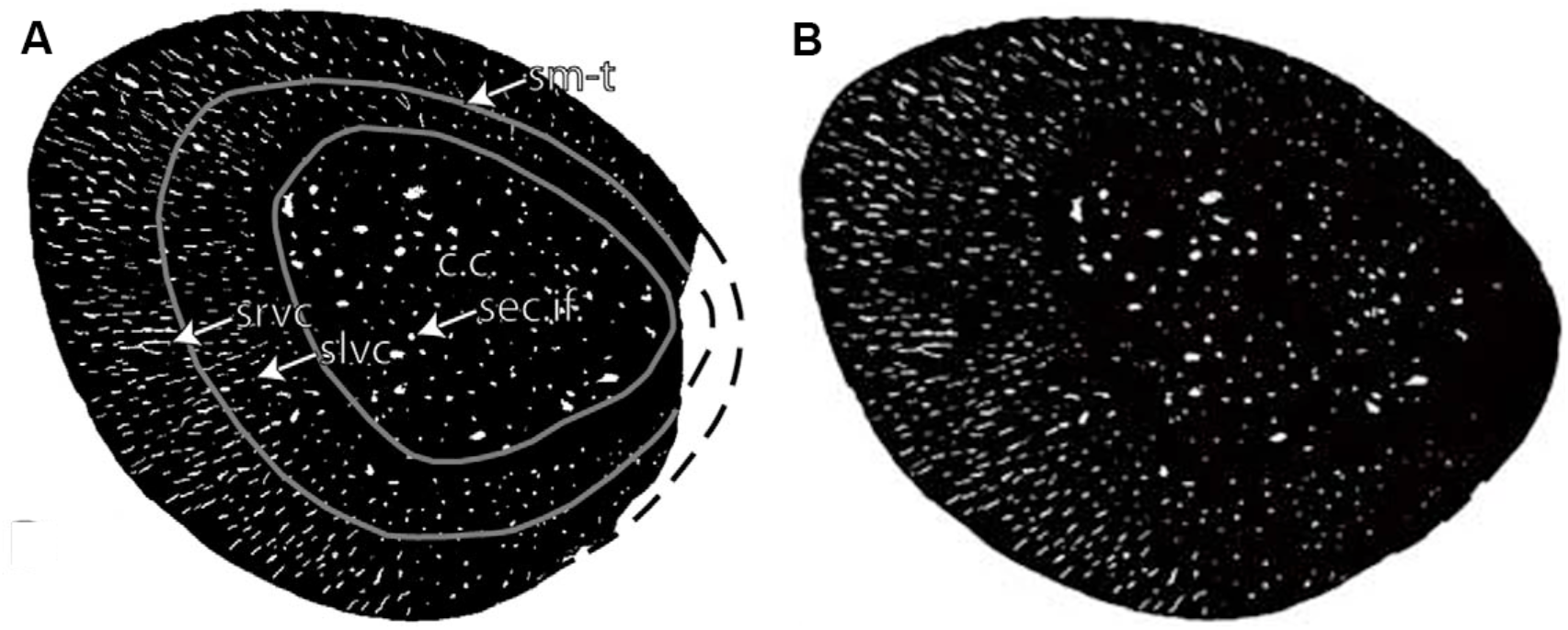
Comparison of the original binarized and digitally repaired versions of a bone cross section from the marine reptile *Neusticosaurus*. (A) Threshold-filtered cross section of *Neusticosaurus edwardii* [106: fig 5, panel D] with dashed lines showing bone lost to erosion. (B) Digitally-repaired image using solid black infill from Fabbri *et al.* [18: Extended data Fig 4].

One can anticipate *Cg* varying along the shaft of long bones or ribs as bone diameter changes and cross sections encounter external trochanters or condylar ends. This can be a significant source of sampling variation. A sequence of thin sections along the shaft of a dorsal rib of the marine reptile *Nothosaurus* [107] shows *Cg* variance of ∼35% (Fig 5). Although this particular specimen was not used by Fabbri *et al.*, nothosaurs compose a significant component of their dataset. *Cg* variation along the shaft of femora or ribs in other taxa has not been documented.

**Fig 5.**
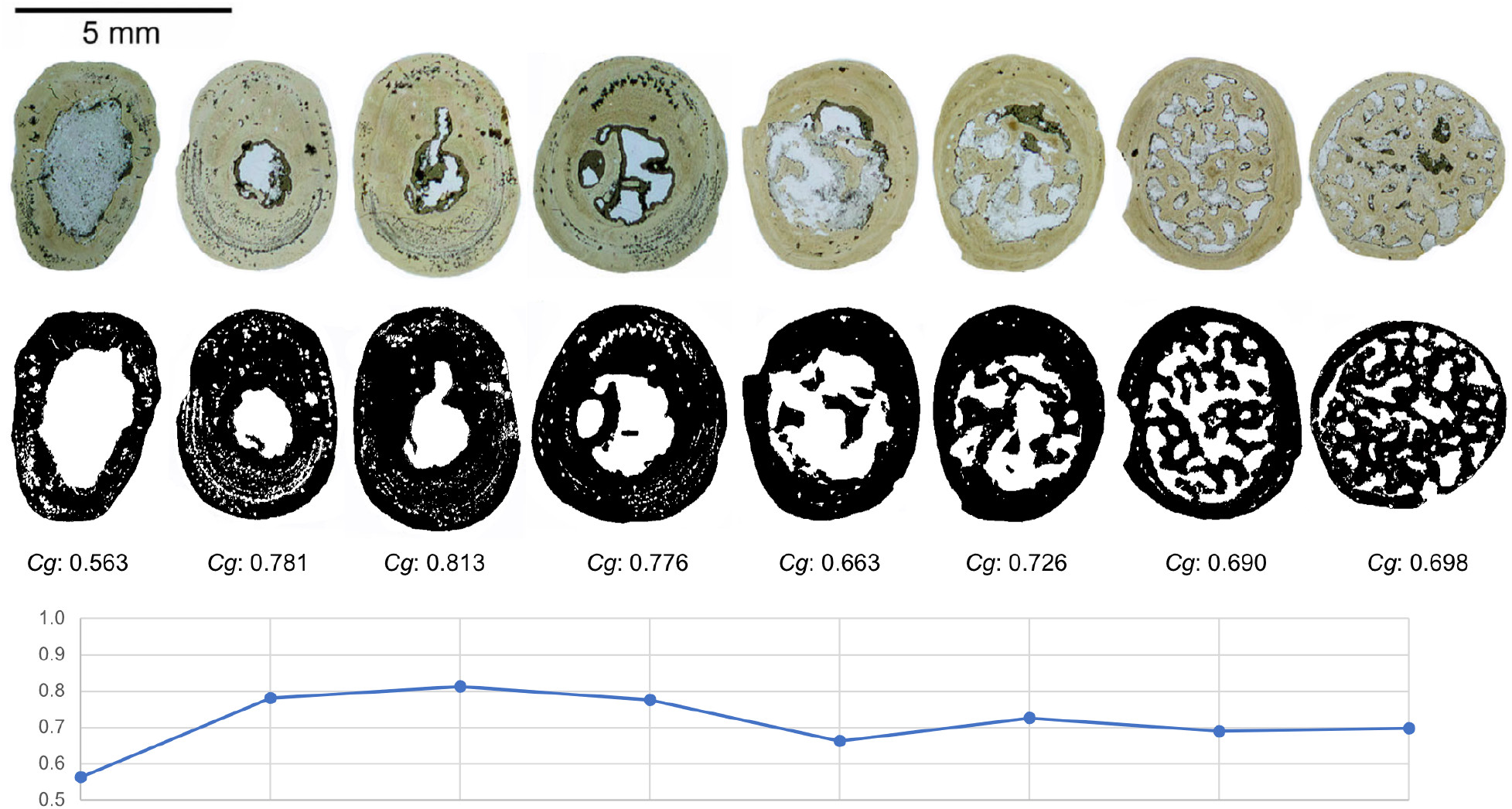
Variation of *Cg* at different points along a single *Nothosaurus* dorsal rib. Images from Klein *et al.* [107: Fig 3, panels B1–B8] were used to measure variation in *Cg* (∼35%) along the rib shaft.

#### Bone density in spinosaurid femora

We report below on significant sampling and methodological variations in reported *Cg* values for all three spinosaurids that were used by Fabbri *et al.* as the basis for evaluating spinosaurid lifestyle as either habitually diving (a “subaqueous forager” seeking underwater prey) or nondiving (a wading terrestrial shoreline predator).

Sampling variation in fossil taxa involves the location of a section on a long bone or rib (the bones most commonly sampled), the developmental stage and potentially the sex of the specimen, and diagenetic and taphonomic factors (*e.g.*, fracturing, deformation, infilling, and external erosion).

Methodological variation in bone density determination includes, but is not limited to the chosen bone compactness metric; the type of bone section analyzed (CT digital scan, mounted thin section); the threshold value used to binarize a section image; and contour, masking, or repair steps taken prior to measurement of *Cg*. These many sources of variation ensure that independent researchers will not obtain the same quantitative value for bone compactness from a single specimen, even when deriving measurements from the same cross section.

##### Spinosaurus aegyptiacus

Fabbri *et al.* reported a very high *Cg* of 0.968 for *Spinosaurus* calculated from a binarized image based on an image taken of a two-part thin section from the femoral shaft of the neotype skeleton [22]. That thin section, which was made by one of us (PCS), was taken on the narrow portion of the femoral shaft below the fourth trochanter (Fig 3C, section 5) and shows complete infilling of the medullary cavity. Fabbri *et al.* calculated their *Cg* on a binarized image of this section that showed a small oval core of low density and an open (white) crack separating a portion of the cortex [18: Fig 1b,28: Fig 1a].

Inspection of the thin section under magnification reveals several details that are otherwise impossible to discern from whole or half thin-section images. First, there is no medullary cavity, despite a dark-stained region in the center of the bone shaft (Fig 6). The central core is entirely filled in with bone that is slightly more cancellous. Second, a vertically oriented dark red zone to the right of the core, which shows up as a less-dense zone in the binarized image of Fabbri *et al*., is an artifact of hematitic stain. Under magnification, there is no difference in the bone texture or density of this zone. Third, a crack separating a portion of the lower left thin section occurred during production and mounting of the thin section. The gap created by the crack should be closed digitally prior to *Cg* measurement. These three factors each slightly elevate the *Cg* of the neotype femoral section to 0.998, which is our best estimate. This section shows an essentially fully in-filled condition, whereas the binarized image reported by Fabbri *et al.* shows what appears to be an ovoid, less-dense core that generated a *Cg* approximately 3% lower (0.968).

**Fig 6.**
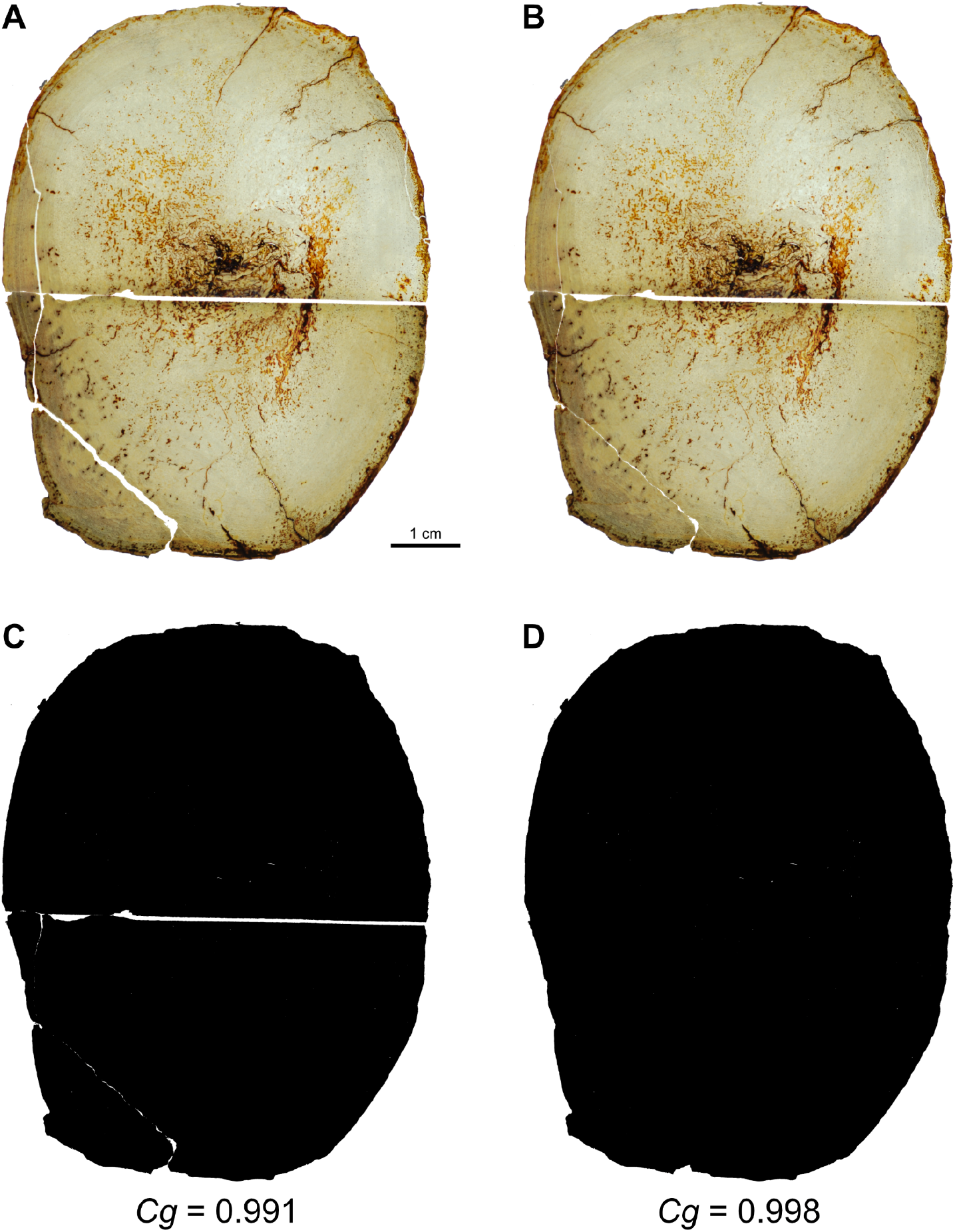
Reevaluation of *Cg* in a two-part thin section from the left femur of the neotype specimen of *Spinosaurus aegyptiacus* (FSAC-KK 11888). (A) Transmitted light image of a two-part thin slice from the mid shaft. (B) Thin slice image modified to close gaps created by natural breaks. (C) Binary image and associated *Cg* value without filling the break between section halves. (D) Binary image and associated *Cg* after filling the gap between section halves.

In response to our critique of Fabbri *et al*. [27], they incorrectly cited our response and introduced misinformation [28]:

> Additionally, based on CT scan imaging, Myhrvold et al.1 accuse us of ignoring a medullary cavity in the femur of the neotypic specimen of Spinosaurus and that we are incorrectly oversampling bone tissue based on a thin section of the femur. As shown in Figure 1, cross sections obtained from the CT scan presented by Myhrvold et al.1 lack adequate contrast and resolution, obscuring any details of its internal structure, contrary to the thin section used in our study.

We never suggested there was a medullary cavity in the neotypic femur, either when the thin section was first published [22] or as later discussed in our critique [27]. The femoral CT scan figured here (Fig 3C, section 5), standard for a large bone CT taken by a medical scanner, is more accurate than the binarized image used to calculate *Cg* in Fabbri *et al.*, as it accurately shows a slight lessening of density toward the core but no medullary cavity.

More significantly, infilling of the medullary cavity of the femur in *Spinosaurus* is variable, as shown by a second specimen of similar body size from the same beds in Morocco [27]. A persistent reduced medullary cavity is exposed by fracturing of the shaft (Fig 3A and 3B) and has been visualized with a CT scan proximal to the break (Fig 7). The absence of matrix infilling of cracks or external erosion sets aside the need for digital repair prior to bone compactness measurement. To calculate *Cg*, we subjected the original gray-scale CT image (Fig 7A) to thresholding using pixel gray values from 0 to 255, transforming it into binary (0, 1) values. Three optional thresholds generated *Cg* values from 0.804 to 0.888, a relative range of 9.9% (Fig 7B–7D). As anticipated, these *Cg* values are significantly lower than reported on the basis of the nearly solid neotypic femur. Our median *Cg* value of 0.849 is somewhat less than that reported by Fabbri *et al.* for *Baryonyx* (0.876).

**Fig 7.**
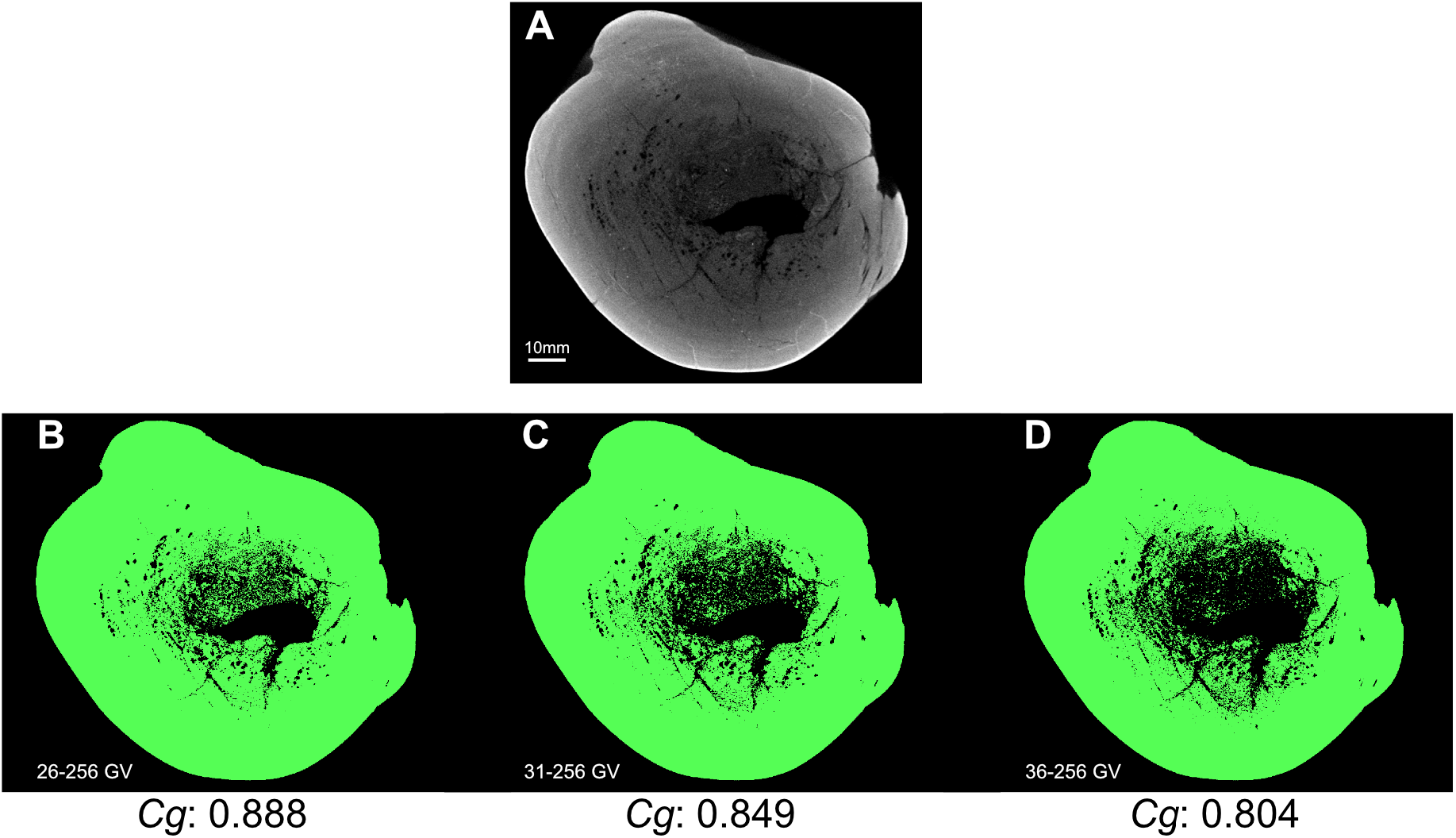
Impact of threshold choice on *Cg* in a cross section of the femoral shaft in a second subadult specimen of cf. *Spinosaurus aegyptiacus* (CMN 41869). (A) CT section from the proximal end of the shaft. (B–D) Section images and corresponding *Cg* values after processing with gray-value (GV) lower thresholds ranging from 26 to 36 GV on a 256 GV gradient. Threshold values determine which pixels are regarded as bone versus non-bone; higher thresholds yield lower *Cg* values.

Given the results from three optional thresholds (Fig 7B–D) for the cross section of the second femur of *Spinosaurus*, we selected the middle image with a *Cg* of 0.849 as the best binary visualization because it registers the less-dense cancellous bone near the medullary cavity without also obliterating what appears to be vascular canals in adjacent cortex on the left and lower sides of the medullary cavity. This *Cg* value is very close to the mean value (0.847) from our thresholding range. In this case, there is no physical thin section to examine under magnification in polarized light to verify what is bone or mineralized infill.

Although this CT-based femoral section (Fig 7A) was not available to Fabbri *et al.*, they later reported its *Cg* as 0.914 [28] without presenting the binarized image they used for *Cg* measurement. Presumably they employed more extreme thresholding than the maximum we considered reasonable (Fig 7B). Extreme thresholding would raise *Cg* by obliterating some of the smaller spaces in the binarized section. In this case, the *Cg* measurements on the same section differ by 7% due to procedures used in preparation of binarized images.

Nonetheless, it is clear from available specimens of *Spinosaurus aegyptiacus* that some individuals nearing maturity maintained a reduced medullary cavity with a femoral-shaft *Cg* under 0.900. We reported accurately on this variable condition of medullary cavities in the long bones and their presence in certain vertebral centra in *Spinosaurus* [27]:

> A second femur of Spinosaurus2 (Fig. 1a, b), which is nearly identical in size to the infilled neotypic femur3 in their study (Fig. 1c), has a significant medullary cavity lined with cancellous bone that would register as significantly less dense as a thin section at mid shaft. Medullary cavities are also variably present in forelimb bones of Spinosaurus (Fig. 1d) resembling those in the long bones of Suchomimus, a fully “terrestrial” spinosaurid by their account. Fabbri et al.1:ED, Fig. 10 state that Spinosaurus and Baryonyx “possess dense, compact bone throughout the postcranial skeleton,” yet all three have pneumatic spaces in their cervical column4 that exceed in volume the variable long bone infilling, as well as large medullary cavities hollowing the centra at the base of the tail. Neither of these features are present in any secondarily aquatic vertebrate divers that employ bone density as ballast.

Commenting on this new information on variability, Fabbri *et al.* introduced several errors [28]:

> Myhrvold et al.1 state that a single phalanx of the neotype of Spinosaurus possess a medullary cavity, invalidating our inference of widespread osteosclerosis across the postcranium of this animal; we show here that a cross section of the phalanx lacks any medullary cavity, as previously described in Ibrahim et al.13–14

and later:

> Caudal vertebrae 1 and 4 of the neotype of Spinosaurus: contrary to what suggested by Myhrvold et al.1, no pneumatization is present in the caudal region of this taxon.

We clearly described the *variable* presence of the medullary cavity in both fore and hind limb long bones in *Spinosaurus*, figuring the medullary cavity along the length of a manual phalanx from an adult individual as opposed to the subadult neotype (Fig 3D). We were aware of the infilled manual phalanges of the neotype. The image they republished of this infilled shaft condition was taken by one of us (PCS in [22]) from a break in the proximal shaft of a proximal manual phalanx, not at midshaft as they indicated [28: Fig 1c]. Medullary cavities are variably present in CT scans of a broader sampling of manual phalanges referable to *Spinosaurus aegyptiacus* from the Kem Kem Group. The centra of anterior caudal vertebrae in *Spinosaurus* and other spinosaurids, likewise, have a capacious medullary space that hollows the interior of the centrum, as we reported [26]. Contrary to Fabbri *et al.* [28], no one has claimed that the hollowed anterior caudal centra in various spinosaurids are *pneumatic*.

We present here CT cross sections from a third femur of *Spinosaurus aegyptiacus* from a very young individual collected in the same beds in Morocco as the first two (Fig 8E). This femur, which measures only 11.8 cm in length [104], has a large medullary cavity extending along the length of its shaft and would pertain to an individual with a body length of approximately ∼2.0 meters. Ontogenetic infilling of the medullary cavity does not appear to have been initiated, with a midshaft *Cg* of approximately 0.695 (Fig 9).

**Fig 8.**
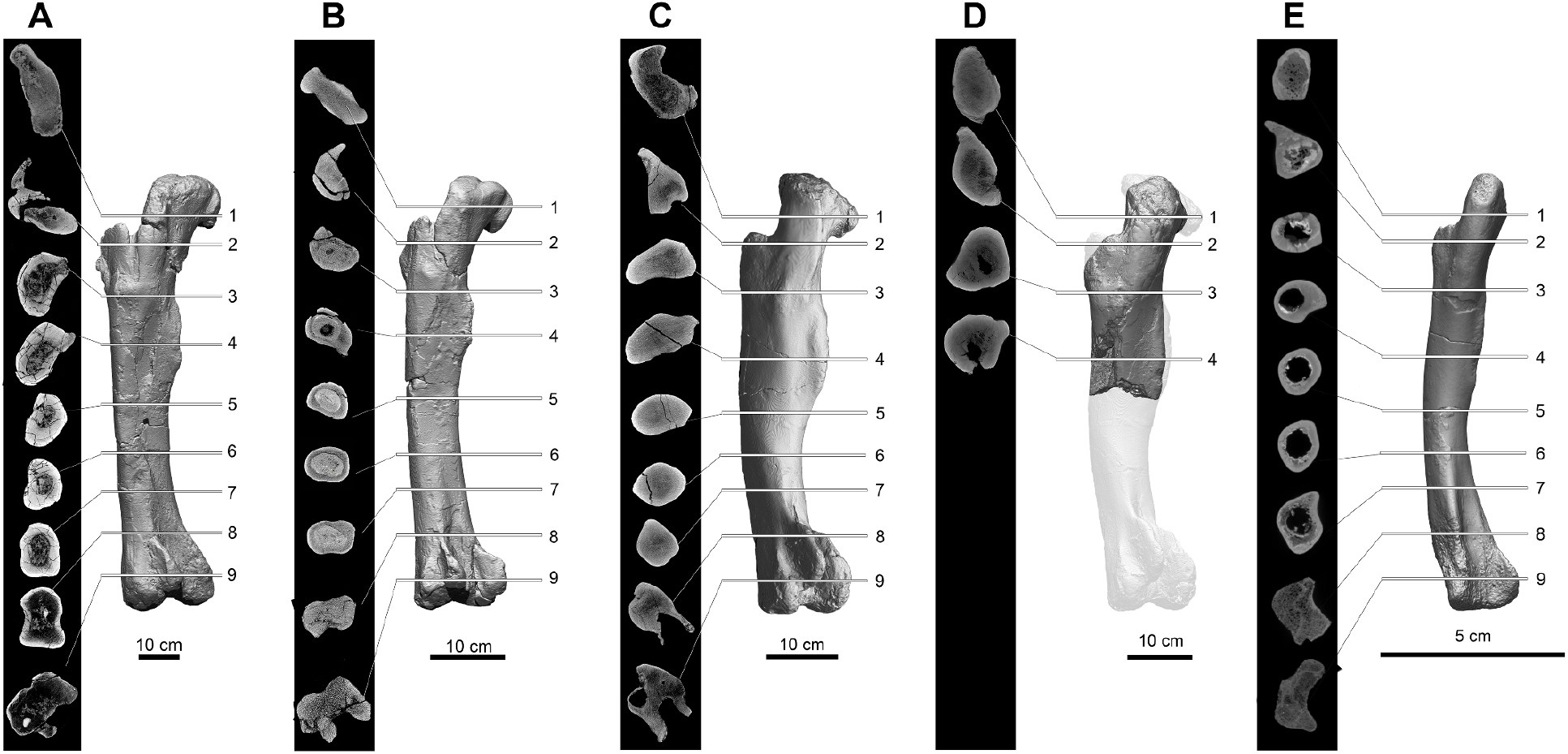
CT cross sections of the femoral shaft in two spinosaurids adjusted to the same side (left) and length. (A) *Suchomimus tenerensis*, adult (holotype), length 107.5 cm (MNBH GAD500). (B) *Suchomimus tenerensis*, juvenile, length 54.6 cm (MNBH GAD72, reversed). (C) *Spinosaurus aegyptiacus*, subadult (holotype), length 61.0 cm (MNBH GAD500). (D) *Spinosaurus aegyptiacus*, subadult, estimated length 61.0 cm (CMN 41869, reversed), (E) *Spinosaurus aegyptiacus*, juvenile, length 11.8 cm (CMN 50382, reversed).

**Fig 9.**
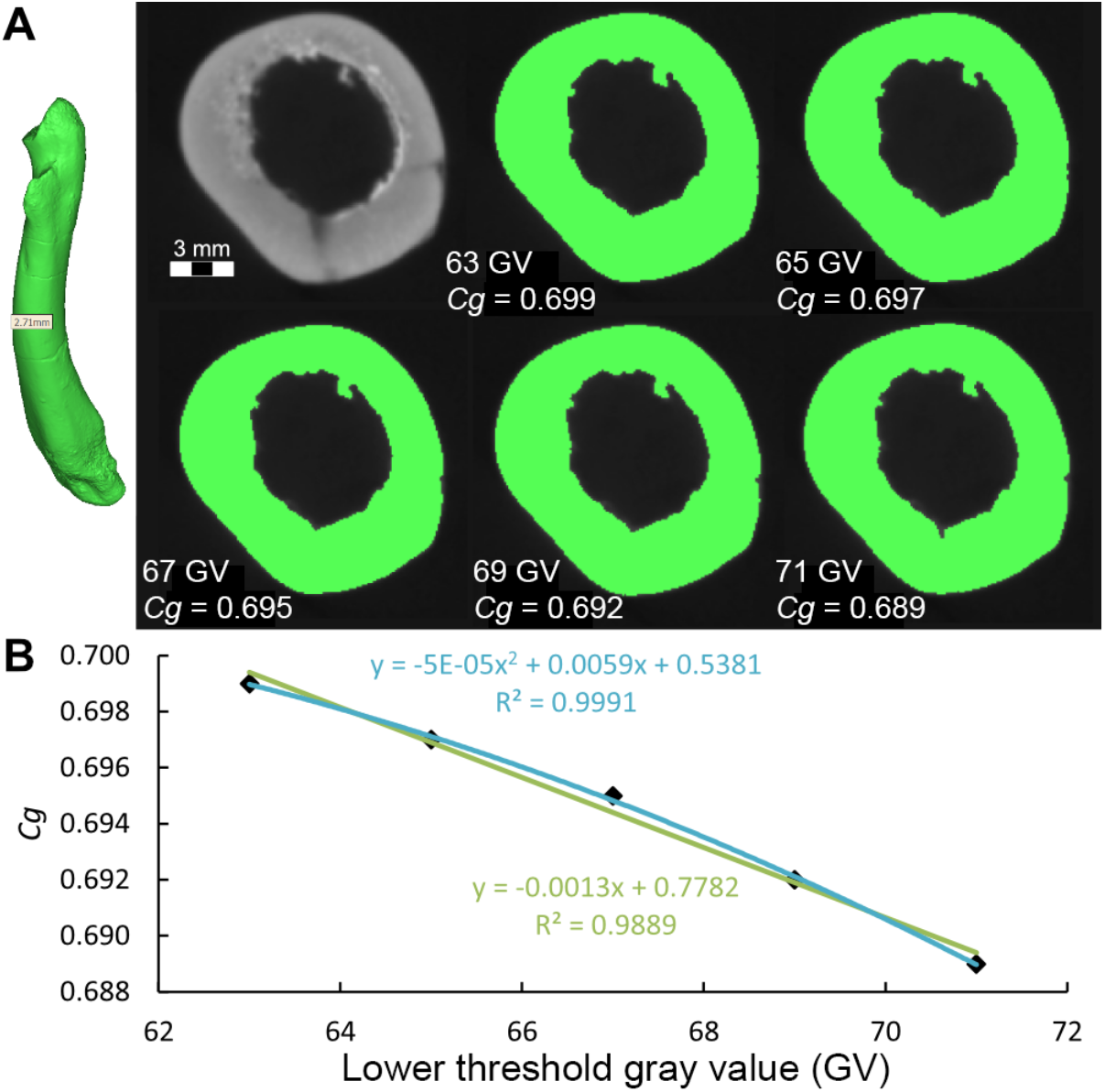
Bone compactness derived from CT scan of a juvenile femur of cf. *Spinosaurus aegyptiacus* (CMN 5038). (A) 3-D rendering and midshaft cross section generated from a CT scan with binarized images (green) differing in their lower threshold gray-value setting. (B) Plot showing linear change of about 10% in *Cg* over threshold range.

#### Baryonyx walkeri

Only the distal one-third of the right femur of the holotype is preserved [108]. There is crushing inward of anterior and posterior intercondylar areas, leaving only a small section of the shaft available for estimating bone compactness. This portion of the shaft was CT-scanned. Fabbri *et al.* used three closely spaced cross sections across ∼6 cm of the shaft to generate three estimates of *Cg* ranging from 0.826 to 0.876 (relative variance of 5.8%). The two most complete sections generated the minimum and maximum *Cg* values [18: Suppl. information, Fig 3e, f]. For the section generating the maximum value, the cracks had been infilled with solid bone and used for *Baryonyx* in their femoral dataset [18: Fig 1b].

We attempted to replicate their *Cg* estimate of 0.876 for *Baryonyx* using the CT scan they published [18: Suppl. Fig 3e]. We prepared three CT sections across ∼2 cm of shaft (Fig 10) in the region of their preferred section. We also infilled the cracks with solid bone density. We prepared two options for removal of matrix from the medullary cavity, each binarized with three different threshold values. The first option attempted to replicate the exact shape of the medullary cavity they defined and removed [Fig 10A–10C, top row of each panel]. As a second option, we examined the CT section and made an independent evaluation of the limits of fossilized bone, adjusting the medullary cavity boundary outward to include more material that did not show bone texture [Fig 10A–10C, bottom row of each panel]. We see no positive evidence in the scan for cancellous bone in this band.

**Fig 10:**
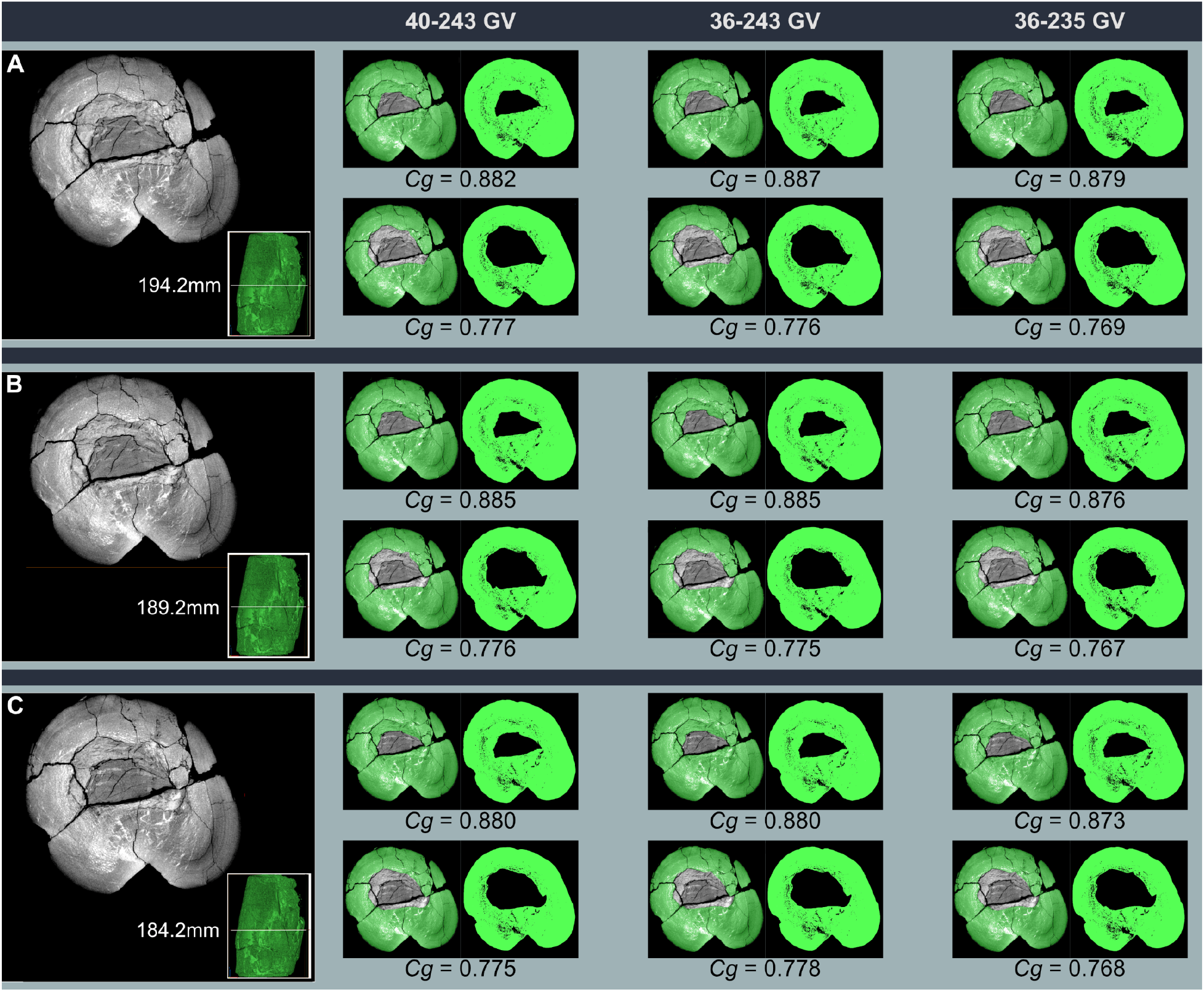
C*g* measured in three adjacent CT cross sections through the distal shaft of the right femur of the spinosaurid *Baryonyx walkeri* (NHMUK 9951). For each of three CT sections (A– C, posterior aspect of femur oriented toward bottom) taken in successively in more distal positions across 1 cm on the distal shaft of the femur in the portion of the shaft used by Fabbri *et al.* [18: Suppl. Fig 3e] for their best estimate of *Cg*. The small inset view shows distal end of the femur in medial view with mm distance from the bottom of the radiograph provided by Fabbri *et al.* To the right are three gray-value (GV) thresholds (left to right; 40–243, 36–243, 36–235) capturing a reasonable range of values that might be selected by researchers to binarize the radiograph. For each threshold, masking of the matrix infilling of the medullary space is shown in transparent (left) and binarized (right) views. Option 1 (top row) attempts to replicate medullary masking as published by Fabbri *et al*. Option 2 (bottom row) eliminates additional medullary material that we confirmed from the CT scan as matrix infill rather than cancellous medullary bone. Fabbri *et al.* reported a *Cg* of 0.876. The *Cg* range for our three slices in the vicinity of their preferred CT section using their masking is 0.773–0.887 (mean 0.830); their *Cg* measure is near the high end of that range. The *Cg* range with our masking is 0.767–0.778 (mean 0.773); their *Cg* measure is well above that range.

When we replicated their medullary cavity masking, their reported measure of 0.876 fell into the high end of the range of *Cg* we obtained for our three sections (0.773–0.877). When we chose our own (slightly larger) masking for the medullary cavity, the range of values obtained (0.767–0.778) excludes their higher value for *Baryonyx* (*Cg* = 0.876). Our mean *Cg* value for the distal shaft of *Baryonyx* (0.773) remains higher than the value reported by Fabbri *et al.* for *Suchomimus* (*Cg* = 0.682), but that value seems artificially low, as discussed in the next section. What seems clear at this point is that *Baryonyx*, like *Suchomimus*, retained an average-sized medullary cavity for a large theropod, the distal shaft of which generates a *Cg* less than 0.800.

#### Suchomimus tenerensis

Fabbri *et al.* figured two magnified thin sections for *Suchomimus tenerensis* identified as “G51” and “G94,” which are field numbers for the holotype (MNBH GAD500) and a referred subadult individual (MNBH GAD70), respectively [18: Suppl. Fig 2d, e]. Neither of these specimens were sectioned, however, and MBNH GAD70 does not preserve more the proximal end of one femur. We do not believe these thin sections pertain to *Suchomimus*.

One of us (PCS) made a four-part thin section from the distal end of an adult femur of *Suchomimus tenerensis* (Fig 11), which has a length (107.5 cm) and distal condylar width (23 cm) identical to that of the holotype. The position of the section on the distal shaft is similar to that taken in *Baryonyx*. An image of this thin section was refigured as a binarized image by Fabbri *et al.* [18: Fig 3d], who reported a *Cg* of 0.682 (Fig 11E). We commented, after reexamining the bone and original thin section, that there was additional cancellous bone not shown in their binarized image that likely lowered their reported *Cg* value [27].

**Fig 11.**
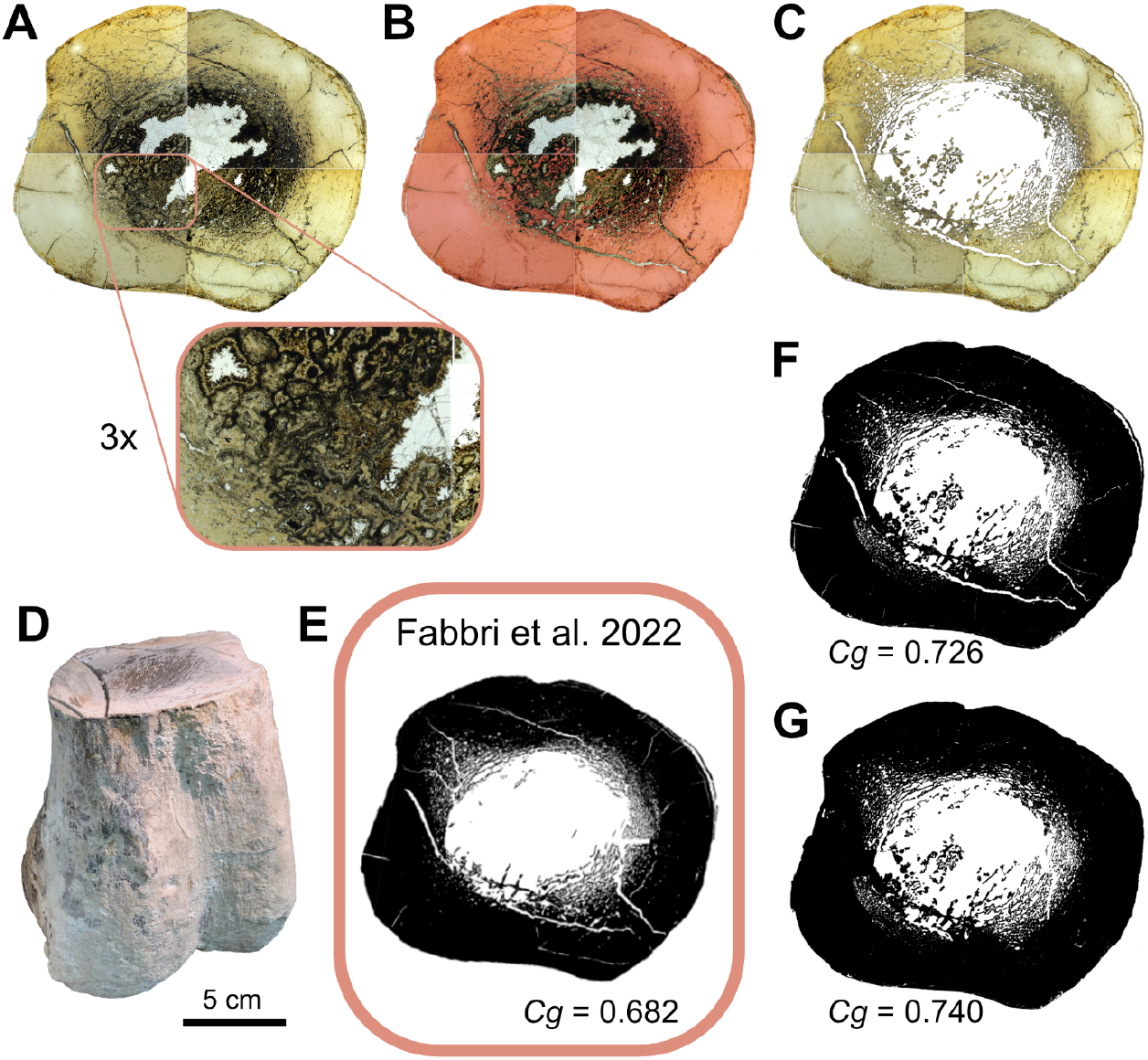
Cg derived from ta thin section from the distal femoral shaft of an adult specimen of *Suchomimus tenerensis* (MNBH GAD99). (A) Composite image of the four-part thin section with an enlargement showing the complex relation between cancellous bone and dark-stained mineral infilling. (B) Cancellous bone (red) adjacent to dark-stained matrix in the core of the femoral shaft. (C) Digital removal of matrix adjacent to cancellous bone. (D) Distal femur showing position of thin section. (E) *Cg* and binarized image from Fabbri *et al.* (F) *Cg* from binarized image after digital removal of matrix adjacent to cancellous bone. (G) Final *Cg* after filling in matrix-filled cracks.

In response, they introduced misinformation without examining either the thin section or host bone [28]:

> Myhrvold et al.1 suggest that we underestimated bone density in Suchomimus during the conversion of the femoral thin section into a black & white figure (the curating step prior to estimation of bone compactness), causing us to mis-identify bone as rock matrix. However, we did not apply our techniques blindly, but instead used careful observation to quantify bone compactness. As shown in Figure 1, the bone tissue in this specimen has a distinct white hue: Myhrvold et al.1 conflate the mineral infilling surrounding the trabecular bone and bone tissue.

Due to color variation, many thin sections including those examined here cannot be properly evaluated without examining them under a stereoscope or at least accessing magnified views of the thin section. In this case, a magnified view of the section clearly shows differentially distributed cancellous bone invading the medullary cavity, especially in the lower two thin-section quadrants (Fig 11A), contrary to Fabbri *et al.* [28]. We differentiated cancellous bone from adjacent dark-stained mineral deposits under stereoscopic magnification of the thin section (Fig 11B). After removal of mineral deposits and binarizing the image, a *Cg* of 0.726 was obtained, which is 6% greater than that reported by Fabbri et al. (Fig 11C, 11E, and 11F). We made that measurement to be fully comparable to Fabbri *et al.* without repair of matrix-filled cracks, which also effectively lower *Cg*. When those cracks are repaired, the final best estimate of the *Cg* of this specimen of *Suchomimus* is 0.740 (Fig 11G), approximately 8% higher than reported by Fabbri *et al*. and only 4% less than our best estimate of *Cg* in *Baryonyx*. Distal femoral shaft sections in *Suchomimus* appear to have *Cg* greater than 0.700.

We also took a thin section from the midshaft of a femur from a juvenile *Suchomimus tenerensis*, with femur length approximately half that of the adult, which shows a relatively large medullary cavity (Fig 8B, S3 Fig). *Cg* in the juvenile would be quite low and increase considerably during growth to adult body size.

### Use of phylogenetic flexible discriminant analysis

#### Methodological origin

Fabbri *et al.* have employed a relatively new statistical procedure, phylogenetic flexible discriminant analysis (pFDA), to reach their conclusions regarding the identification of habitual behaviors in extinct tetrapods. pFDA, a phylogenetic adaptation of flexible discriminant analysis (FDA), was first applied to study nocturnality in dinosaurs via statistical analysis of eye and scleral ring shape [29,30]. FDA, in turn, was generalized by Hastie *et al.* [109] from Fisher’s much earlier linear discriminant analysis (LDA) [110].

Fisher created LDA to separate point clouds that follow multivariate normal distributions. Later work has shown that LDA is closely related to ANOVA and regression techniques. It represents an extension that creates a discriminant or decision boundary, a line that that can be used to estimate which data points more likely to belong to two or more multivariate normal distributions. LDA has been used with bone-compactness data to create discriminants between groups [6,29,30,99,111] without incorporating phylogenetic data in the analysis.

The general form of the probability density function for a bivariate normal distribution is given by Equation (1), where *x* is a two-dimensional position vector, *μ* is a two-dimensional position of the centroid of the distribution, Σ is a 2 × 2 covariance matrix, and |Σ| is its determinant, and the superscript ^T^ denotes matrix transpose.

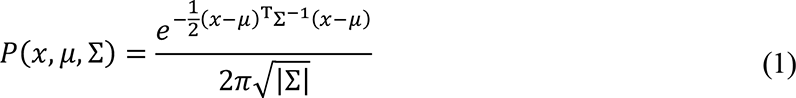

In the case of 2-class or binary classification, LDA assumes that there is a different distribution for each class, with centroids *μ = μ_1_, μ_2_* that are distinct (*μ_1_ ≠ μ_2_)*, but that both distributions have the same covariance matrix Σ. Mathematically, this assumption ensures that the decision boundary is a line.

LDA classifies points by computing the Mahalanobis distance from a test point to the centroid of two or more reference groups, using the pooled, within-group covariance matrix. The squared Mahalanobis distance appears in an argument to the exponential function in Equation (1). In the case of a distribution with unit variance and a covariance matrix that is the identity matrix, *i.e.*, 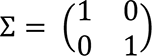, it reduces to the Euclidean distance.

In LDA and FDA, a fundamental assumption is that a test point can be classified by assigning it to the group that has the smallest Mahalanobis distance between the point and the group centroids 𝜇_1_, 𝜇_2_ (*i.e.*, the multidimensional means of the classes). The locus of points equidistant between group centroids corresponds to the decision boundary. Hastie *et al.* [109] generalized LDA to FDA by including nonlinear decision boundaries, as well as a Bayesian approach to integrate prior probabilities with a penalized Mahalanobis-distance metric, to create a system for scoring test points.

Motani and Schmitz [95] introduced pFDA as a specific instance of FDA in which a phylogenetic-bias correction is done in a similar fashion to PGLS using branch lengths from phylogenetic trees that cover the taxa in the analysis to determine phylogenetic correlation between taxa under an evolutionary model, such as Brownian motion. In principle, FDA could allow the use of nonlinear decision boundaries, but pFDA as implemented by Motani and Schmitz (and thus by Fabbri *et al.*) uses linear boundaries, thereby assuming that both groups have the same covariance matrix as in Equation (1). pFDA is thus a phylogenetic version of LDA.

The procedure advocated by Motani and Schmitz [95] is to use only extant taxa that have well-constrained phylogenies and branch lengths for the training set. They do not recommend including extinct taxa. To cope with the phylogenetic uncertainty, Fabbri *et al.* created a set of 100 random trees, each having its own phylogenetic covariance matrix. The matrices are sequentially passed to code that uses them to perform FDA. Each such trial results in a classification probability for each test taxon and for each random tree. A point is considered classified if, among all 100 trees, the median probability of its belonging to a given class exceeds 50%.

We note in passing that use of the median is not justified; by construction, the random trees are equally likely models of past evolution. Using medians to eliminate extreme values is therefore unjustified. Further analysis than that reported by Fabbri *et al.* would be required to determine how accurately a sample of only 100 branch-length trials characterizes possible topological and branch-length errors in the tree and how this would impact the analysis.

In principle, the phylogenetic signal could have a strong effect. In practice, Fabbri *et al.* find very little evidence of phylogenetic signal in their dataset, with Pagel’s λ parameter taking values 0.02 ≤ λ ≤ 0.07 across the various datasets and trials. This is consistent with other studies of *Cg* with comparable datasets [38,41] that seek to analyze convergent features across many clades. As a result, one would expect little difference between these results and those obtained with ordinary LDA. In view of that and the uncertainty in the tree for extinct taxa, we question whether this dataset is worth analyzing with a phylogenetic method.

To illustrate the properties of LDA and pFDA, we consider a special case of Equation (1) for two distributions having the properties given in Equation (2), where the covariance matrix Σ is identical for both distributions and is a multiple *σ*^2^/2 of the 2 × 2 identity matrix.

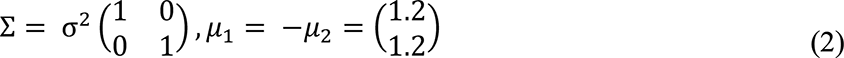

The centroids of the two distributions, *μ*_1_ and *μ*_2_, are reflected in the origin across the line *y* = −*x*. We can take these two groups of points and use them as the training dataset. It is easily shown that the optimal decision boundary must be the perpendicular bisector of the line between the centroids *μ*_1_ and *μ*_2_, which in this case is given by the line *y* = −*x*. We show a plot of 1000 points drawn from each of two distributions that follow Equation (2), denoted group 1 and group 2 (Fig 12). In both cases, *σ* = 0.55, which plays a role in these bivariate distributions that is very similar to the parameter in a conventional univariate normal distribution. The distance between the centroid of either distribution and the decision boundary is *d* = 1.7 = 3.1*σ*. As a result, the concentration of points matches what one would expect of a univariate normal distribution: most of the points are concentrated near the centroid and thus appear on the same side of the decision boundary as the centroid. Such points would be correctly classified by the decision boundary.

**Fig 12.**
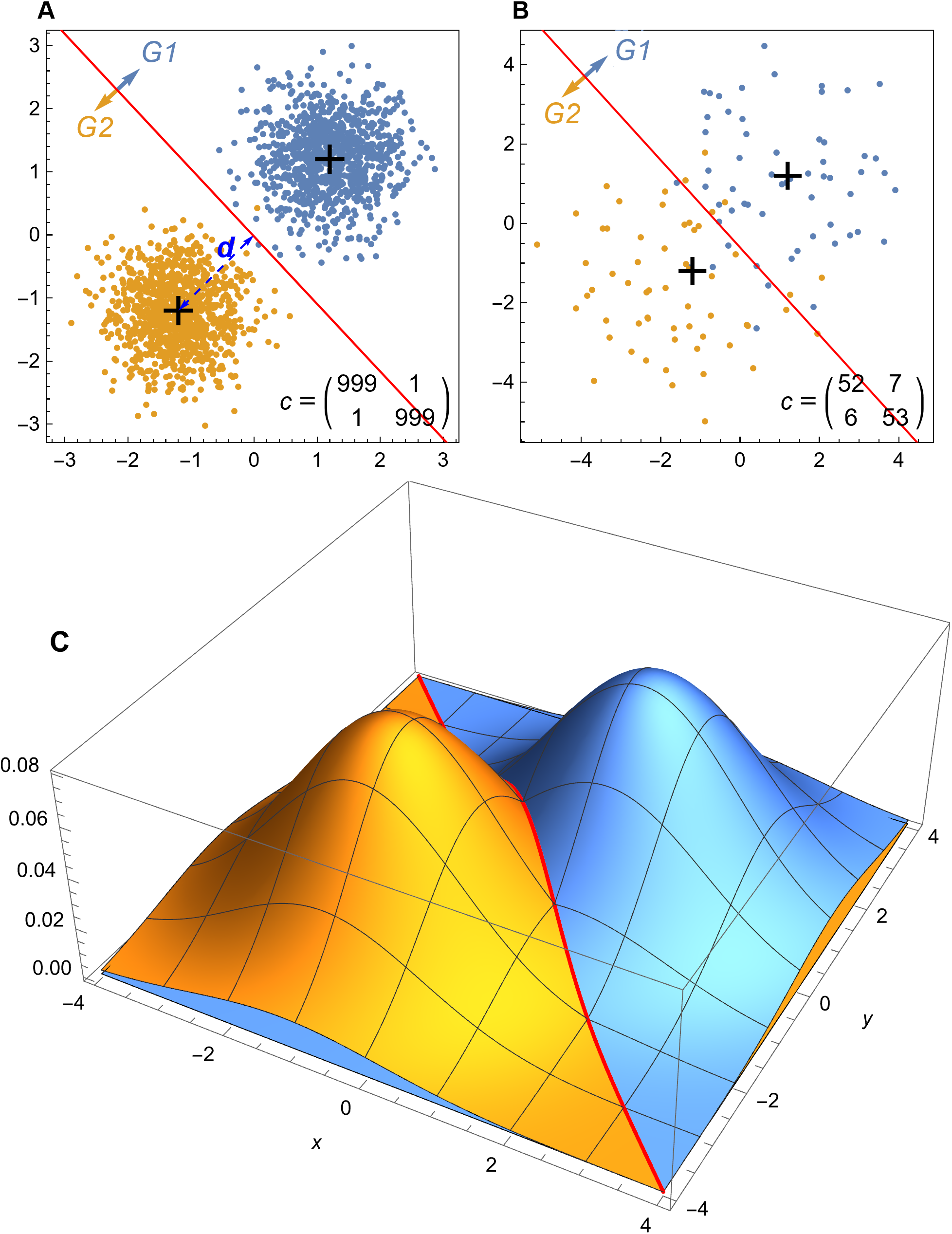
Simulated data plots for LDA methods. (A) 1000 pseudorandom points drawn from each of two multivariate normal distributions given by Equations (1) and (2) and *σ* = 0.55 are plotted, with points from each distribution colored according to the legend. The decision boundary for LDA is given by the red line; points above the line are classified as group 1, points below the line are classified as group 2. Note that one point from group 1 lies on the other side of the decision boundary and is incorrectly classified as group 2. One point from group 2 is similarly misclassified. The centroid of each distribution is denoted by the black cross mark, the distance from the centroid to the decision boundary *d* is denoted by a dashed blue line. The confusion matrix *c* (Equation (4) in S1 Appendix) is shown. (B) 59 points from distributions with the same centroids as (A) but with *σ* = 1.414. The higher value of *σ* leads to a larger number of points being misclassified. (C) The underlying probability density functions for the same distributions as in (B). The distributions of blue and gold points are equal at the red decision boundary line *y* = −*x*. Abbreviations: G1, group 1; G2, group 2.

Points that fall on the opposite side of the decision boundary are considered *misclassified*. Because these points are part of the training dataset, they would be termed training-data errors [112]. Because the points are highly concentrated and the decision boundary is relatively far in terms of *σ*, there are only a few of these points in the random sample shown. Fig 12B shows an example with the same distribution centroids, but this time with 59 points in each group, and *σ* = 1.414, which means that *d* = 1.2*σ*. The shorter distance in terms of *σ* greatly increase the number of training set errors.

The fundamental idea behind LDA is shown in a plot of the probability density functions for the multivariate normal distributions given by Equations (1) and (2) (Fig 12C) with the same *σ* = 1.414 (Fig 12B). The two normal distributions meet where they cross each other. This is a line in space which falls along the line *y* = −*x* when projected onto the (*x*, *y*) plane. That is the decision boundary. Along that line there is an equal probability from either probability density function, so one cannot say which it belongs to. At other points, the probability is higher of belonging to one point or another.

One can calculate the exact probability that a point will lie on the wrong side of the decision boundary by integrating the probability density function over the half plane defined by the wrong side of the decision boundary to yield Equation (3), where erfc() is the error function and *d* is the distance from the distribution centroid to the decision boundary.

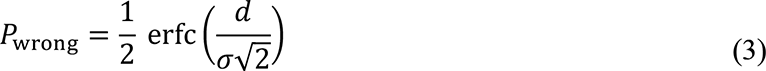

This relation matches the familiar case of the marginal distribution of points in a normal distribution, expressed in terms of the standard-deviation-adjusted distance. Thus, we expect from Equation (3) that 68.27% of the points would be misclassified if *d* = *σ*, 2.5% of the points to be misclassified if *d* = 1.96 *σ*, and 1% if *d* = 2.33 *σ*, following usual rules of thumb.

In the example shown in Fig 12B, *P*_wrong_ = 0.115, so we can expect about 11.5% of points in each group will be misclassified in a very large sample. The section above on variation in *Cg* shows data suggesting (very roughly) that the median variation in *Cg* is 18% or a variation of 0.18 on the scale of *Cg*. If we assume that the *Cg* errors are normally distributed, then effect on this on classification error is 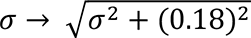, which could be a large effect, depending on the size of 𝜎 without error and the distance between centroids *d*.

For the distributions in Fig 12A, *P*_wrong_ = 0.00955, which is just below 1 in 1000. As a result, we expect zero to a few errors in a training set of 1000 points. That is what we see in the example plotted in Fig 12A: one error misclassifying a group 1 point as group 2, and one error misclassifying group 2 as group 1. As the number of trials increases, the number of incorrect points converges toward *n* × *P*_wrong_, with some statistical variation.

Equation (3) reveals an important principle: even when we use synthetic data drawn from multivariate normal distributions, classification via LDA or FDA can *never be error-free*. That follows from the simple fact that the domain of the multivariate normal distribution ranges across the interval (−∞, ∞) in each independent variable, whereas the distance from the distribution centroids to the decision boundary is finite. Therefore there can always be valid points from one distribution that lie on the other side of any decision boundary—not as an outlier (which implies an erroneous point) but rather as an entirely valid data point that LDA will misclassify. Note that this effect does *not* depend on the sample size. As the number of data points in the training set grows to infinity, the error converges to Equation (3).

#### When should classification be believed?

All practitioners of statistical analysis face a common challenge. What probability of incorrect results due to random effects is acceptable? In normal statistical analysis, all results are accompanied by estimates of their statistical quality, such as the *p* value, confidence level, confidence interval, or other quantitative error estimates. Unfortunately, pFDA is relatively novel and does not natively produce a formal *p* value, confidence interval, or other metric of random effects—nor can we find any in prior literature. Instead, there are two primary sources of classification within pFDA: posterior probabilities and empirical classification performance on known cases.

An invocation of a pFDA classifier returns a list of values for each of the test taxa to be classified. These values can be denoted *P*_2_—the posterior probability that the point belongs to the class with categorical variable *D = 2*. A minor complication is that, in the application of pFDA by Fabbri *et al.*, each test point is classified for 100 random phylogenetic trees, so the result for a single taxon is typically a list of *P*_2_ values of length 100. The criterion that Fabbri *et al.* use is to classify the taxon as *D =* 2 if the median value of the *P*_2_ list is greater than 0.5. Fabbri *et al.* discussed this criterion [18: 859]:

> “We summarised our results by providing the median value of those 100 posterior probabilities and the number of times a particular taxon is predicted as subaqueous forager (median probability of 50% or more). This gives us two proxies of the likelihood of each taxon to be an actual subaqueous forager. For instance, a taxon could be predicted 100 times as subaqueous forager with a median probability of 51% which means the evidence for this extinct species to be an actual subaqueous forager is very weak and this inference has to be considered very unlikely. Median probabilities need to be within the range of 80–100% to be considered as strong evidence of subaqueous forager.”

Fabbri *et al.* clearly recognized the weakness in the criterion, as a value of *P*_2_ only marginally greater than 0.5 is weak evidence indeed. Since there are two classes, a classification probability of 0.5 is equivalent to a random guess, such as flipping a coin. Normally, a result that is only infinitesimally better than random would be accorded little probative value. Despite their clear recognition of the weakness inherent in this approach, the criterion used for classification remains *P*_2_ > 0.5 rather than 1.0 ≥ *P*_2_ ≥ 0.8, as discussed in the passage.

If *P*_2_ was an absolute probability, then *P*_2_ = 1.0 would indicate absolute certainty, and there could be no possibility of misclassification. But this interpretation is incorrect. *P*_2_ is *not* an absolute probability—instead it is a classification score which at best is *a possibly erroneous estimate* of the *relative* probability of being in one class versus the alternative, conditioned on the assumption that the classes are multivariate normal distributions.

Indeed, the predicted probability *P*_2_ can, and typically does, erroneously classify at least some of the points in the training dataset. Fabbri *et al.* mention classification performance only in this passage [18: 856]:

> The correct classification rates of our phylogenetically flexible discriminant analyses ranges are 84–85% (femora) and 83–84% (ribs) (Figs. 2, 3, Supplementary Materials, Supplementary Tables 7–10). This increases to 90% in both datasets when excluding graviportal and deep diving taxa (Figs. 2, 3, Supplementary Tables 7–10).

The Supplementary Tables 7–10 they cite in the passage do not define precisely what is meant by “correct classification rates.” Significant ambiguity exists because there are multiple classification performance metrics (see S1 Appendix, section 4). The referenced tables contain only posterior probability predictions *P*_2_ for the dinosaur test taxa, including the spinosaurids, rather than taxa of known class in the training datasets, so they cannot be used as a basis for a correct classification rate. The most likely source of these “correct classification rates” is that they are training-set classification errors.

If the method has “84–85%” correct classification on its training set, then the erroneous classifications must be 15% to 16%. If the method fails 15% of the time, then it is performing at least three times worse than the usual threshold for random errors in statistical methods, which is 5%. Such a result would normally be considered not statistically significant.

Adding an unfortunate complication, we discovered a flaw in the pFDA code that systematically misstates the confusion matrix from which classification performance is measured (S1 Appendix, section 5). Our replication attempts produce classification rates slightly different from those reported by Fabbri *et al.* This issue may be why they do not match exactly.

Although it may seem intuitively obvious that increasing the *P*_2_ classification threshold would offer better evidence, the situation is actually quite complex. Increasing the classification threshold does make for a more stringent criterion, but it also means that a higher percentage of the training dataset will be misclassified (S1 Appendix, section 3).

If the goal is to achieve a result that is of statistical quality comparable to normal statistical significance, we would ask for no more than a 5% chance of an error due to random effects. Naively this would imply that the classification errors in the training set ought to be less than 5% and also that the posterior probability *P*_2_ > 0.95. Both criteria should be met with 95% confidence. Ideally, pFDA would produce a formal *p* value, confidence interval, or other mechanism to estimate the total statistical error in the classification, but unfortunately the method lacks this critical mathematical feature. Until it does, requiring both a low classification-error rate in the training set and a high posterior probability seems prudent, but is not mathematically rigorous. Currently there is no mathematically rigorous way to perform pFDA with quantitative assessment of the impact of statistical errors on the results.

#### Effects of sample size on classification

A tacit assumption in Fabbri *et al.*, as in most other statistical analyses in biology, is that biological factors underlying the dataset classes produce a true statistical distribution of the variates. The pFDA method assumes that the true distribution for each class conform to a multivariate normal distribution, or a close approximation thereof. But the parameters of those true distributions are *unknown*. The pFDA method must estimate the parameters from the finite sample in the training dataset. This situation is common to virtually all statistical analyses but strangely seems to have been overlooked in the literature on pFDA. It has also been largely overlooked in biological applications of LDA, save for a few examples [113].

The sample size is a key element determining the statistical power and precision of a statistical analysis because it controls the how well the finite sample approximates the underlying biological distribution. Neither Fabbri *et al.* nor any other pFDA study of which we are aware offers any analysis of how the size of the training dataset affects classification accuracy.

Fig 13 presents the results of Monte Carlo simulations of an LDA classifier that explore sample-size effects for the case of symmetric multivariate distributions of the form given by Equations (1) and (2). The accuracy of binary classification has been long studied, and many different mathematical metrics have been developed to measure it, as discussed further in S1 Appendix, section 4.

**Fig 13.**
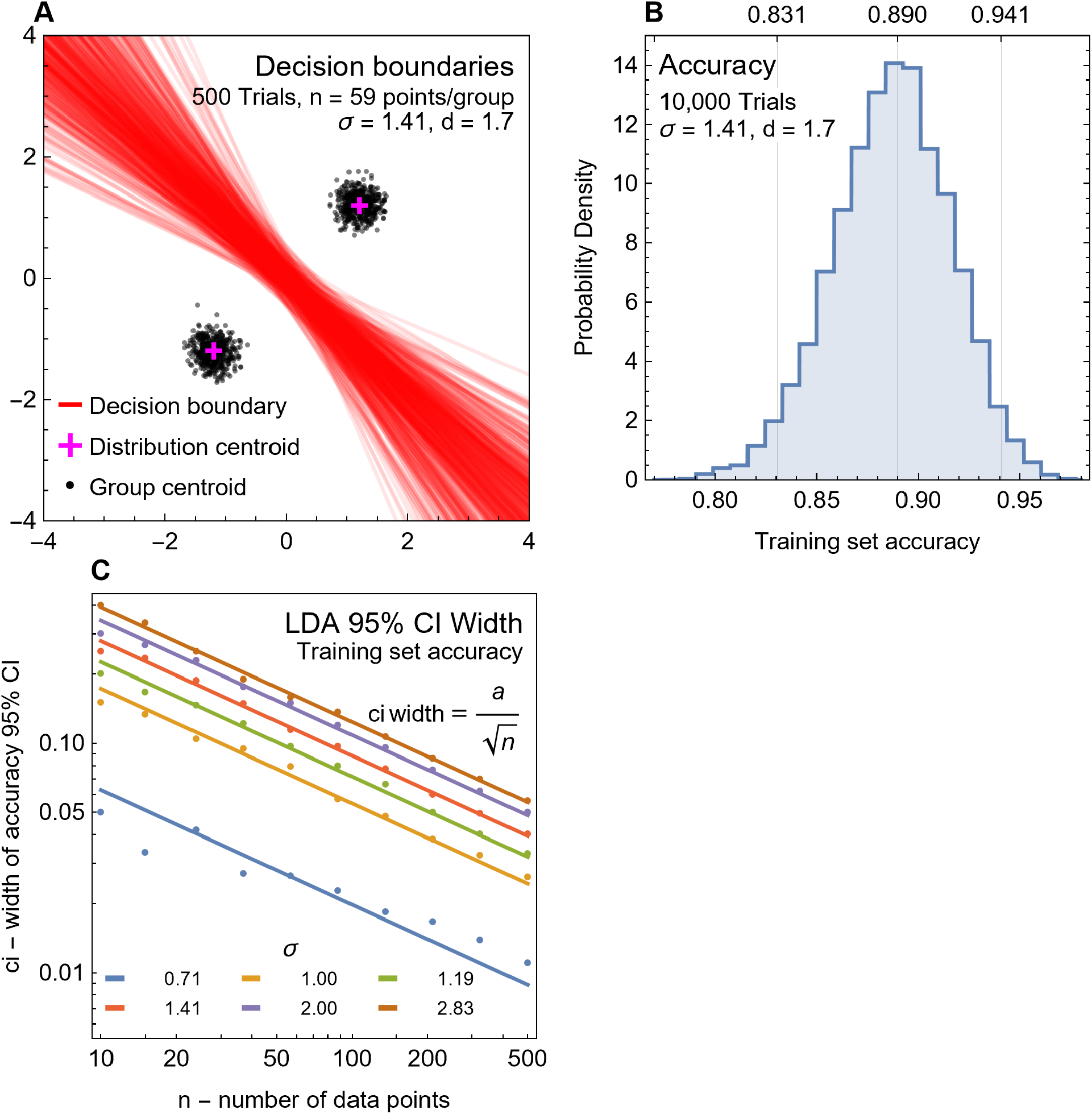
Sample-size effects in LDA/FDA. (A) Decision boundaries and centroids of point groups for 500 trials of 59 points drawn from a multivariate normal distribution of Equations (1) and (2) with specified values of *σ* and *d*. The centroids of the distributions are shown by the magenta crosses; empirical centroids of each group of 59 points are black dots. The decision boundaries are red lines. Groups of 59 points are insufficient to accurately estimate the distribution centroid. The estimation error leads to variations in both the empirical-group centroids and the decision boundaries. (B) A histogram of the training-dataset classification accuracy is shown for 10,000 trials with the same parameters as (A). The theoretical accuracy for the values of 𝜎 and 𝑑 is 0.885, but the 95% confidence interval extends from 0.831 to 0.941, a width of 10%. (C) Monte Carlo simulations of classification accuracy for point groups with *d* = 1.7 and varying values of *σ* and *n* points per group. Lines show an empirically derived relationship for the width of the 95% confidence interval in classification accuracy: CI width = *a*/n^1/2^, where is *a* is a fitting constant determined for each value of *σ*. Abbreviations: CI, confidence interval.

There is noticeable scatter among the empirical centroids derived from groups of 59 pseudorandom data points (Fig 13A). The empirical centroids only roughly approximate the true centroids of the distributions from which they were drawn. The decision boundaries from these groups also show considerable scatter in both midpoint and slope. Assessing the classification accuracy of 10,000 trials of two groups of 59 points yields a histogram, which peaks at the theoretical classification accuracy of 0.885 with considerable scatter (Fig 13B). With a 95% confidence interval spanning 0.831 to 0.941, there is 10% accuracy.

Repeating this 10,000-run Monte Carlo experiment for multiple points per group 10 ≤ *n* ≤ 500 and for values of the standard-error parameter 0.707 ≤ *σ* ≤ 2.83, we find that the width of the 95% confidence interval on classification accuracy closely follows an empirically derived relation (Fig 13C). The general behavior is that the width of the confidence interval scales proportionately to 1/*n*^1/2^ for sample size *n*, as is typical for the normal distribution. The sample size effect is thus of critical importance to the application of FDA or LDA classification because it greatly lowers expected classification performance at small points per group counts.

A consequence of the relationship shown in Fig 13C is that, depending on the values of *σ* and *d*, the classification performance of LDA might not meet the criterion *a* = 0.05 for significance (see Equation (6) of S1 Appendix, section 4), unless there are hundreds of points in each class. However, for cases of low classification error (Fig 13C), statistical significance might be achieved at fairly low point counts.

Conversely for *d*/*σ* < 1.64, Equation (3) tells us that even an infinite number of data points will not achieve statistical significance. This pattern is an example of the “ecological fallacy,” a common error in statistical inference. Briefly stated, one generally cannot accurately classify a point by comparing it to its statistical distribution, or the average and variance derived from the distribution; specific cases may work, but only if the variance of the distributions is sufficiently small. The ecological fallacy is discussed further in S1 Appendix, section 1.

These results pertain to classification accuracy of the training dataset, but a similar phenomenon occurs for any metric of classification performance. Classification accuracy is linear in the confusion-matrix components, whereas some metrics, such as *MCC*, are nonlinear in the components (S1 Appendix, section 4). The exact form of the relation between 95% CI width and the distribution parameters will thus change, but we expect the overall behavior to be qualitatively similar. In actual practice, we do not know the exact distribution and instead have only the finite sample to work with. Another complication is that pFDA, as used in Fabbri *et al.*, has an additional source of random variation due to the creation of randomly generated phylogenetic trees.

Here we take the Fabbri *et al.* datasets and apply a bootstrap approach [114] to estimate the finite sample-size effects on pFDA (see Methods and materials). In essence, a straightforward bootstrap creates a training dataset for each trial by randomly sampling the existing training dataset (with replacement). The data is then processed with the Fabbri *et al.* R script and associated datasets for pFDA, including the creation of 100 random trees for each trial. Each random tree creates its own decision boundary.

Fig 14A presents results for 100 trials and 10,000 random trees. Each decision boundary has a corresponding classification of the points, yielding a confusion matrix, which can then be converted into classification accuracy for the training dataset. The distribution of accuracy values is plotted as a histogram (Fig 14B). The bootstrap samples also affect the posterior classification probabilities *P*_2_ (Fig 14C). These results are qualitatively what one would expect, given the result on the simplified synthetic dataset (Fig 13). The effect of small sample size leads to scatter in both the group centroids and the decision boundaries.

**Fig 14.**
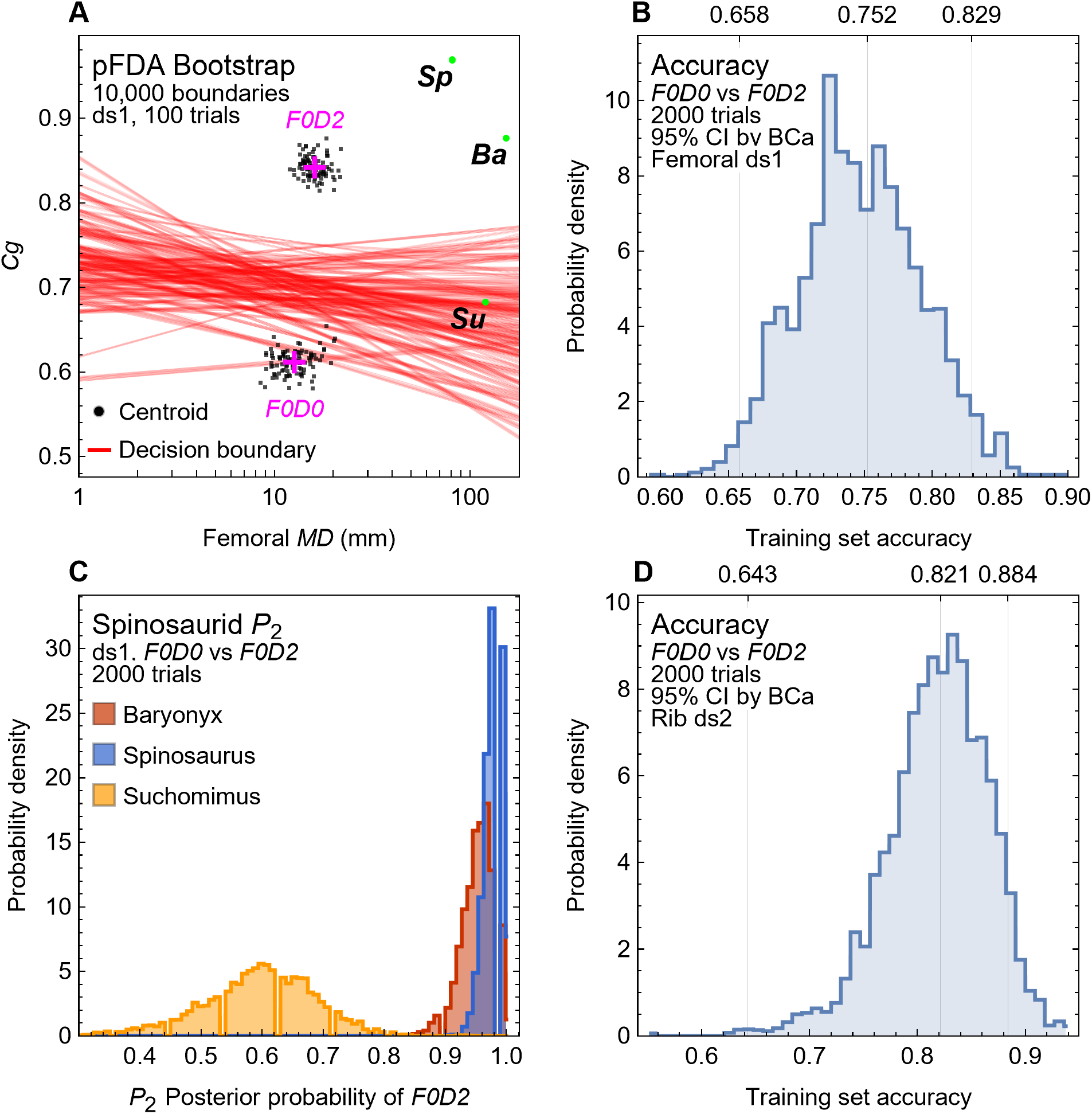
Sample size effects for pFDA with data from Fabbri *et al.* [18]. (A) Decision boundaries (red lines) and point-group centroids (black dots) for 100 trials created using a bootstrap method described in the text, operating on the *F0D0* and *F0D2* subsets of the Fabbri *et al.* femoral dataset. Each bootstrap trial draws 100 trees at random, each with its own decision boundary, as generated by the Fabbri *et al.* R script. As in Fig 13, considerable scatter is evident in both the centroid positions and decision boundaries. Data points for *Spinosaurus*, *Baryonyx*, and *Suchomimus* are plotted in green. The downward slopes of most of the decision boundaries, as well as the leftward offset of the centroids of *F0D0* subset versus that for *F0D2*, show the effect of lower *MD* for *F0D0*. (B) A histogram of classification accuracy of the training dataset is shown for 2000 trials of a parametric bootstrap as in (D). The median training set classification accuracy is 0.752, and the 95% confidence interval is 0.658 to 0.829, a width of 0.171. (C) Histograms of 𝑃_$_, the posterior probability of belonging to group *D = 2*, for Spinosaurid taxa across 2000 trials of the same dataset as (A) and (B). (D) Histogram of training-set classification accuracy similar to (B), but for rib data. The median training-set classification accuracy is 0.821, and the 95% confidence interval is 0.643 to 0.884, a width of 0.241. Abbreviations: *Sp*, *Spinosaurus*; *Ba*, *Baryonyx*; *Su*, *Suchomimus*; CI, confidence interval; BCa, bias-corrected-and-accelerated method.

The 95% confidence interval can be estimated using bootstrap methods. We performed 2000 bootstrap trials for each dataset and tabulated the training dataset errors. The bias-corrected-and-accelerated (BCa) method was used to assure good results on the confidence interval [114]. Because each case also has 100 random trees, 200,000 results were used for the creation of the confidence intervals (Table 7). The training-set error rate is widely considered to be overoptimistic, and our use of it is thus very conservative.

**Table 7.** Bootstrap-estimated 95% confidence intervals for training-set classification performance metrics.

| Dataset | Metric | *F0D0* vs. *F0D2* |  |  | Many vs. *F0D2* |  |  |
| --- | --- | --- | --- | --- | --- | --- | --- |
|  |  | Median | 95% CI |  | Median | 95% CI |  |
| ds1 | *A* | 0.752 | 0.658 | 0.829 | 0.826 | 0.777 | 0.899 |
|  | *B* | 0.751 | 0.658 | 0.831 | 0.821 | 0.794 | 0.904 |
|  | *MCC* | 0.539 | 0.392 | 0.710 | 0.623 | 0.553 | 0.788 |
| ds2 | *A* | 0.821 | 0.643 | 0.884 | 0.809 | 0.642 | 0.846 |
|  | *B* | 0.819 | 0.656 | 0.878 | 0.799 | 0.657 | 0.843 |
|  | *MCC* | 0.637 | 0.328 | 0.764 | 0.573 | 0.309 | 0.664 |
| ds3 | *A* | 0.794 | 0.618 | 0.863 | 0.894 | 0.789 | 0.938 |
|  | *B* | 0.794 | 0.608 | 0.863 | 0.891 | 0.763 | 0.928 |
|  | *MCC* | 0.615 | 0.258 | 0.730 | 0.764 | 0.548 | 0.855 |
| ds4 | *A* | 0.913 | 0.739 | 0.924 | 0.89 | 0.713 | 0.926 |
|  | *B* | 0.895 | 0.720 | 0.909 | 0.868 | 0.691 | 0.917 |
|  | *MCC* | 0.812 | 0.440 | 0.838 | 0.715 | 0.349 | 0.808 |

To determine how well taxa in the training dataset are correctly classified by pFDA, we measured three classification performance metrics: *A*, *B*, and *MCC* (described in S1 Appendix, section 4). In each case, 2000 bootstrap trials, each with 100 random trees for a total of 200,000 data points, were gathered for *A*, *B*, and *MCC*. The BCa bootstrap method was used to construct accurate 95% confidence intervals for both the method of Fabbri *et al.* [18] (columns “Many vs. *F0D2*”), which retain taxa with $F = 0, 1, 2$ and $D = 0, 1$ as the alternative to the *F0D2* group, and also for the better-supported *F0D0* vs. *F0D2* approach. The lower bound on the accuracy metrics formed by the minimum of the 95% CI shows that the training-data classification with these datasets has errors ranging from approximately 20% to 40%—higher in some cases. Abbreviations: CI, confidence interval.

The primary effect of the sample size is that the 95% confidence interval is much broader than the point estimates. The importance to the interpretation of pFDA classification results is that they are even more uncertain than one would expect from simply evaluating the training set error from a single run. Consider dataset ds1: using the method of Fabbri *et al.* (Table 7 columns “Many vs. *F0D2*”), the median value of the accuracy metric *A* (Equation (5) in S1 Appendix, section 5) is 0.826, which is roughly consistent with the claim in that the correct classification rate is “84–85% (femora)” [18]. However, the 95% CI for this value ranges from 77.7% to 89.9%. If one uses the better-supported method of only comparing the *F0D0* and *F0D2* subsets, the median drops slightly to 75.2% error, and the lower bound of the 95% CI is 65.8%. This implies that at 95% confidence, the classification error on training data could be as high as 33%, more than six times the random-error threshold of 5% typical for statistical studies.

As noted above, Fabbri *et al.* removed “pelagic” and “graviportal” taxa from data sets ds1 and ds2 to yield the ds3 and ds4 data sets. The stated purpose was to improve classification rates, and one can see that for *F0D0* vs. *F0D2*, the median values for accuracy *A* do improve between ds1 (0.752) and ds3 (0.794). However, in practical terms this improvement is undermined by the greater uncertainty in the smaller ds3 dataset; the lower bounds of the 95% CI are 0.658 and 0.618 for ds1 and ds3, respectively. Thus, at the 95% confidence level, accuracy *decreased* with the smaller ds3 dataset. This effect does not occur for the Many vs. *F0D2* approach because it contains far more taxa.

If we wish to use a similar error threshold to that typically used for *p* values and confidence levels, then the conventional choice of *a* = 0.05 would, in terms of the performance metrics *A*, *B*, and *MCC* (Equations (5) and (6) in S1 Appendix, section 4), be translated into the criteria *A* > 0.95, *B* > 0.95, and *MCC* > 0.9. More theoretical work would have to be done to create metrics analogous to *p* values or confidence intervals. Nevertheless, these crude metrics express the basic premise that a 5% chance of random error is acceptable. None of the 95% intervals in Table 7 would meet that usual criteria for statistical evidence that supports a scientific conclusion, as the lower bound on their classification errors are well below the criteria above.

A similar effect occurs with *P*_2,_ the posterior probability of membership in the *F0D2* category. Rather than being a single value, *P*_2_ becomes a distribution of values (Fig 14). We can use BCa bootstrap to create 95% confidence intervals from this distribution.

In order to assess the impact of variations in the data for the spinosaurid test taxa, we tested the data points used by Fabbri *et al.*, as well as hypothetical modifications to them (Table 8). We are not proposing that these modified values are necessarily more correct or believable; our intention is to perform a sensitivity analysis to see how *P*_2_ for a taxon is affected by changes in its test data point.

**Table 8.** Alternative test-data points for spinosaurids.

| Dataset | Taxon | MD | Cg | MD source | Cg source |
| --- | --- | --- | --- | --- | --- |
| **ds1**<br>**(Femur)** | *Baryonyx 0* | 154 | 0.876 | ref. [18] | ref. [18] |
|  | *Baryonyx 1* | 154 | 0.887 | ref. [18] | Fig 10, high |
|  | *Baryonyx 2* | 154 | 0.826 | ref. [18] | Fig 10, median |
|  | *Baryonyx 3* | 154 | 0.767 | ref. [18] | Fig 10, low |
|  | *Spinosaurus 0* | 81.52 | 0.968 | ref. [18] | ref. [18] |
|  | *Spinosaurus 1* | 81.52 | 0.804 | ref. [18] | Fig 7, low |
|  | *Spinosaurus 2* | 81.52 | 0.849 | ref. [18] | Fig 7, med |
|  | *Spinosaurus 3* | 81.52 | 0.888 | ref. [18] | Fig 7, high |
|  | *Spinosaurus 4* | 81.52 | 0.914 | ref. [18] | ref. [28] |
|  | *Spinosaurus 5* | 133.434 | 0.968 | Scaled by 1.64 | ref. [18] |
|  | *Spinosaurus 6* | 133.434 | 0.804 | Scaled by 1.64 | Fig 7, low |
|  | *Spinosaurus 7* | 133.434 | 0.849 | Scaled by 1.64 | Fig 7, med |
|  | *Spinosaurus 8* | 133.434 | 0.888 | Scaled by 1.64 | Fig 7, high |
|  | *Spinosaurus 9* | 133.434 | 0.914 | Scaled by 1.64 | ref. [28] |
|  | *Suchomimus 0* | 120.6 | 0.682 | ref. [18] | ref. [18] |
|  | *Suchomimus 1* | 120.6 | 0.628 | ref. [18] | Scaled by 0.92 |
|  | *Suchomimus 2* | 146.4 | 0.682 | Actual max GAD500 | ref. [18] |
|  | *Suchomimus 3* | 146.4 | 0.628 | Actual min GAD500 | Scaled by 0.92 |
| **ds2**<br>**(Rib)** | *Baryonyx 0* | 42.2 | 0.921 | ref. [18] | ref. [18] |
|  | *Baryonyx 1* | 42.2 | 0.8289 | ref. [18] | Scaled by 0.9 |
|  | *Spinosaurus 0* | 35.10 | 0.931 | ref. [18] | ref. [18] |
|  | *Spinosaurus 1* | 35.10 | 0.838 | ref. [18] | Scaled by 0.9 |
|  | *Spinosaurus 2* | 57.45 | 0.931 | Scaled by 1.64 | ref. [18] |
|  | *Spinosaurus 3* | 57.45 | 0.838 | Scaled by 1.64 | Scaled by 0.9 |

The variations cover three principal approaches. The first covers the maximum diameter *MD*. The *Spinosaurus* neotype has been estimated at 72% of full size. On allometric grounds, one would expect that *MD* would therefore scale by a factor of 1.64 = (1/0.72)^1.5^. This factor follows from the assumption that body mass scales as the cube of linear size (*i.e.,* isometrically), while *MD* scales as the square root of load. This scaling is not relevant for the other spinosaurids in the analysis, many of which are subadults short of maximum size but not juveniles. However our scanning of an adult *Suchomimus* femur MNBH GAD500 (S3 Fig) did reveal a quite different maximum diameter than that reported by Fabbri *et al.*, so we use our value as a variation.

The values of *Cg* are also varied based on our attempts to replicate the measurements of Fabbri *et al.* using CT scans, as discussed above and shown in Figs 7 and 10. In the case of the rib data, we did not have alternative measurements and instead considered a hypothetical scaling of *Cg* by 0.9, equivalent to a 10% reduction.

The results of the *P*_2_ confidence intervals are shown in Table 9. The first broad conclusion is that finite-size effects have a strong impact on *P*_2_ for the original test points of Fabbri *et al.* The point *Baryonyx 0* in ds1 has a median of 0.97 in the Many vs. *F0D2* case, which seems to be strong evidence. But finite-size effects mean that to 95% confidence its lower bound is 0.94, slightly below the 95% threshold discussed above. Using the better-supported *F0D0* vs. *F0D2* method, the lower bound drops to 0.84. Qualitatively similar results occur for the *Spinosaurus 0* and *Suchomimus 0* data points in both ds1 and ds2.

**Table 9.** 95% confidence intervals for posterior probability prediction *P*_2_ for spinosaurids.

| Dataset | Variant | *F0D0* vs. *F0D2* |  |  | Many vs. *F0D2* |  |  |
| --- | --- | --- | --- | --- | --- | --- | --- |
|  |  | Median | 95% CI |  | Median | 95% CI |  |
| **ds1**<br>(Femur) | *Baryonyx 0* | 0.96 | 0.84 | 0.98 | 0.97 | 0.94 | 1.00 |
|  | *Baryonyx 1* | 0.97 | 0.85 | 0.98 | 0.97 | 0.90 | 0.99 |
|  | *Baryonyx 2* | 0.93 | 0.75 | 0.95 | 0.91 | 0.75 | 0.95 |
|  | *Baryonyx 3* | 0.85 | 0.59 | 0.90 | 0.74 | 0.46 | 0.85 |
|  | *Spinosaurus 0* | 0.98 | 0.90 | 0.99 | 0.99 | 0.98 | 1.00 |
|  | *Spinosaurus 1* | 0.88 | 0.71 | 0.90 | 0.83 | 0.66 | 0.88 |
|  | *Spinosaurus 2* | 0.93 | 0.79 | 0.95 | 0.92 | 0.82 | 0.95 |
|  | *Spinosaurus 3* | 0.96 | 0.85 | 0.97 | 0.96 | 0.89 | 0.98 |
|  | *Spinosaurus 4* | 0.97 | 0.86 | 0.97 | 0.98 | 0.95 | 0.99 |
|  | *Spinosaurus 5* | 0.99 | 0.95 | 1.00 | 0.99 | 0.98 | 1.00 |
|  | *Spinosaurus 6* | 0.89 | 0.68 | 0.92 | 0.83 | 0.59 | 0.89 |
|  | *Spinosaurus 7* | 0.94 | 0.79 | 0.96 | 0.92 | 0.80 | 0.96 |
|  | *Spinosaurus 8* | 0.96 | 0.83 | 0.97 | 0.96 | 0.89 | 0.98 |
|  | *Spinosaurus 9* | 0.97 | 0.86 | 0.98 | 0.98 | 0.96 | 1.00 |
|  | *Suchomimus 0* | 0.61 | 0.35 | 0.73 | 0.32 | 0.12 | 0.47 |
|  | *Suchomimus 1* | 0.43 | 0.24 | 0.63 | 0.13 | 0.04 | 0.25 |
|  | *Suchomimus 2* | 0.62 | 0.34 | 0.75 | 0.32 | 0.12 | 0.50 |
|  | *Suchomimus 3* | 0.44 | 0.24 | 0.66 | 0.13 | 0.04 | 0.27 |
| **ds2**<br>(Rib) | *Baryonyx 0* | 1.00 | 0.96 | 1.00 | 0.95 | 0.86 | 0.98 |
|  | *Baryonyx 1* | 0.97 | 0.81 | 0.99 | 0.85 | 0.67 | 0.92 |
|  | *Spinosaurus 0* | 0.99 | 0.92 | 1.00 | 0.94 | 0.81 | 0.98 |
|  | *Spinosaurus 1* | 0.96 | 0.79 | 0.98 | 0.84 | 0.69 | 0.91 |
|  | *Spinosaurus 2* | 1.00 | 0.96 | 1.00 | 0.95 | 0.82 | 0.98 |
|  | *Spinosaurus 3* | 0.97 | 0.80 | 0.99 | 0.86 | 0.64 | 0.93 |

The sensitivity analysis using hypothetical points shows that *P*_2_ is highly dependent on data point values. In the case of *Baryonyx*, the lower bound on *P*_2_ from 0.84 (*F0D0* vs. *F0D2*) drops to 0.75 for the median value found in Fig 7 to 0.59 for the low value from Fig 10 (*Baryonyx 3*, ds1), and even lower to 0.46 in the Many vs. *F0D2* case. Similar effects are seen in the ds2 (rib) datasets: *Baryonyx 0* has a lower bound of 0.95 for *F0D0* vs. *F0D2*, but this drops to 0.81 in *Baryonyx 1*, which differs only by the *Cg* scaled by 0.9. Since the median percent difference in *Cg* found for multiple specimens of the same taxon (Tables 5 and 6) is 18.6%, this 10% variation seems quite conservative. Yet it shifts the expected value of *P*_2_ from significant to dubious. Qualitatively similar results hold for the variations in *Spinosaurus* and *Suchomimus*. The only *Spinosaurus* data point that has a lower bound on *P*_2_ ≥ 0.95 is one of the variation data points: *Spinosaurus 5*, which is based on scaling *MD* to adult size and using the same *Cg* value as Fabbri *et al.* did.

The basic result of this section is that finite-size effects, which occur due to the relatively small number of datapoints relative to the variance in those datapoints, greatly reduce our confidence in the key parameters of training-set classification performance and posterior probability *P*_2_. None of the original datapoints for spinosaurids used by Fabbri *et al.* should be accorded much evidentiary weight. Our sensitivity analysis shows that even small variations in *Cg* (10% or less) can have a decisive effect on *P*_2_.

#### Verification of assumptions for pFDA

Statistical methods have validity only if they are applied to datasets that match the assumptions used in developing the method. Normal statistical practice is to test those assumptions, but Fabbri *et al.* do not report such tests. Here we perform several simple tests of the data.

As discussed above, the pFDA method is based on FDA and LDA, which were originally derived for multivariate normal distributions. However, it is clear upfront that the distributions of (log_10_(*MD*), *Cg*) points cannot closely follow a normal distribution in the *Cg* axis because normal distributions are defined on the open interval (−∞,∞), whereas *Cg* is restricted to the interval (0,1]. Note that this is true even after adjustment for phylogenetic bias; multiplication by a matrix with finite elements cannot make the *Cg* range become infinite.

One approach to testing the assumptions directly is to examine the discriminant values generated by the pFDA algorithm. This has two advantages. First, the discriminant values are directly used to calculate posterior probabilities, so the assumption of normality for them is quite important. Second, the discriminant values have already been corrected for phylogenetic bias correction, and the dimensionality has been reduced. The smoothed kernel distributions derived from the discriminant values are plotted in Fig 15.

**Fig 15.**
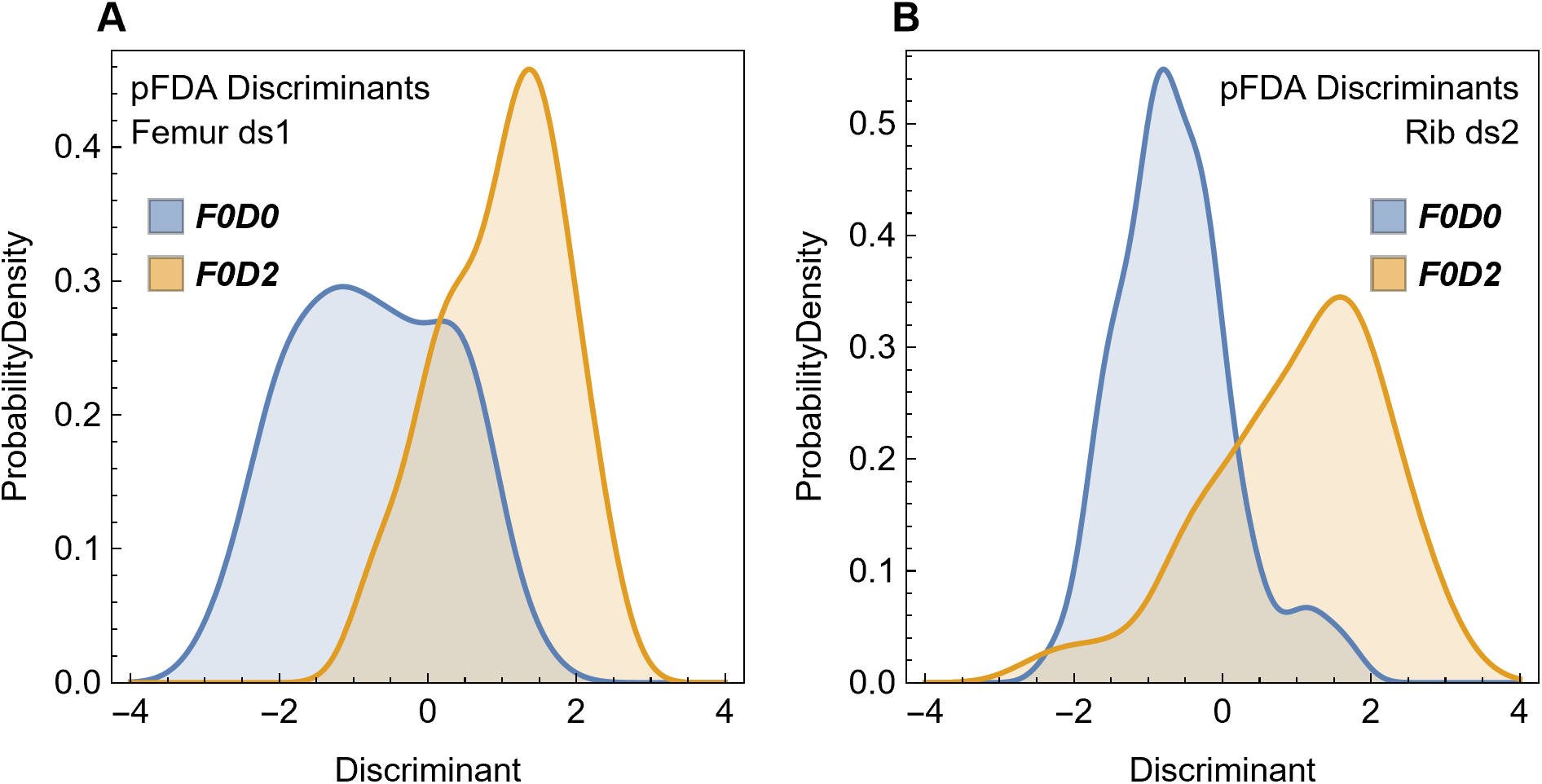
Smoothed kernel distributions for the pFDA discriminants from the (A) femur ds1 and (B) rib ds2 datasets of Fabbri *et al.* [18]. Distributions for the discriminants from both the *F0D0* and *F0D2* subsets of each dataset are plotted. There is considerable overlap between the *F0D0* and *F0D2* groups for both femoral and rib datasets. None of these distributions appear to be normal distributions; statistical tests confirm that they are not.

As seen in Fig 15, the discriminants do *not* closely follow a normal distribution, nor does each pair of distributions appear to have equal variance. In addition, there is considerable overlap between the discriminants for two groups, indicating a high classification-error rate in the training sets. High overlap is demonstrated for the original datasets in Fig 1C and 1D, and via simple effect-size statistics (S5 Fig). Phylogenetic-bias correction does not eliminate the overlap between groups, which is to be expected given the very low values of Pagel’s λ found by Fabbri *et al*. The overlap between the *F0D0* and *F0D2* groups is a clear example of the ecological fallacy (S1 Appendix, section 1).

In order to assess the deviation from normality, we made maximum-likelihood estimates of the best-fitting distributions, including both standard distributions and mixtures of them. The parameters of the best-fitting distributions are shown in Table 10, along with their log-likelihood values, the information-theoretic fitting metrics BIC and AIC, and an overall fitting score, which is based on a Bayesian estimate that combines BIC, log-likelihood, and prior probabilities. These distributions are plotted in Fig 16.

**Fig 16.**
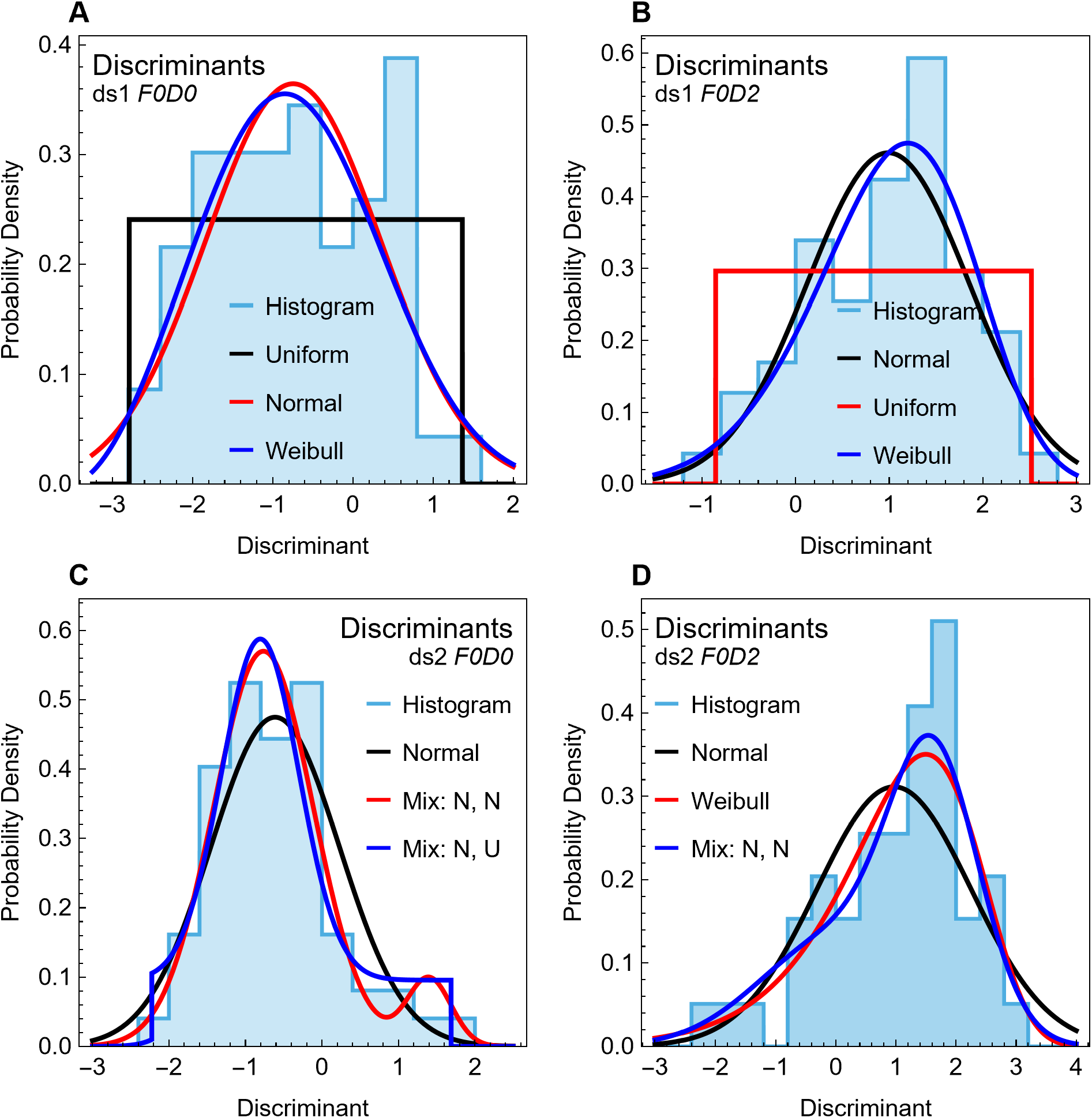
Histograms and fitted distributions for pFDA discriminants. (A, B) Histograms (light blue) of the discriminants for the *F0D0* and *F0D2* subsets of ds1, with fitted distributions (black, red, and blue curves). (C, D) Comparable plots for ds2. In each panel, the best-fitting distribution by Bayesian score is plotted in black, second best in red, and third in blue. Abbreviations: N, normal; U, uniform; Mix, a mixture of two distributions.

**Table 10.** Parameters and fitting metrics of best-fitting distributions to pFDA discriminants.

| Set | Dist. | Mix | Param1 | Param2 | Param3 | Var | BIC | AIC | Log-Like | Score |
| --- | --- | --- | --- | --- | --- | --- | --- | --- | --- | --- |
|  | Uniform | 1 | -2.79075 | 1.36502 |  | 2 | -2.92564 | -2.92172 | -1.4245 | -2.97255 |
| **ds1**<br>**F0D0** | Normal | 1 | -0.745417 | 1.09452 |  | 2 | -3.00533 | -3.00142 | -1.46435 | -3.02978 |
|  | Weibull | 1 | 2.81477 | 3.13253 | -3.52991 | 3 | -3.0111 | -3.00725 | -1.44807 | -3.13576 |
| **ds1**<br>**F0D2** | Normal | 1 | 0.98883 | 0.865144 |  | 2 | -2.52778 | -2.52329 | -1.22593 | -2.55182 |
|  | Uniform | 1 | -0.85619 | 2.52029 |  | 2 | -2.50959 | -2.5051 | -1.21684 | -2.55571 |
|  | Weibull | 1 | 6.0638 | 4.76879 | -3.43045 | 3 | -2.5206 | -2.51581 | -1.20336 | -2.64315 |
| **ds2**<br>**F0D0** | Normal | 1 | -0.612085 | 0.840423 |  | 2 | -2.41616 | -2.41011 | -1.17116 | -2.43903 |
|  | Mix: | 0.929453 | -0.764716 | 0.650705 |  | 5 | -2.40066 | -2.39462 | -1.10802 | -2.44692 |
|  | Normal,<br>Normal | 0.070547 | 1.39881 | 0.287749 |  |  |  |  |  |  |
|  | Mix: | 0.626313 | -0.806336 | 0.507906 |  | 5 | -2.37859 | -2.37254 | -1.09699 | -2.45469 |
|  | Normal,<br>Uniform | 0.373687 | -2.22421 | 1.68769 |  |  |  |  |  |  |
| **ds2**<br>**F0D2** | Normal | 1 | 0.968889 | 1.28094 |  | 2 | -3.26207 | -3.26519 | -1.58912 | -3.29101 |
|  | Weibull | 1 | 24.7765 | 26.0346 | -24.4948 | 3 | -3.19447 | -3.20205 | -1.53436 | -3.34204 |
|  | Mix: | 0.413881 | 0.010717 | 1.26865 |  | 5 | -3.28384 | -3.30682 | -1.53713 | -3.34238 |
|  | Normal,<br>Normal | 0.586119 | 1.64549 | 0.746214 |  |  |  |  |  |  |

Inspection of Table 10 shows that all the fitted distributions have strong support under AIC or BIC, differing only slightly in the various metrics. The Bayesian score indicates that a uniform distribution is the best fit for ds1 *F0D0*. The log-likelihood metric also would select the uniform distribution. The BIC and AIC metrics would select the fit to a Weibull distribution as best. In the case of ds1 *F0D2*, log-likelihood selects the Weibull distribution, while BIC, AIC, and the Bayesian score select the normal distribution.

In order to correctly predict class membership and calculate the posterior probabilities, pFDA requires both *F0D0* and *F0D2* datasets to be normal distributions. In the case of ds1, the best choices vary with the metrics, but no metric chooses the normal distribution as best for *F0D0*; for *F0D2* the best chosen vary between normal and Weibull. Under these model selection criteria, we find no statistical support for applying pFDA to the ds1 dataset.

The ds2 dataset is split; by Bayesian score, both *F0D0* and *F0D2* are best fit by normal distributions and thus qualify for pFDA. In contrast, the BIC, AIC, and maximum log-likelihood metrics would choose non-normal distributions, indicating that ds2 is also not suitable for pFDA.

Two issues produce these conflicting results. First, the datasets are inadequate, producing pFDA discriminants that are equivocal as to which distribution they support—there is very little variation in the actual metrics among the supposedly “best” choice and others.

The second issue is that each of the metrics measures a different aspect of fit. Although these metrics all have value, the most salient characteristic for their use in pFDA is whether the left tail of the *F0D0* distribution and the right tail of the *F0D2* distribution are accurate, because those are the portions of the distributions involved in the computation of the decision boundary and the assignment of posterior classification probabilities. This can be visualized from the overlap of the distributions in Fig 15.

A quantile-quantile plot allows direct comparison of the quantiles of the discriminant set with those of the fitted distributions (Fig 17). Significant deviation is observed in the right tail of *F0D0* versus a normal distribution (Fig 17A), in contrast to the much closer adherence by a uniform distribution (Fig 17B).

**Fig 17.**
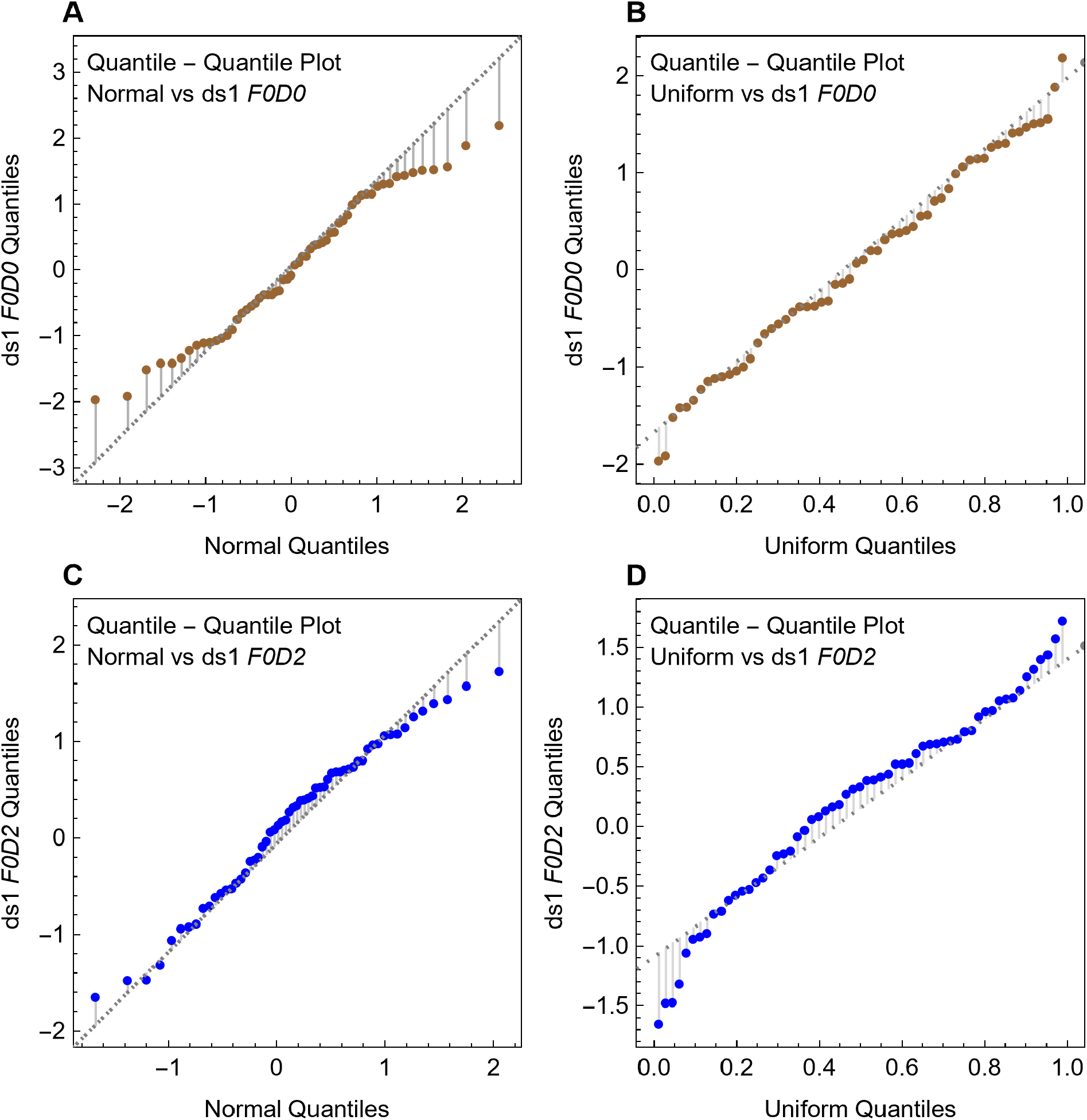
Quantile-Quantile plots of pFDA discriminants from dataset ds1 subsets *F0D0* and *F0D2*. Quantiles of normal (A, C) and uniform (B, D) distributions are shown as heavy brown (for *F0D0*) or blue (for *F0D2*) dots plotted against plot the quantiles of the discriminant distributions (thin dotted lines). (A) The pFDA discriminant for ds1 *F0D0* shows strong deviation from the normal distribution for the right tail of the distribution. (B) The deviation is much smaller for the uniform distribution. (C, D) Similar but smaller effects are observed for the left tails of the distributions for ds1 *F0D2*. Corresponding plots for the ds2 dataset are provided in S4 Fig.

A quantile-quantile plot for the ds2 dataset appears in S4 Fig. There we see that the right tail of ds2 *F0D0* strongly deviates from the normal distribution (S4A Fig), but closely matches the mixture of two normal distributions (S4B Fig). For ds2 *F0D2*, neither the normal nor Weibull distributions closely match the left tail.

The net result of both the distribution fitting and the quantile-quantile analysis is that there is little statistical confidence that the ds1 and ds2 datasets meet the assumption that they are normally distributed. The pFDA discriminants are not normally distributed under the most common model selection criteria, or at best are equivocal.

A second assumption required by pFDA is that the *F0D0* and *F0D2* subsets being compared have discriminants with the same variance. This prerequisite is fundamental to both LDA and the subset of FDA used by pFDA. Conventional variance equivalence tests can be used, if care is taken to choose those that are robust to deviations from a normal distribution, in light of the results above. Results of such tests show that neither the ds1 nor ds2 datasets meet the assumption of equal variances (Table 11).

**Table 11.** Results of variance-equivalence tests for pFDA discriminants.

| Test | ds1 *F0D0* vs. *F0D2* |  | ds2 *F0D0* vs. *F0D2* |  |
| --- | --- | --- | --- | --- |
|  | Statistic | *p* | Statistic | *p* |
| **Brown-Forsythe** | 5.29316 | 0.023212 | 7.62504 | 0.006756 |
| **Conover** | 2.33821 | 0.019377 | -2.67815 | 0.007403 |
| **Levene** | 5.18783 | 0.024593 | 9.70682 | 0.002347 |

Three tests of variance equivalence, an assumption of pFDA, were performed on the discriminants of the *F0D0* and *F0D2* groups being compared. These tests were chosen because they are considered robust to deviations from normality in the distributions. In both the ds1 and ds2 cases, the null hypothesis that the variances are equal is rejected with $p < 0.05$ by all three tests.

Linear discriminant analysis, as used by pFDA, is not appropriate where classes have unequal variances. Quadratic discriminant analysis (QDA) would be appropriate (assuming normal distributions in both classes) because the decision boundary between the datasets would be a quadratic curve (conic section). If LDA were applied to such a dataset, however, one would expect highly inaccurate classification because the straight-line assumption is violated [112]. As currently conceived, pFDA does not address phylogenetic QDA, but conceivably a pQDA could be developed.

Normal statistical practice in clustering or classification problems is to use the Hopkins statistic to assess whether the points have any genuine clustering [36,115–117]. The null hypothesis under this test is that the data points are distributed randomly in space. Failure to reject the null hypothesis implies that any apparent clusters are illusory and attributable to random chance. Here we apply the standard Hopkins statistic, as well as two variations by Lawson and Jurs [36] and Fernandez Pierna and Massart [35] (see Materials and methods), to the *F0D0* and *F0D2* subsets of both ds1 and ds2 (Table 12).

**Table 12.** Results (*p* values) from Hopkins statistic tests on datasets from Fabbri *et al.* [18].

| Dataset | Class subset | Hopkins *p* values |  |  |
| --- | --- | --- | --- | --- |
|  |  | Original | FPM | LJ |
| **ds1** | *F0D0* | 0.391 | 0.864 | 0.977 |
|  | *F0D2* | 0.984 | 0.238 | 0.005 |
| | *F0D0*: $\lambda = 0.06$ | 0.548 | 0.449 | 0.706 |
| | *F0D2*, $\lambda = 0.06$ | 0.821 | 0.580 | 0.587 |
| **ds2** | *F0D0* | 0.621 | 0.140 | 0.409 |
|  | *F0D2* | 0.145 | 0.193 | 0.602 |
| | *F0D0*: $\lambda = 0.07$ | 0.738 | 0.234 | 0.865 |
| | *F0D2*, $\lambda = 0.07$ | 0.432 | 0.231 | 0.542 |

Three versions of the Hopkins statistical test for clustering were applied to the *F0D0* and *F0D2* subsets of ds1 and ds2, for both the original datasets and phylogenetically adjusted datasets that are corrected with the same $N$ matrix used within pFDA (see Materials and methods), at the optimal values of Pagel's $\lambda$ found by Fabbri *et al.* [18]. Hopkins $p$ values apply to the null hypothesis that the dataset is indistinguishable from a uniform random distribution. None of the test results indicate significant clustering in any of the data sets. Abbreviations: FPM, Fernandez Pierna and Massart variant [35]; LJ, Lawson and Jurs variant [36].

In each case, and for each variation of the Hopkins statistic test, we find that we cannot reject the null hypothesis. The datasets are thus *statistically indistinguishable from a uniform random distribution* in the (log_10_(*MD*), *Cg*) space under the various Hopkins statistic tests. This is true both for the original, untransformed datasets as well as those that have been phylogenetically corrected using the same optimal values of Pagel’s λ found by Fabbri *et al.* This result is visualized in Fig 18, which shows as one example a plot of *F0D0* from ds1 compared to a uniformly random distribution that has been clipped to the same convex hull.

**Fig 18.**
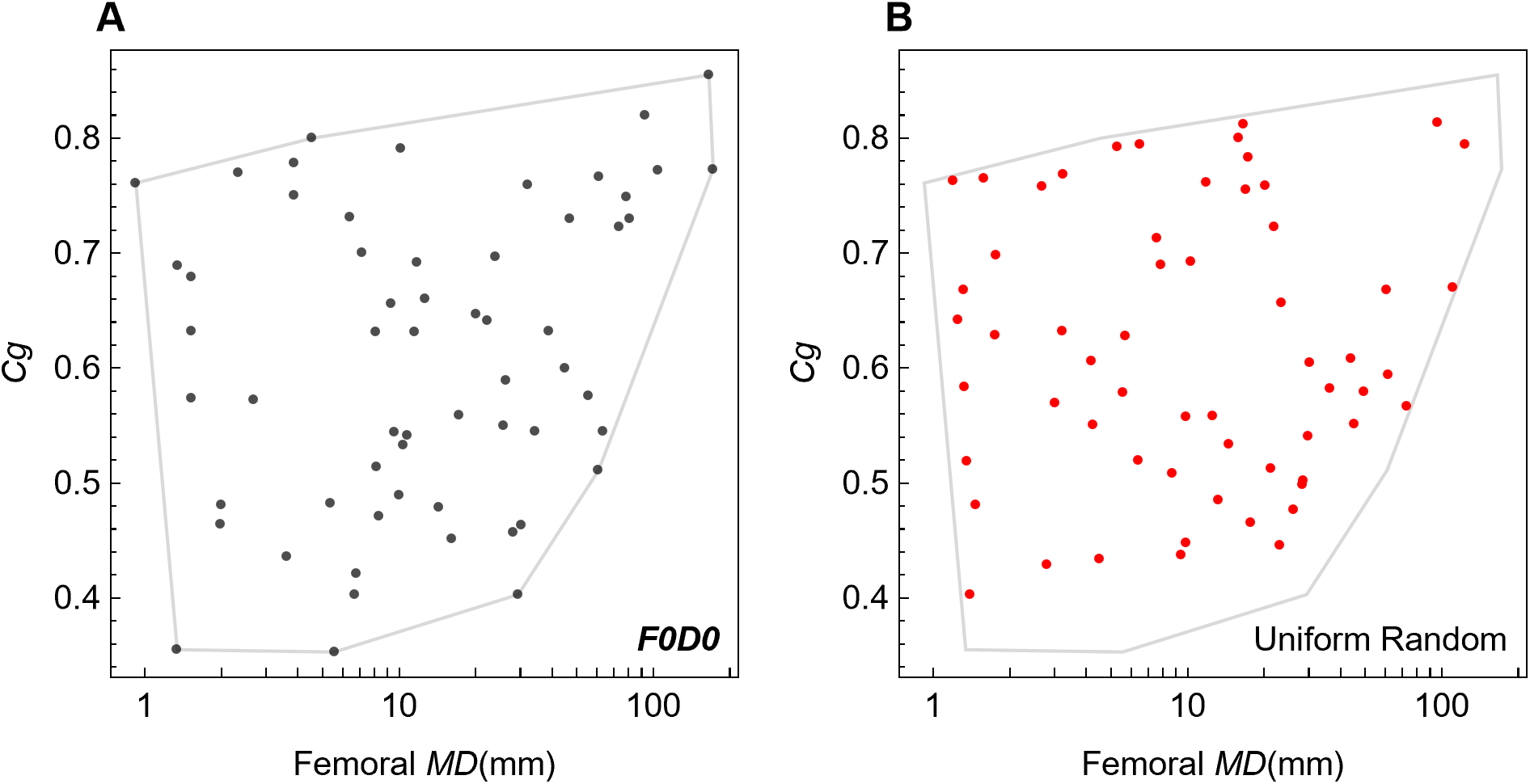
Comparison of real data from Fabbri *et al.* [18] to uniform random points. (A) Data points for the terrestrial (*F0D0*) group (black dots) in the femoral dataset of Fabbri *et al.* are plotted along with (B) uniform random points (red dots), both clipped to the convex hull enclosing the *F0D0* data. The apparent absence of any non-random concentration or clustering of the *F0D0* data is confirmed by statistical tests (Table 12).

For pFDA to be valid, both class subsets being compared must have a multivariate normal distribution and therefore the clustering that produces the normal distribution. But pFDA also requires that data not have more than one cluster because a bimodal or multi-modal distribution would also not be suitable. Our finding that the datasets appear to be uniformly random with no detectable clustering puts an important constraint on using these datasets for pFDA.

This result is consistent with the uniform distribution being a best or near-best fit for some the pFDA discriminants class subsets. The relatively strong performance of mixture distributions in Fig 16 and Table 10 suggests that, for some datasets, the distributions might be bimodal. The Hopkins-statistic results suggest (but do not prove) that apparent bimodal behavior in the discriminants may be an artifact of the low data count.

A uniformly random distribution of data points may seem strange, but biologically this corresponds to the points in (log_10_(*MD*), *Cg*) space being equally likely, at least within some range of values in each parameter. The effect might be accidentally enhanced if the goal of the creating the datasets included some notion of sampling a diversity of values in both *MD* and *Cg*. Such a goal could bias selection of taxa for the dataset toward greater spread and less clustering.

Phylogenetic bias removal might also come into play; if a cluster of values in (log_10_(*MD*), *Cg*) space is due to closely related taxa, then they may be deemphasized by the phylogenetic correction. These are general observations; whether either or both of these factors apply to the datasets under examination is unknown. A more likely factor is the effect of low *n*. Datasets that use 49 to 62 points across many clades may simply be too small to show evidence of clustering.

## Discussion

Using a bone-compactness index (*Cg*) assessed from thigh bone (femur) and trunk rib, Fabbri *et al.* [18] published two novel claims: first, that spinosaurids evolved initially as “subaqueous foragers,” a lifestyle characterization defined on habitual diving in pursuit of underwater food resources; and second that the spinosaurids *Baryonyx* and *Spinosaurus* maintained this lifestyle whereas a third, *Suchomimus*, reverted to a terrestrial, nondiving lifestyle despite its close phylogenetic and morphological affinity to *Baryonyx*.

We tested these hypotheses by examining how bone compactness, as captured in the single metric *Cg*, was sampled and measured, how lifestyles were defined and categorized, and on what basis taxa were included or excluded from datasets prior to statistical analysis. We also examined the assumptions and statistical properties of the relatively new method they utilized to make both of their principal claims, an adapted statistical procedure called phylogenetic flexible discriminant analysis (pFDA).

Our examination reveals irregularities in the composition of the dataset. The use of pFDA to infer properties of dinosaurs rests on a fundamental assumption that extinct and extant taxa have comparable values of bone compactness (*Cg*). Yet some of the datasets assembled and analyzed by Fabbri *et al.* actually demonstrate the opposite: they show a large and statistically significant bias in *Cg* values for extent taxa compared to extant taxa.

Other aspects of femoral and rib datasets raise serious questions. In some datasets, either extinct or extant taxa predominate, and some taxa are either included or excluded on questionable grounds. Are there not sufficient extant taxa available for training datasets—or at least enough to balance extinct taxa? Why are flying birds and tiny shrews and voles relevant points of comparison for flightless spinosaurids that had body masses a million times greater? Why are *Nothosaurus* and its close relatives heavily represented, while other relevant groups are ignored altogether?

The datasets include taxa scored for two variables, body size (log_10_(*MD*)) and bone compactness (*Cg*), despite the authors’ own PGLS regression analysis showing that *Cg* alone is approximately 50 times more powerful in explaining the habitual diving (*D2*) categorical variable. Taxa with categorical variables found to have little to no statistical correlation with *D2* (*i.e., F1, F2, D1*) are retained in the analysis for unknown reasons. Fabbri *et al.* created two derivative datasets (ds3, ds4) by culling so-called “graviportal” and “pelagic” taxa, claiming clear anatomical signals as the criteria. These exclusions are questionable, especially since Fabbri *et al.* did not follow their own stated anatomical criteria.

At a more basic level, the study of bone compactness in femora and trunk ribs is motivated by the idea that denser bones can act as ballast or otherwise assist diving. This is an established correlation for some extant terrestrial groups. However, there is an equally well-established correlation between increased *Cg* and body size, which the *Fabbri et al.* analysis fails to explore as an alternative hypothesis or confounding factor.

Little is known about the effects of *Cg* on femora or ribs of taxa with large body size that have substantial vertebral pneumaticity, such as the spinosaurids. Given that this pneumaticity demonstrably has a much larger effect on buoyancy than could be counteracted by increased *Cg* in limb bones or ribs, it is unclear that *Cg* increases observed in other clades are relevant.

We document a strong subjective component to assessing *Cg*, particularly from CT scans. Despite diligent efforts, we could not replicate bone-density measurements reported by Fabbri *et al.* as reasonable or without less dense alternatives for Fabbri *et al.*’s singular spinosaurid specimens. Instead, we found measurements skewed toward increased or decreased bone density. Determining *Cg* in fossilized long bones requires investigators to assess what is bone versus matrix to be masked, what cracks or missing fragments should be digitally removed or “restored,” and what thresholding intensity to use in transforming CT scans to binary images. None of these decisions are governed by specific protocols.

Although *Cg* plays a critical role in the analysis of Fabbri *et al*., often on the basis of single measurements per taxon, experts’ understanding of its variability remains rudimentary. We have found no studies on extinct or extant tetrapods that systematically compare *Cg* from multiple specimens of the same taxon, multiple bones within an individual, or even multiple places along the shaft of a bone. Sporadic examples collected from the literature support median variation of 18.6% among different individuals of the same taxon, which is about 33% of total variation in *Cg* in datasets of Fabbri *et al.* Such high intraspecific variation and lack of large-*n* benchmark studies suggest that the suitability of *Cg* for classification is far from a foregone conclusion.

We also examined the prerequisites and limitations of pFDA for this kind of classification analysis. We find that no previous study has examined the effect of finite sample size on accuracy of classification or on the predicted posterior probability of class membership. Using bootstrap trials, we constructed confidence intervals on the relevant statistical metrics. We find that the lower bound of the 95% confidence interval suggests that for most of the datasets, pFDA has an error rate of 20% to 33% in classifying its own training data. Random guesses, by comparison, would generate a 50% error rate. What scientific conclusions can be drawn from a method and dataset that is only marginally better than random chance when applied to known cases?

The posterior probability of membership in the *F0D2* class (nonflying divers) is similarly greatly weakened when taking the 95% confidence interval into account. Our sensitivity analysis shows that small variations in *Cg*, either pro-forma or from our attempted replication, also have a strong impact.

Finally, we attempted to verify that the datasets meet the distribution assumptions of pFDA by examining the classification discriminant values. For pFDA to be valid, the two datasets being compared (*e.g.*, *F0D0*/*F0D2*; terrestrial/nonflying divers) must both have normal distributions, different means, and the same variance. We find no statistical support that these conditions are met by any of the datasets. To the contrary, common clustering tests show that the data points in key datasets are statistically indistinguishable from a uniform random distribution.

In order to have any power of inference, statistical analysis must be applied to suitable data and datasets. Is *Cg* or any other metric for bone compactness free enough from measurement imprecision and biological variation to represent a species with a single data point? Can one plausibly infer “foraging” from a dataset that catalogs propensity for “diving”? Is the foraging of grazing herbivores biomechanically similar enough to active predation that it is a relevant point of comparison? This critique across many levels aims to improve future use of pFDA and other quantitative statistical methods in paleontology, because sound datasets and statistical analysis can generate inferences that go beyond structural or functional hypotheses based on select taxa.

## Acknowledgments

We thank Wayt Gibbs for editorial assistance, Lauren Conroy for assistance with several figures, Jordan Mallon for assistance in obtaining CT scans of specimens in his care, and Cem Ozen for programming assistance.

## Supporting information

S1 Table, S1–S9 Fig, S1 Appendix, and Equations (4)–(7) may be found in the Supporting information document.

