## Supporting information for "Diving dinosaurs? Caveats on the use of bone compactness and pFDA for inferring lifestyle"

#### Contents

S1 Table
S1–S5 Fig
S1 Appendix (includes S6–S9 Fig, Equations (4)–(7))

**S1 Table. Settings for computed-tomographic scans of each of the specimens described.**

Scans conducted at UChicago Hospital medical imaging facility by Nicholas Gruszauskas and David Klein on a Philips Brilliance iCT 256-slice multi-detector CT scanner

| Specimen | Pixel spacing (mm) | Spacing between slices (mm) | Exposure | Exposure time (ms) | X-ray tube current (uA) | KVP (kV) | Convolution kernel | Projections |
|---|---|---|---|---|---|---|---|---|
| MNBH GAD500 dorsal vert. 2 | 0.5104 | 0.4 | 99 | 1913 | 52 | 120 | YD | 2038 |
| FSAC-KK 11888 dorsal vert. 6 | 0.9766 | 0.4 | 39 | 1347 | 29 | 120 | YC | 2613 |
| FSAC-KK 11888 dorsal vert. 8 | 0.6510 | 0.335 | 175 | 1309 | 134 | 120 | YC | 2734 |
| MNBH GAD500 dorsal vert. 12 | 0.5234 | 0.4 | 99 | 1913 | 52 | 120 | YD | 2247 |
| MNBH GAD500 sacral vert. 2 centrum | 0.6510 | 0.335 | 175 | 1309 | 134 | 120 | YC | 2733 |
| FSAC-KK 11888 sacral vert. 3 centrum | 0.4688 | 0.4 | 99 | 1913 | 52 | 120 | YD | 2107 |
| MNBH GAD500 left femur | 0.6250 | 0.335 | 200 | 1309 | 153 | 140 | YC | 3397 |
| FSAC-KK 11888 left femur | 0.4518 | 0.4 | 99 | 1913 | 52 | 120 | YD | 2376 |
| UCRC PV8 right phalanx I-1 | 0.2747 | 0.335 | 175 | 1309 | 134 | 120 | YC | 1492 |

Scans conducted at the Transportation Safety Board of Canada Engineering Laboratory by Vincent Bolduc and Jordan Mallon on a North Star Imaging X-view X500 CT scanner

| Specimen | Voxel Size (mm) | Focal spot size (um) | Frame Averaging | Exposure (ms) | Current (uA) | Voltage (kV) | Filter | Projections |
|---|---|---|---|---|---|---|---|---|
| CMN 50382 femur | 0.0911 | 11 | 4 | 250 | 60 | 190 | 1 Cu | 1440 |
| CMN 41869 partial right femur | 0.0911 | 0 | 2 | 250 | 61 | 190 | 1 Cu | 3003 |

Scans used from previously published papers

| | |
|---|---|
| BSPG-2006-I-54 ant. dorsal vert | https://figshare.com/articles/dataset/DICOM_BSPG_2006_I_54/1471654/1 |
| NHMUK R 9951 right femur | https://www.morphosource.org/concern/parent/000406876/media/000406878 |

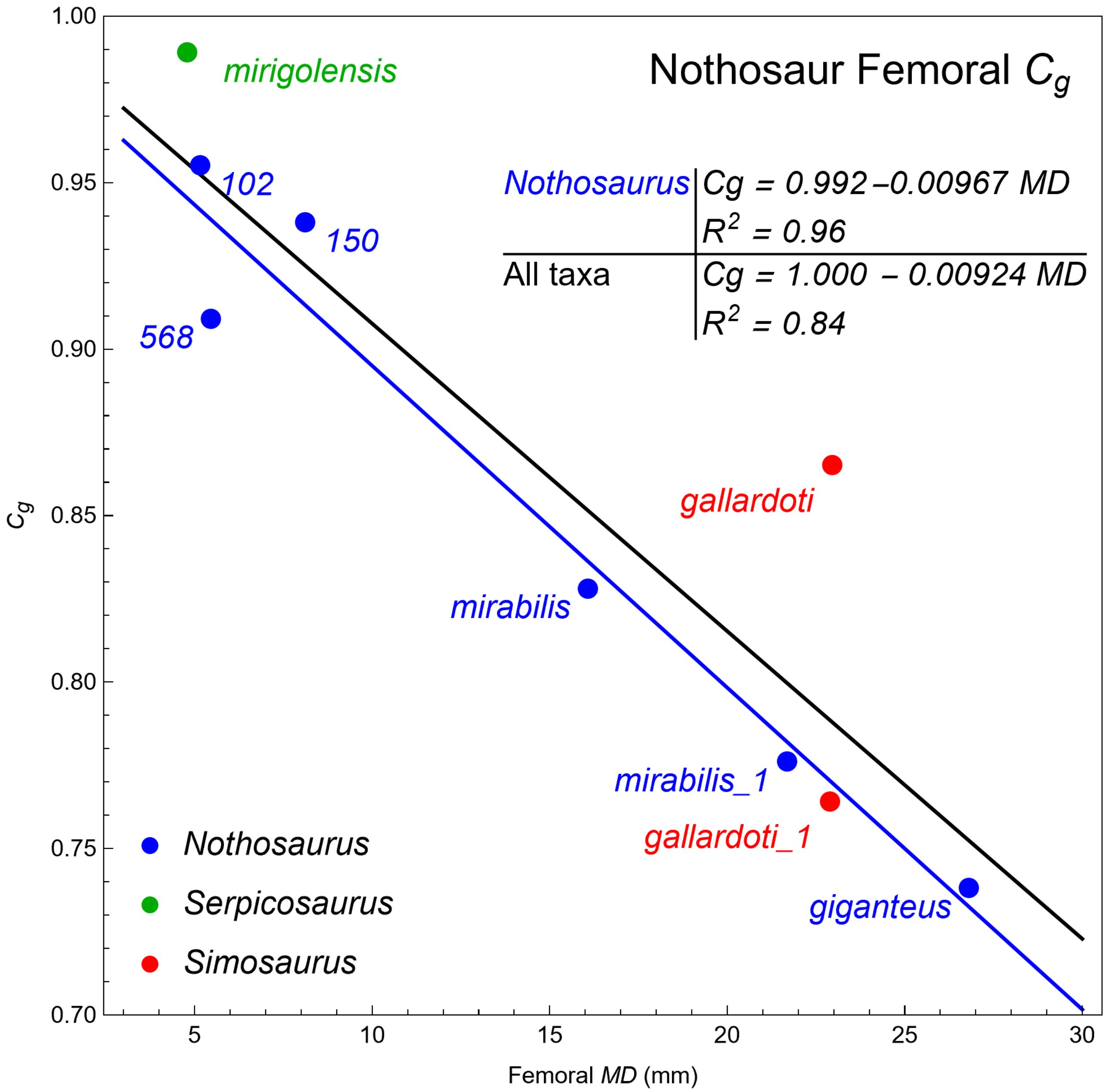


**S1 Fig. Correlation between global bone compactness (*Cg*) and femoral maximum diameter (*MD*) in Sauropterygia.** Fabbri *et al.* [1] include data from six specimens of *Nothosaurus*, two of the related nothosaur *Simosaurus,* and one related pachypleurosaur, *Serpicosaurus*. Each point is labeled with the identifier used in the Fabbri *et al.* datasets. A strong inverse correlation is shown between global bone compactness (*Cg*) and femoral *MD*, which is commonly used as a proxy for body size. The blue regression line only includes data points for *Nothosaurus,* the black regression line includes all taxa in the plot. Regression parameters are shown in the inset table. The coefficient of determination is extremely high ($R^2 = 0.96$) for *Nothosaurus* alone but still very high ($R^2 = 0.84$) for these sauropterygia taxa pooled together. The source of this strong trend is unknown to us; it could be a real biological effect, or a data artifact, or some combination thereof. Note that if extrapolated these trends would have *Cg* = 0 at *MD* = 103 mm for *Nothosaurus* and *MD* = 108 mm for all taxa, which is biologically impossible.

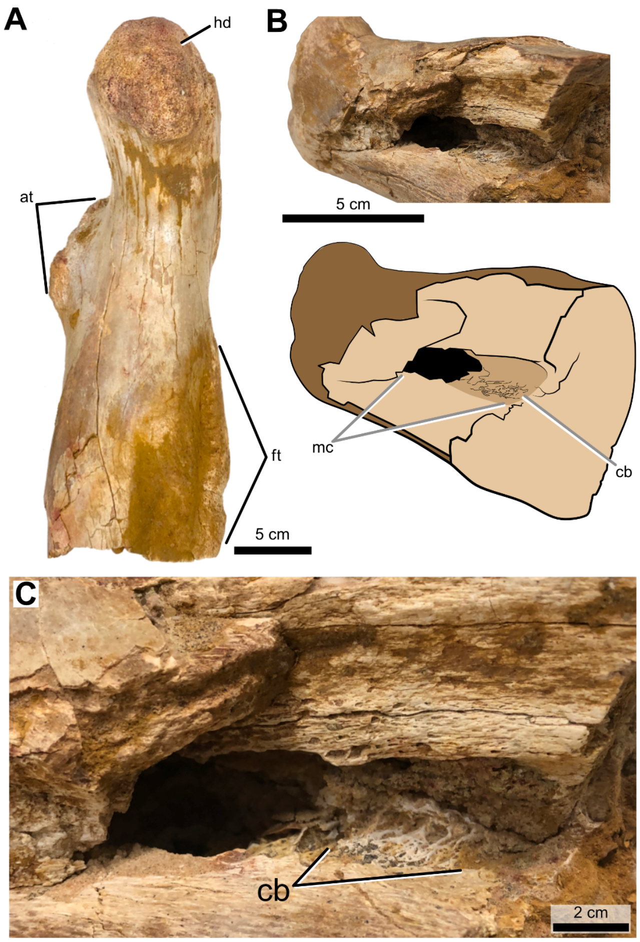


**S2 Fig. Spinosaurus aegyptiacus subadult femur retaining medullary cavity (CMN 41869).** (A) Proximal half of the right femur in medial view. (B) Medullary cavity in ventrolateral view. (C) Bone lining the medullary cavity. Abbreviations: at, anterior trochanter; cb, cancellous bone; ft, fourth trochanter; hd, head; mc, medullary cavity.

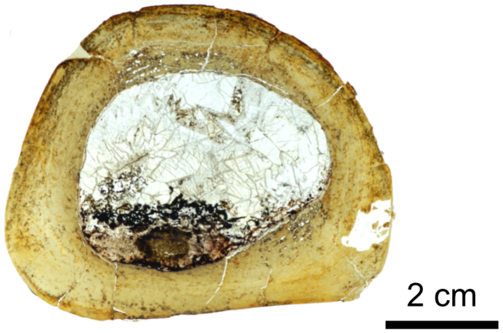


**S3 Fig. *Suchomimus tenerensis* juvenile femoral mid shaft thin section.** Thin section of the midshaft of a right femur of a juvenile individual (femur length 55.3 cm; MNBH GAD72).

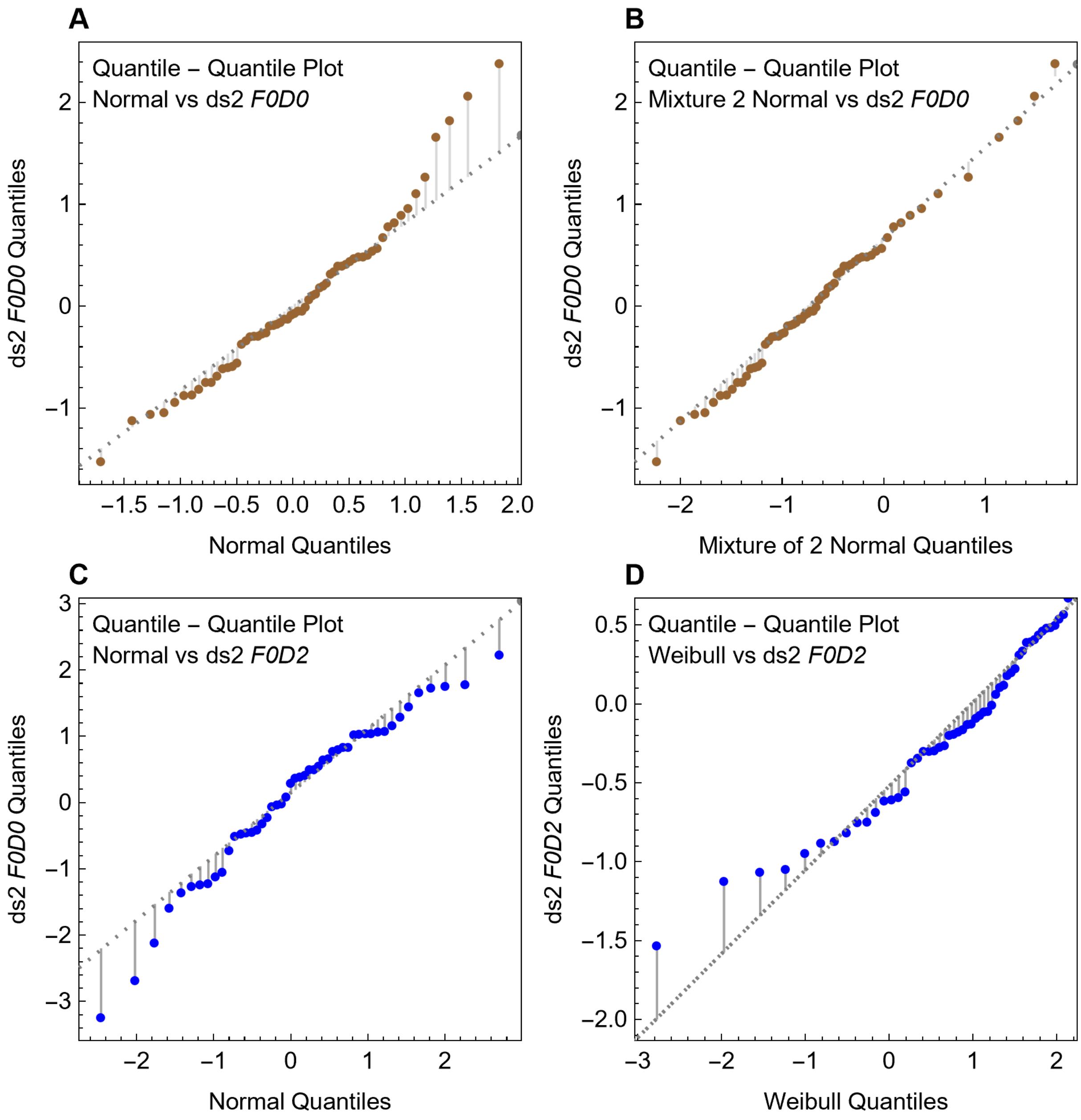


**S4 Fig. Quantile-Quantile plots of pFDA discriminants from dataset ds2 subsets *F0D0* and *F0D2*.** In these panels, the quantiles of the discriminant distributions versus those of a normal or uniform distribution (heavy black points) can be compared to plots of the normal or uniform distribution with itself (thin dotted lines). Comparable figures for ds1 are presented in Fig 17.

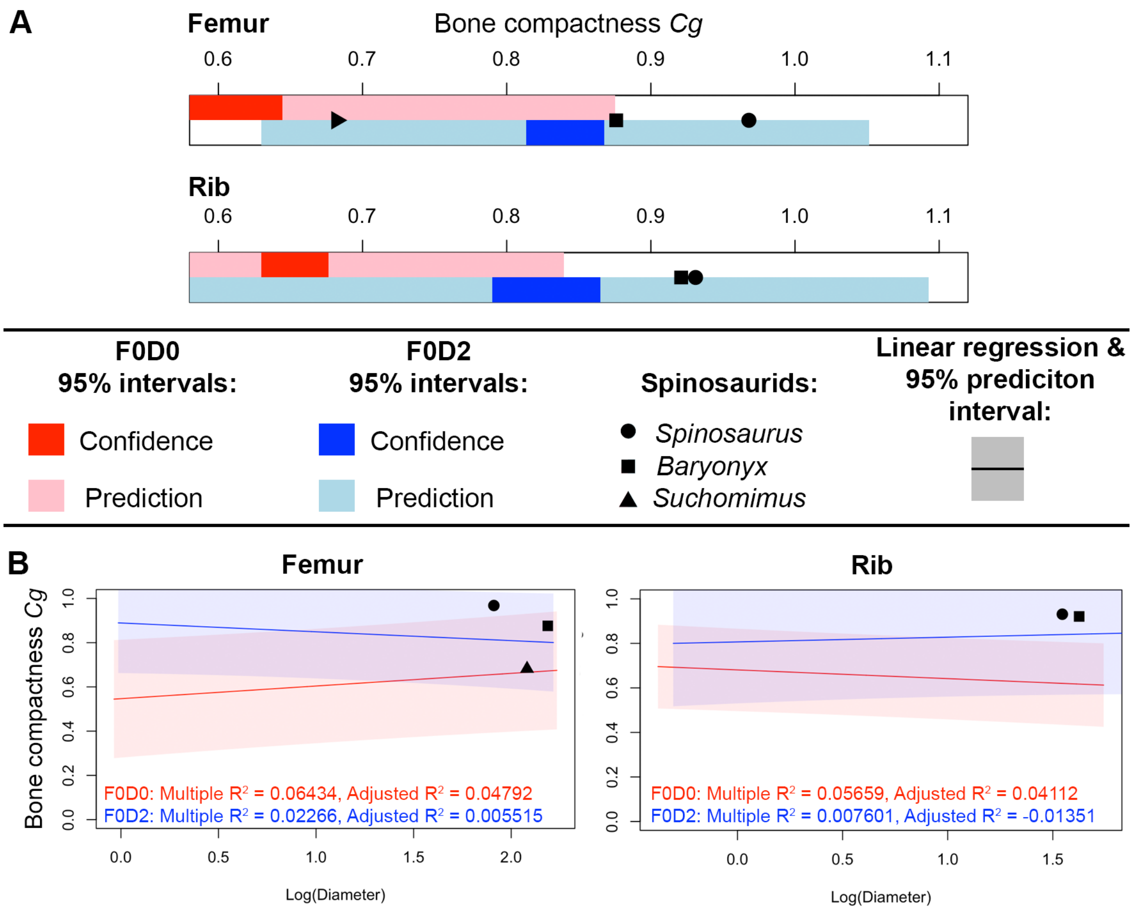


**S5 Fig. One-dimensional and two-dimensional effect size statistics on the Fabbri et al. and corrected training datasets, compared to original and remeasured spinosaurid values.** (A) The plots use bars to show the 95% confidence interval and 95% single-prediction intervals for the mean value of *Cg* in the femoral and rib training sets. Within each training set, the intervals for the *F0D0* group are shown as red (95% CI) and pink (95% prediction) bars; the intervals for *F0D2* are shown in blue and cyan, respectively. The values for spinosaurid taxa used in Fabbri *et al*. [1] are marked with solid black markers. The confidence and prediction intervals for the mean provide a simple one-dimensional view of the overlap in distributions between the *F0D0* and *F0D2* groups. In the femoral dataset, the 95% confidence interval of the mean of *F0D2* lies entirely within the prediction interval of *F0D0*, showing that even the mean *Cg* in *F0D2* would be plausible as a member of *F0D0*. In the rib dataset, the mean 95% CI for *F0D2* is mostly within the prediction interval for *F0D0*. The 95% CI for the mean of *F0D0* overlaps with the prediction interval of *F0D2* for femoral data and falls entirely within the interval for rib data. In each case we see that an average value of *Cg* distribution of one gro up (say, *F0D2* divers) is plausible as a member of the opposite group (the *F0D0* non-divers) and vice versa. The overlap in *Cg* for the groups occurs not only at the edge cases of a group but also extends to group average. (B) Linear regressions (performed without phylogenetic bias adjustment) of (*Cg,* Log(10, *MD*)) are plotted with their with 95% prediction interval for the *F0D0* and *F0D2* groups of femoral and rib datasets. Outputs for $R^2$ from the lm() function in R are reported. The two-dimensional intervals show that the overlap evident in the *Cg* plots of (A) is also present when diameter is considered. The regression results show that these two-dimensional regressions have extremely weak correlation and have somewhat minor impact on our interpretations, although they often produce *F0D0* 95% prediction intervals even closer to *Spinosaurus* values in the bivariate space. The weak correlations support the conclusion by both Fabbri *et al.* and ourselves that including bone diameter likely does not improve the predictive ability of the model.

---

### S1 Appendix

#### 1. The ecological fallacy

A common error in statistical inference known as the ecological fallacy occurs when inference is made about an individual based on aggregate data for a group [2]. In a large dataset for human stature, for example, the sex of an individual cannot be ascertained with any certainty from body mass alone, even though stature is known to differ in males and females [3]. Although the average male is heavier than the average female, overlap in the distributions eliminates the possibility of using the group-wide average or other properties derived from the aggregate (*e.g.*, equiprobable mass) for identifying the proper classification of an outside individual (S6 Fig). This approach to classification would work with high accuracy only if the distributions had little overlap.

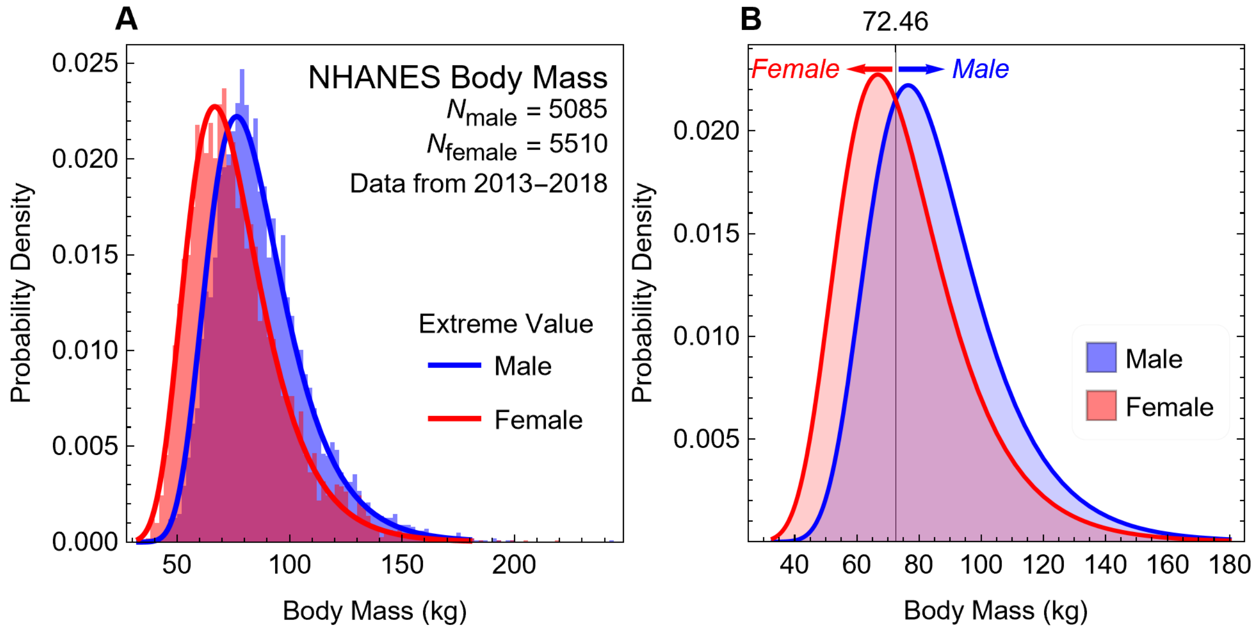


**S6 Fig. The 'ecological fallacy' illustrated with data on human adult body mass.** (A) Histogram of body mass distributions with fits to the extreme value distribution. (B) Probability curve for classification of sex by body mass.

---

To determine the accuracy of classification, one must fit a smooth distribution to the histogram and find the body mass that is equally probable in each distribution. The histograms are well modeled by the extreme value distribution, and the equiprobable point occurs as a mass of 72.46 kg (S6 Fig). This probability-based method will classify an individual specimen of unknown group membership into the groups male or female, depending on whether they are above or below the equiprobable point of 72.46 kg. Overlap in the distributions causes errors about 38% of the time: half of the males below 72.46 kg and females above that value are misclassified, assuming there is an equal probability of the unknown subject being male or female. A random guess of the group will be in error 50% of the time, so using body mass to classify in this manner is only a ~12% improvement.

The failure to classify an individual, by comparison, arises when using group-wide aggregate measures, such as the distribution. Statistics provide many analytical tools for comparing a sample group to another group to assess whether they were drawn from the same underlying population, such as Student's t-test, Welch's t-test, the Mann-Whitney test, and the *F*- and Z-tests. When the sample size *n* is small, these methods provide low statistical power. Classifying a single human—or in the case of Fabbri *et al.* [1], classifying *Spinosaurus*—is an $n = 1$ example, and these statistical tools simply do not work at such tiny sample sizes. The *ad hoc* method of Fabbri *et al.* has no support in the statistical literature.

In the large human study referenced above, group composition was carefully chosen to accurately model population-wide adult human body mass [3]. Data points potentially contaminated by confounding contributions to body mass were eliminated (*e.g.*, by removing pregnant women). In contrast, the groups identified by Fabbri *et al.* have not been controlled for confounding factors such as body mass, burrowing, and other factors known to contribute to bone compactness, and only two categorical variables (flying and diving) are considered, thus overly constraining classification to only two possibilities ("subaqueous forager" versus not).

The analysis of Fabbri *et al.* is more complex that the human example because bone density data was analyzed with PGLS-based linear regression models rather than simple distributions in order to reduce potentially confounding effects of phylogenetic signal. This analytical method does not avoid the ecological fallacy, however, because the comparison group is still $n = 1$ for *Spinosaurus*. Even when including all spinosaurids, a group of $n = 3$, statistical power remains weak. Grouping the spinosaurids also assumes incorrectly that the three species had equal degrees of secondary aquatic function.

Another difference between the human body mass example and the pFDA method of Fabbri *et al.* is that pFDA starts with a two-variable dataset (*Cg,* Log(10, *MD*)). However the FDA aspect of the method dimensionally reduces this to a single variable comparison between discriminant values, shown in Fig 15. The high degree of overlap shown in Fig 15 is an example of the ecological fallacy that is entirely analogous to the group overlap shown here in S6 Fig.

#### 2. ROC curves and whether $P_2 > 0.5$ is the best threshold

The receiver operating characteristic (ROC) curve is a longstanding graphical means of characterizing the performance of a binary (*i.e.*, two-group) classifier as the classification threshold is varied from $0 \leq P_{\text{thres}} \leq 1.0$. These curves originated with the study of radar detection of objects but have since found extensive application in both statistical evaluation of medical diagnostics [4,5] and machine learning classifiers [6–8].

S7 Fig plots ROC curves for both femoral (ds1) and rib (ds2) data in the basic datasets. The curves are obtained by using $k = 10$ cross validation, which yields a $P_2$ value for each member of the training set, obtained when that point and others are "held out" from the analysis and treated as a test point.

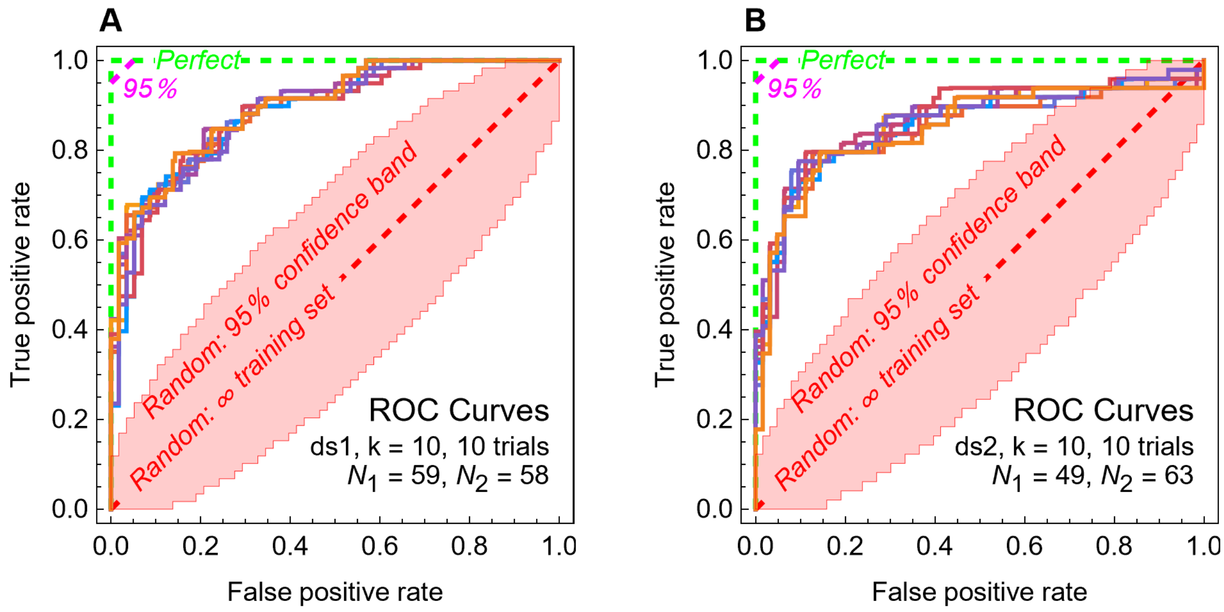


**S7 Fig. ROC curves for posterior probability threshold.** ROC curves plot the performance of a binary (two-class) classifier. (A) The basic ROC curve plots the false-positive rate *FPR* on the *x*-axis and true-positive rate *TPR* from Equation (5) as the classification threshold $P_{\text{thres}}$ that defines classification (*i.e.,* $P_2 > P_{\text{thres}}$ yields classification as $D = 2$) is varied from $0 \le P_{\text{thres}} \le 1$. Each colored curve is from a different trial of $k = 10$-fold cross-validation, repeated for 10 trials. The dashed red line is the performance of a random classifier, and the dashed green line is the performance of a hypothetical perfect classifier. As $P_{\text{thres}} \to 1$, both femoral and rib curves converge toward the random classifier. (B) The same ROC curves for rib data. Note that the rib classification is generally less successful since it is further away from the perfect classifier line.

---

The classic ROC curves show the tradeoff between the true-positive rate *TPR* and the false-negative rate *FPR* calculated from Equation (5) below, as $P_{\text{thres}}$ is varied. A hypothetical ideal binary classifier would follow the green dashed line in S7A Fig. A hypothetical classifier that would be good enough to meet a threshold of $\alpha < 0.05$, appropriate for a 95% confidence of correct predictions, will reach deep into the top left corner and must hit or exceed the dashed magenta line. At the other end of performance, a random binary classifier with an infinite training set would follow the dashed red line. The red shaded area shows the 95% confidence band of a random classifier that predicts class by a process random draw with probability $P_{\text{thres}}$.

The colored solid lines in S7 Fig plot the ROC curves from 10 trials of $k = 10$ cross-validation. The rib classifier has lower classification performance because it is farther from the corner that defines the perfect classifier. S7 Fig shows that the specific performance of the classifier depends on the classifier threshold in a complicated way—the ROC curves vary by trial and are not smooth. The ROC curve has $P_{\text{thres}}$ as an implicit parameter, but S7 Fig explicitly shows how three of the performance metrics of Equation (5) vary with $P_{\text{thres}}$.

When $P_{\text{thres}} = 0$, the random classifier will assign class $D = 0$ to each data point. It will misclassify all of the $D = 2$ points but will correctly classify the $D = 0$ points. When $P_{\text{thres}} = 1$, the opposite occurs. Therefore the confidence band for accuracy $A$ (S8A Fig) is tilted compared to the band for Matthews correlation coefficient $MCC$ (S8B Fig). Note that this dataset has different numbers of points in each class.

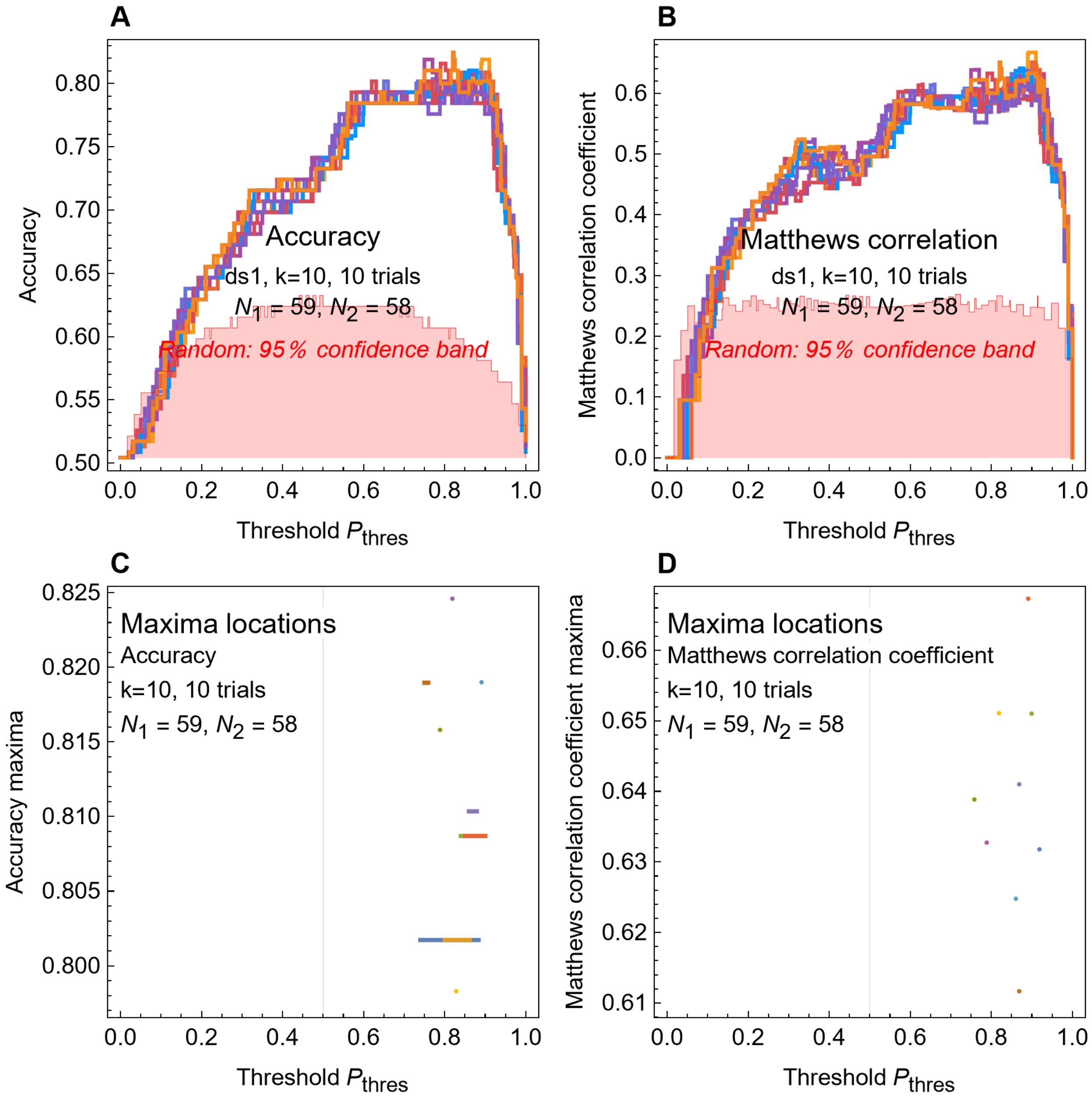

**S8 Fig. Classifier metrics as a function of posterior probability threshold for femoral classifiers.** (A,B) Three performance metrics $A$, $B$, and $MCC$ (Equation (5)) are plotted as a function of the classification threshold $P_{\text{thres}}$. Each colored curve represents the $P_2$ values from one of 10 different trials of $k = 10$-fold cross-validation of the femoral dataset. (C,D) The maximum accuracy values attained by each of the three metrics are plotted for a value or linear range of $P_{\text{thres}}$ values at which the maxima occur. In general, it is not possible to achieve the maximum accuracy or other classification metric by choosing a fixed value of $P_{\text{thres}}$ because optimization depends both on the trial curve and also on the metric used.

---

In S8C Fig and S8D Fig, the horizontal extents of the lines indicate the range of values $P_{\text{thres}}$ for which each curve achieves its maximum value. The colors of the lines correspond to the matching curves in the upper panels. S8 Fig illustrates the problem inherent in the remark by Fabbri *et al.* that they might need higher values of $P_{\text{thres}}$. Although changing $P_{\text{thres}}$ might help with the classification of an individual test data point, it would also alter the overall classification performance. A higher value of $P_{\text{thres}}$ produces fewer false positives (predicting class 2 for a datum that is actually class 1), but more false negatives (predicting a valid class 2 as class 1). Each of the different cross-validation trial curves achieves its maximum at different values of $P_{\text{thres}}$. S9 Fig shows the same kind of curves for pFDA classifiers built from rib training data. The curves are very different from those in S8 Fig.

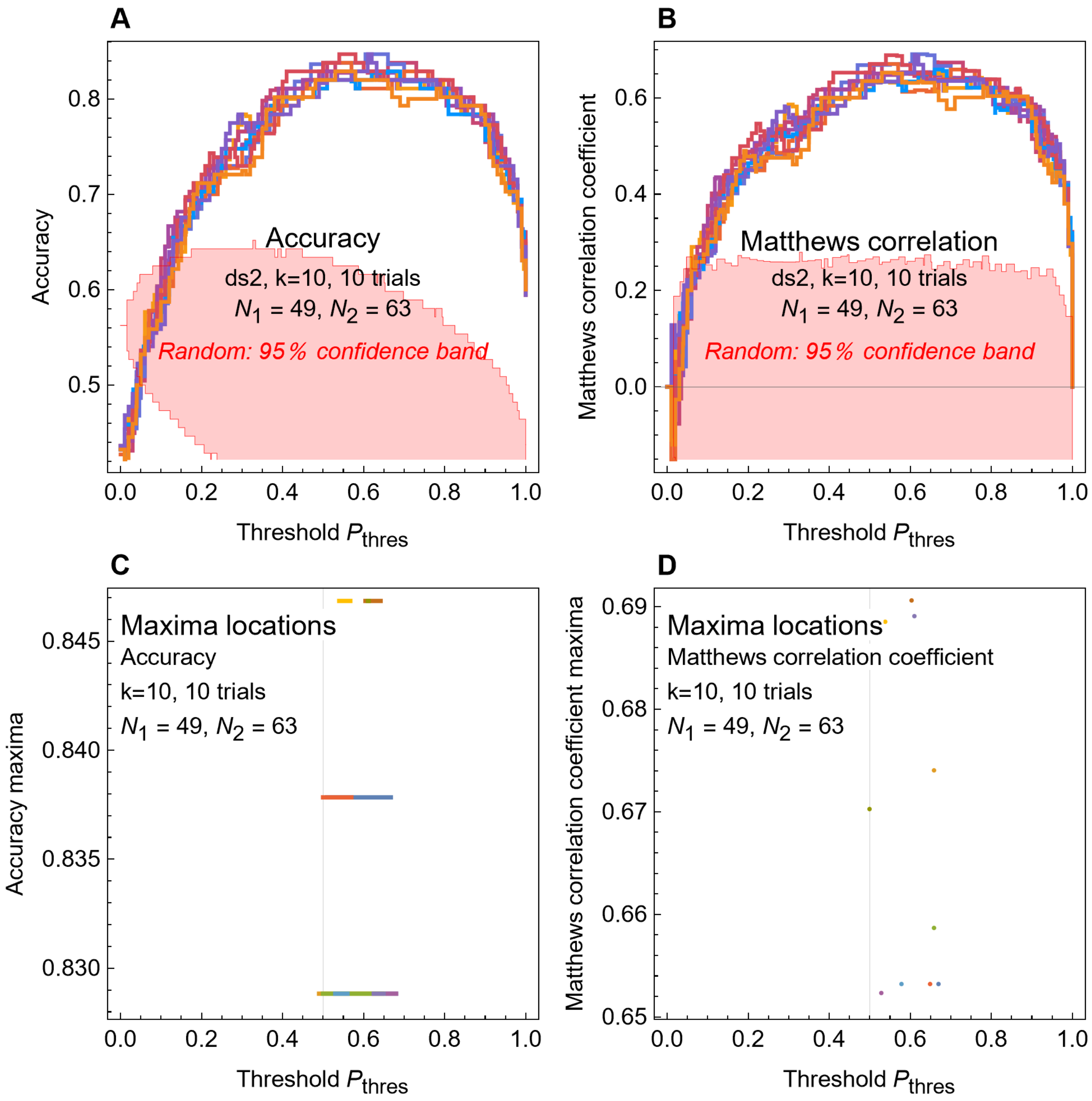


**S9 Fig. Classifier metrics as a function of posterior probability threshold for rib classifiers.** (A,B) The three performance metrics from Equation (5) are plotted as a function of the classification threshold $P_{thres}$. Each colored line is based on the $P_2$ values from a single trial of $k$ = 10-fold cross-validation of the femoral dataset; there are 10 such trials plotted. (C,D) Plots show the location of the maximum values attained by each of the three metrics. The maximum value may be obtained for a single point, or for a linear range of $P_{thres}$ values.

---

S8 Fig and S9 Fig show that obtaining optimum performance for the classifiers is quite complex, not a simple matter of moving from $P_{thres} = 0.5$ to $P_{thres} = 0.8$. The maxima is highly dependent on the data set and the value of $P_{thres}$. It even varies from one random $k$-fold cross-validation trial to the next.

Fabbri *et al.* suggest that it may be necessary to have $P_{\text{thres}} \geq 0.8$, but that is not true for any of the cases examined. Femoral (dataset ds1) training data classifiers generally achieve maxima for accuracy for $P_{\text{thres}} > 0.5$, but the exact value differs between trials (S8 Fig). Whereas balanced accuracy (ds1) has no maxima for $P_{\text{thres}} > 0.8$, the Matthews correlation coefficient has maxima for several curves near $P_{\text{thres}} \approx 0.9$ (S8B Fig).

The rib data classifier is almost entirely different—none of the metrics achieves a maximum for $P_{\text{thres}} > 0.6$; increasing the threshold above that value seriously reduces performance (S6 Fig). At $P_{\text{thres}} = 1$, balanced accuracy and Matthews correlation coefficient have values no better than random guessing.

The ROC curves and specific metric curves demonstrates that classifier performance is a nonlinear effect that can be counterintuitive. Increasing the value of $P_{\text{thres}}$ might seem like a way to improve classifications, but in general it is not. The results here are for $k = 10$-fold cross-validation, but qualitatively similar complexity exists for other values of $k$, as well as for LOOCV and for other performance metrics.

#### 3. *p*-values and the $p < 0.05$ threshold

There has been an ongoing debate about the role of *p*-values and the choice of $a = 0.05$ in scientific research. Discussion has focused on several problematic issues, including inability to replicate results [9] and strong incentives to find "significant" effects that lead to "*p*-hacking" and other kinds of data manipulation [10]. A common criticism among statisticians is that *p*-value, by itself, does not completely capture the nature of the statistical results. Effect size is also important; a "significant" effect that is very small may not be very important [11].

Fabbri *et al.* provide a good example of this latter issue in their PGLS regression of bone compactness versus their categorical variable *D*. They follow the normal practice of using $p < 0.05$ as their significance threshold. Yet the actual effect size they find in the regression is remarkably low, with $R^2 = 0.178$ for femoral data and $R^2 = 0.108$ for rib data. These values indicate that the correlation between *Cg* and the categorical variable *D,* even if it is statistically significant, explains just 11% to 18% of the variation across the dataset.

The debate about appropriate values of the threshold *a* has primarily been focused on *lowering* the threshold to $a = 0.01$ [12] or $a = 0.005$ [13,14]. Conversely, there are some examples where very strong effects have been published in the scientific literature at $a = 0.1$. However, we are not aware of any scientific results that are considered to be valid without qualifications based on statistical analyses that have 20% to 33% or more classification error inherent in them.

#### 4. Classification performance metrics

Counting misclassifications is not sufficient to measure how well or poorly a classifier works. In a two-class classification problem, the usual terminology is to call one case "positive" and the other "negative."

The most basic element on which performance metrics are constructed is the confusion matrix $c$, which in the case of two-class classification problem is given by

$$c = \begin{pmatrix} tp & fn \\ fp & tn \end{pmatrix}, n_{\mathrm{p}} = tp + fn, n_{\mathrm{n}} = tn + fp \tag{4}$$

where *tp* is the number of true positives—*i.e.*, data points that are part of the positive class, and which are classified to be positive, and *fn* is the number of false negatives—*i.e.*, the number of data points that are actually part of the positive group despite being classified as negative. These two counts sum to $n_{\mathrm{p}}$, the total number of points in the positive group. The other parameters are the number of false positives *fp* and true negatives *tn*, which sum to the number of points in the negative group $n_{\mathrm{n}}$. Several of the most common metrics based on the confusion matrix are shown in Equation (5).

$$\begin{gathered} A = \frac{tp + tn}{tp + fn + fp + tn} \\ B = \frac{1}{2}\left(\frac{tp}{tp + fn} + \frac{tn}{tn + fp}\right) \\ MCC = \frac{tp * tn - fp * fn}{\sqrt{(tp + fp)(tp + fn)(tn + fp)(tn + fn)}} \\ TPR = \frac{tp}{tp + fn} \\ FPR = \frac{fp}{fp + tn} \end{gathered} \tag{5}$$

The simplest of these metrics is accuracy *A*, the total number of correct classifications divided by the total number of data points. This metric has intuitive appeal, but it has many known weaknesses—for example it can understate or overstate the accuracy if the number of points in the groups or the misclassification rates vary between the two classes. Balanced accuracy *B* was developed to overcome those issues, but it too has certain weaknesses. Many other metrics exist, but arguably the best single-valued metric is the Matthews correlation coefficient *MCC* [15]. *MCC* has the value of 1 for perfect classification, 0 for classification that is no better than random, and a minimum value of −1 for a completely anticorrelated result (*i.e.*, a classifier that classifies all members of the positive group as negative and vice versa).

For the simple, highly symmetric example distribution of Equations (1) and (2) and the misclassification probability (3), we can derive a theoretical value of the accuracy:

$$A = \frac{1}{2}\left(2 - \operatorname{erfc}\left(\frac{d}{\sigma\sqrt{2}}\right)\right) \tag{6}$$

In addition to the different metrics for assessing the confusion matrix, there are also different ways to calculate it. Training-set errors—misclassifications that occur within the training set itself—may underestimate the true misclassification rate that would occur with test points that have not been processed by the algorithm in the same way that the training set has. We found misclassified points in Fabbri *et al.* that were in the training set yet were still misclassified.

#### 5. pFDA bug or misunderstanding

While working with the pFDA code of Motani and Schmitz [16], we discovered that the confusion matrix for training-set classification errors returned the classification matrix with a different layout than any other reference or source we could find. The normal confusion matrix is given by Equation (4). We found that pFDA instead returns

$$c_{pFDA} = \begin{pmatrix} tp & fp \\ fn & tn \end{pmatrix} \qquad (7)$$

This is the matrix transpose of the confusion matrix as it is usually defined. The pFDA returns the matrix in this form because that is also the convention for the mda R library, which implements FDA without the phylogenetic correction done by pFDA. The documentation for the function responsible does not specify the form of the confusion matrix but simply refers to "the confusion matrix" without elaboration [17]. The documentation includes an example that uses the famous Fisher dataset for LDA, which includes measures of Iris flower petals for different varieties of Iris. It is clear from the example that the layout of the confusion matrix obeys Equation (7) rather than Equation (4) because the confusion matrix in the format of (4) has rows that sum to the total number of points (50 in the case of the Iris dataset). The result shown on the manual page has columns that sum to 50, whereas the rows sum to 50, 49, and 51.

It is unclear whether this reflects a software bug—*i.e.,* the mda library intended it to be the normal convention but mistakenly returns the transpose—or a deliberate decision to return a non-standard confusion matrix. Either way, it is potentially a trap for the unwary user of mda and, by extension, pFDA. Using a different confusion matrix format would alter the results of classification accuracy metrics in Equation (5). We do not know whether the "correct classification rates" reported by Fabbri *et al.* were based on the confusion matrix returned by pFDA, but if so, that could account for some of the differences between our accuracy calculations and theirs.
